# Lysergic acid diethylamide reverses aging- and neurodegeneration-associated brain transcriptional programs

**DOI:** 10.64898/2026.05.26.727809

**Authors:** Aurora Savino, Carla Liaci, Ilaria Bertani, Simona Rando, Mattia Camera, Giorgio R. Merlo, Lidia Avalle, Valeria Poli, Nereo Kalebic, Francesco Iorio

**Affiliations:** Human Technopole, Computational Biology, Palazzo Italia, Viale Rita Levi Montalcini 1, 20157 Milan, Italy; Molecular Biotechnology Center, Department of Molecular Biotechnology and Health Sciences, University of Turin, via Nizza 52, Turin 10126, Italy; Department of Science and Technological Innovation, University of Piemonte Orientale, Viale Teresa Michel 11, 15121 Alessandria, Italy

**Author notes:** co-senior authors (Valeria Poli, Nereo Kalebic, Francesco Iorio).

**Keywords:** Lysergic Acid Diethylamide, Brain Aging, Neurodegeneration, Gene Expression, Transcriptome, Amyloid beta-Peptides

## Abstract

Psychedelic compounds such as lysergic acid diethylamide (LSD) are increasingly studied for their neuroplastic effects and potential relevance to brain aging and neurodegeneration. However, the molecular mechanisms linking psychedelic-induced plasticity to age-associated cognitive decline remain unclear. Brain aging and dementia are characterized by coordinated transcriptional programs that underlie synaptic dysfunction and altered neuron-glia interactions. If psychedelic-induced plasticity engages opposing molecular programs, it could counteract these conserved trajectories. In computational drug discovery, this concept has been formalized as the principle of transcriptional signature reversal, whereby compounds inducing gene expression states opposite to disease-associated programs may exert a therapeutic effect by counteracting disease-associated phenotypes.

Here, we combine cross-species transcriptomic analyses with experimental validation to test whether LSD opposes conserved signatures of brain aging and dementia. By comparing transcriptional profiles induced by chronic LSD treatment in rodents with age- and dementia-associated gene expression changes in the human prefrontal cortex, we show that LSD induces gene expression patterns strongly anti-correlated with aging and neurodegeneration programs. This reversal is specific compared to other pharmacological perturbations and is reproducible across datasets and species. Moreover, LSD counteracts amyloid-β-induced structural and molecular alterations in primary cortical neurons, linking transcriptomic opposition to functional rescue under neurodegenerative stress.

Together, our findings suggest that LSD modulates molecular and cellular pathways associated with brain aging and neurodegeneration, linking systems-level gene expression changes to structural and functional resilience in neurodegeneration-relevant contexts.

## Introduction

Psychedelic compounds are increasingly recognised as promising therapeutic agents for a range of psychiatric disorders (De Gregorio et al. 2021). Among the most extensively studied are psilocybin, lysergic acid diethylamide (LSD), 2,5-dimethoxy-4-iodoamphetamine (DOI), and dimethyltryptamine (DMT). Despite their chemical diversity, these substances share a common pharmacological feature: high-affinity binding to and activation of the serotonin 5-HT_2_A receptor, which is widely considered central to their psychoactive and therapeutic effects (Banks et al. 2021). A growing body of evidence indicates that classic psychedelics function as psychoplastogens, rapidly inducing both structural and functional neuroplasticity (Inserra et al. 2021; Banks et al. 2021; Ly et al. 2018; Calder & Hasler 2023; Aleksandrova & Phillips 2021). Consistently, these compounds promote neurogenesis both in vitro (Morales-García et al. 2017) and in vivo (Lima da Cruz et al. 2018), drive dendritic spine remodelling in pyramidal neurons (Ly et al. 2018; Yoshida et al. 2011; Shao et al. 2021; Jones et al. 2009), and upregulate the expression of neurotrophic factors such as brain-derived neurotrophic factor (BDNF) (de Vos et al. 2021).

These convergent cellular and molecular effects suggest that psychedelics engage transcriptional programs supporting synaptic maintenance and neuronal resilience: processes that are progressively compromised during brain aging and neurodegeneration. On this basis, psychedelics have been proposed as potential therapeutic agents to slow cognitive decline and mitigate the risk or progression of Alzheimer’s disease (Saeger & Olson 2022; Aday et al. 2020; Winkelman et al. 2023; Haniff et al. 2024; Kozlowska et al. 2022; Sinha et al. 2024; Androni et al. 2025). However, clinical investigation in this area remains at an early stage, with completed studies limited to Phase I trials assessing safety and tolerability (Family et al. 2020; Bouchet et al. 2024), and an ongoing pilot trial evaluating the potential of psilocybin for the treatment of early-stage dementia (ClinicalTrials.gov identifier: NCT04123314).

Recently, the first direct links between psychedelics and the hallmarks of aging have begun to emerge. Psilocin, the active metabolite of psilocybin, has been shown to extend cellular lifespan in vitro without inducing malignant transformation, consistent with a delay of, rather than an escape from, replicative senescence. In the same study, intermittent administration of psilocybin to aged mice significantly improved survival, pointing to a longevity-promoting effect even when treatment was initiated late in life (Kato et al. 2025). Together, these findings raise the possibility that psychedelics may modulate conserved molecular programs associated with aging, but whether they can counteract age-associated transcriptional changes remains largely unexplored.

Foundational work in computational drug discovery and repositioning has established that compounds inducing transcriptional states anti-correlated with disease-associated gene expression signatures may have therapeutic potential (Arloth et al. 2015; Iorio et al. 2013; Lamb et al. 2006)(Dudley et al. 2011; Kunkel et al. 2011; Dönertaş et al. 2018; Wagner et al. 2015)(Arloth et al. 2015; Iorio et al. 2013; Lamb et al. 2006) Importantly, it has also been experimentally validated, with quantitative evidence showing that compounds reversing disease-associated transcriptional programs can improve disease phenotypes in vivo, as demonstrated in a mouse model of dyslipidemia (Wagner et al. 2015).

Building on these premises, we sought to determine whether psychedelics induce transcriptional states that oppose aging-associated gene expression programs in the human cortex, thereby providing a mechanistic framework to assess their potential relevance to age-related cognitive decline. To this end, we systematically compared gene expression signatures elicited by chronic LSD treatment in the rat medial prefrontal cortex with age-associated transcriptional changes in the human prefrontal cortex, leveraging large-scale RNA-seq datasets and public transcriptomic repositories. This enabled the construction of a comprehensive transcriptomic resource that captures ageing-related gene expression changes in the human prefrontal cortex. To assess the robustness and cross-species concordance of our observations, we additionally generated and analyzed transcriptomic data from the prefrontal cortex of mice treated with LSD. We then experimentally validated key predictions using mouse cortical neuronal cultures, uncovering transcriptional and cellular signatures through which LSD opposes aging-associated programs.

## Results

### LSD-induced transcriptional changes oppose age-associated gene expression programs in the human prefrontal cortex

We analysed the largest RNA-seq dataset to date of chronic LSD-induced gene expression changes in the rat medial prefrontal cortex (mPFC) (Martin et al. 2014; Savino & Nichols 2022), with the goal of determining whether the transcriptional signatures induced by LSD could be classified as opposing aging-associated gene expression programs in the human brain.

To derive a consensus transcriptomic signature of human brain aging, we queried the Gene Expression Omnibus (GEO) database (Barrett et al. 2013) for prefrontal cortex (PFC) gene expression datasets with annotated donor age (**Tables S1-S3**). We integrated gene expression data from healthy individuals, encompassing 3,086 samples spanning ages from 2 days to 106 years (**Figure S1**). To maximize comparability, we restricted the analysis to the dorsolateral prefrontal cortex (DLPFC), which provided the largest number of samples (n = 681; **Figure S2**). Across studies, we retained 4,893 genes consistently measured in all datasets.

We then computed gene-wise Pearson correlations between expression levels and chronological age across the integrated dataset and ranked all genes according to their correlation with age. This ranked list, therefore, defined a genome-wide aging trajectory, ranging from genes upregulated to genes downregulated with age.

To test whether LSD-induced transcriptional changes oppose aging-associated programs, we performed gene set enrichment analysis (GSEA) (Subramanian et al. 2005), using the age-correlation-ranked list as background gene-list and the sets of genes up- or downregulated by chronic LSD treatment in rats (Martin et al. 2014) as query signatures (**Methods**). In this way, we assessed whether LSD-upregulated genes were enriched among genes whose expression decreases with age (thus toward the bottom of the age-correlation-ranked list), and whether LSD-downregulated genes were enriched among genes whose expression increases with age (thus toward the top of the age-correlation-ranked list), consistent with reversal of aging-associated transcriptional programs.

We found that genes upregulated by LSD were significantly enriched among those whose expression was negatively correlated with age, thus toward lower end of the age-ranked gene list (GSEA p = 1.8 × 10^−25^, normalised enrichment score (NES) = -5.81, **Figure S3**).

Consistently, genes downregulated by LSD were enriched among those positively correlated with age, located at the upper end of the ranked list downregulated genes: GSEA p = 3.6 × 10^−44^, NES = 5.54, **Figure S3**).

We next repeated the analysis across individual datasets. This strategy increased coverage by avoiding batch correction and not restricting the analysis to genes commonly measured across all datasets. It also allowed us to include data from the medial prefrontal cortex (mPFC), the brain region analysed in the LSD treatment dataset. Despite substantial inter-dataset heterogeneity, most datasets exhibited a consistent overall enrichment pattern of the LSD induced signature (**Figure 1A**, **Figure S4**), with a slightly stronger and more reproducible reversal effect for genes upregulated by LSD than for the downregulated ones. Meta-analysis of dataset-specific statistics using Fisher’s method (Fisher 1992) yielded a combined p-value < 2.2 × 10^−308^ for upregulated genes and 5.2 × 10^−100^ for downregulated genes, further supporting the potential of LSD to reverse age-associated transcriptional signatures.

**Figure 1.**
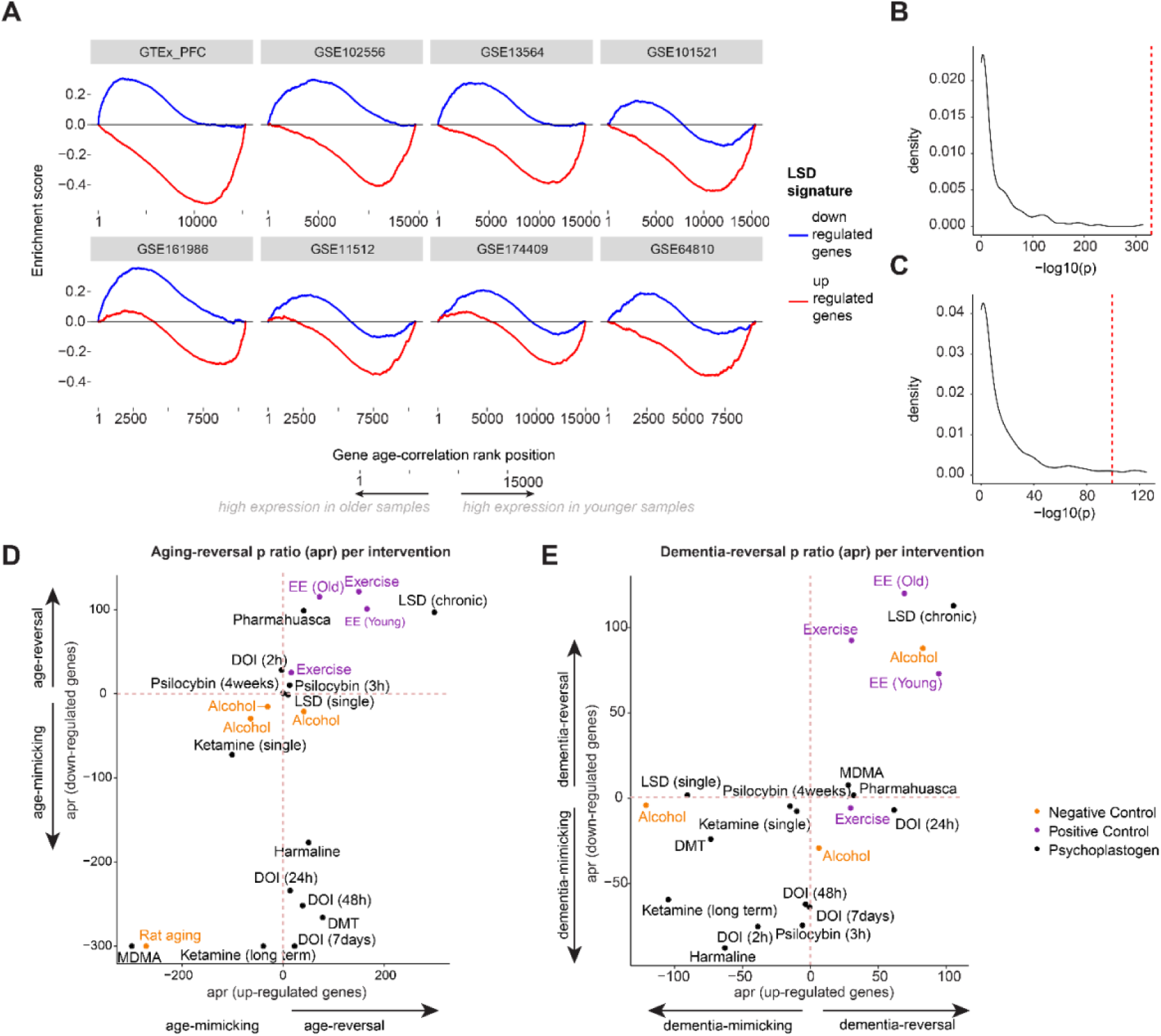
A. Gene Set Enrichment Analysis (GSEA) of genes ranked according to the correlation of their expression with age across prefrontal cortex datasets using differentially expressed genes upon chronic LSD administration (up- or down-regulated, as indicated by the different colours) as query signatures. Shown are the eight datasets exhibiting the highest absolute enrichment scores (out of fifteen total datasets, see also Supplementary Fig 4). B, C. Distribution of age-signature reversal p-values for drugs profiled in the Connectivity Map and tested in neural progenitor cells, computed using sets of drug-dependent upregulated (B) or downregulated (C) genes as query signatures, following the same GSEA-based framework summarized in panel A. The red dashed line indicates the p-value obtained for chronic LSD exposure. D. Summary of transcriptional age-signature reversal and mimicking properties of various treatments: a set of psychoplastogens in black, positive controls in purple, negative controls in orange. EE=enriched environment. For each intervention dataset (psychoplastogen, exercise, enriched environment), we computed the meta-analytic p-value for aging reversal and mimicking across human aging datasets, separately for genes up- and down-regulated upon the intervention. We then plotted the ratio between the age reversal and age mimicking p-values (-log10), indicating a higher likelihood of the intervention having an age-reversal than an age-mimicking effect. E. Same plot as D, starting from a dementia dataset instead of aging.

To assess the robustness of our findings, we evaluated whether the age-associated transcriptional patterns captured by the datasets analysed here were consistent with an independent, previously published brain aging signature (Dönertaş et al. 2018).

Specifically, we repeated a GSEA using genes reported to increase or decrease expression with aging in that study as query signatures. We observed a strong concordance across datasets, as reflected by highly significant Fisher-aggregated p-values (3.2 × 10^−227^ and 7.8 × 10^−322^ for genes with increasing/decreasing expression with age, respectively; **Figure S5**), indicating that our datasets robustly capture canonical aging-related transcriptional effects. Together, these findings indicate that chronic LSD treatment induces gene expression changes that oppose age-associated patterns, consistent with a potential reversal of aging-related transcriptional programs.

### LSD exhibits the strongest age-signature reversal effect among psychoplastogens and pharmacological perturbations

To assess the specificity of the observed age-signature reversal and to exclude the possibility that it reflects a generic property of perturbational transcriptomic profiles, we compared the age-reversal potential of LSD against a broad panel of pharmacological perturbations. Specifically, we analysed transcriptional signatures from 193 compounds profiled in neural progenitor cells within the Connectivity Map (CMap) resource (Subramanian et al. 2017), applying the same GSEA-based framework used for LSD. Under a non-specific or confounded scenario, a substantial fraction of drug-induced signatures would be expected to exhibit comparable age-reversal effects. Instead, we found that LSD ranked above all tested compounds when considering upregulated genes, and above 97% of compounds when considering downregulated genes, based on Fisher-aggregated p-values (**Figure 1BC**, **Figures S6** and **S7**, **Methods**), supporting a high degree of specificity of the LSD-associated transcriptional reversal of aging signatures.

We next asked whether age-signature reversal was specific to LSD or could also be observed for other psychedelics, entactogens, and dissociative compounds (collectively referred to as “psychoplastogens”; **Table S4**) for which rodent prefrontal cortex transcriptomic data were available. As benchmarks, we considered physical exercise and enriched environment (EE) as positive controls, ethanol exposure as a negative control, and rat aging signatures as an additional age-mimicking reference (**Table S5**).

For each condition, we identified differentially expressed genes (FDR < 0.05) and evaluated their directional enrichment along aging-associated ranked gene lists using the same GSEA-based framework applied to LSD (**Methods**). Dataset-specific statistics were aggregated using Fisher’s method, and we quantified the net directionality of each intervention as the age-reversal p-value ratio (apr), reflecting the relative strength of age-reversal versus age-mimicking signals (**Figure 1D**, **Methods**).

Chronic LSD exposure exhibited the strongest overall age-reversal effect (average apr across up- and down-regulated genes = 198), exceeding that observed for physical exercise and EE, which, as expected, also showed positive apr values (largest average apr = 136; **Figure 1D**, **Tables S6** and **S7**). In contrast, the rat aging signature displayed the most pronounced negative average apr (-286), consistent with strong age-mimicking activity. Ethanol treatment similarly produced an age-concordant transcriptional profile (largest average apr = -47).

Among the remaining psychoplastogens, only repeated pharmahuasca exposure (average apr = 70) and, to a much lesser extent, acute psilocybin treatment assessed 3 hours post-administration (average apr = 12) showed mild reversal effects compared to that of the positive controls. Most other interventions exhibited age-mimicking profiles, with MDMA displaying the strongest age-concordant transcriptional signal (average apr < -300; **Figure 1D**).

Extending the analysis to dementia-associated signatures, mostly Alzheimer’s Disease-related, but comprising also dementia with Lewy bodies, Parkinson’s disease dementia and Hungtington Disease (**Table S8**) yielded a similar pattern: chronic LSD treatment showed a robust reversal effect (average apr = 109), exceeding that observed for EE (largest average apr = 95) and physical exercise (largest average apr = 61; **Figure 1E**, **Tables S9** and **S10**).

### Acute LSD treatment in mice recapitulates the transcriptional age-reversal signature across doses

Next, we generated an independent transcriptomic dataset from mice following a single intraperitoneal administration of LSD (**Figure 2A**). We tested multiple doses (50, 100, 200, and 500 µg kg^−1^) and profiled gene expression in the PFC 90 minutes after treatment. Across all doses, DEGs (FDR < 0.05) exhibited transcriptional changes opposing age-associated signatures, for both up- and downregulated gene sets (average apr across all concentrations = 100, **Figure 2B**, **Tables S11** and **S12**). Notably, higher doses did not produce stronger age-reversal effects, with the most robust signal observed at the intermediate dose of 200 µg kg^−1^ (average apr = 265).

**Figure 2.**
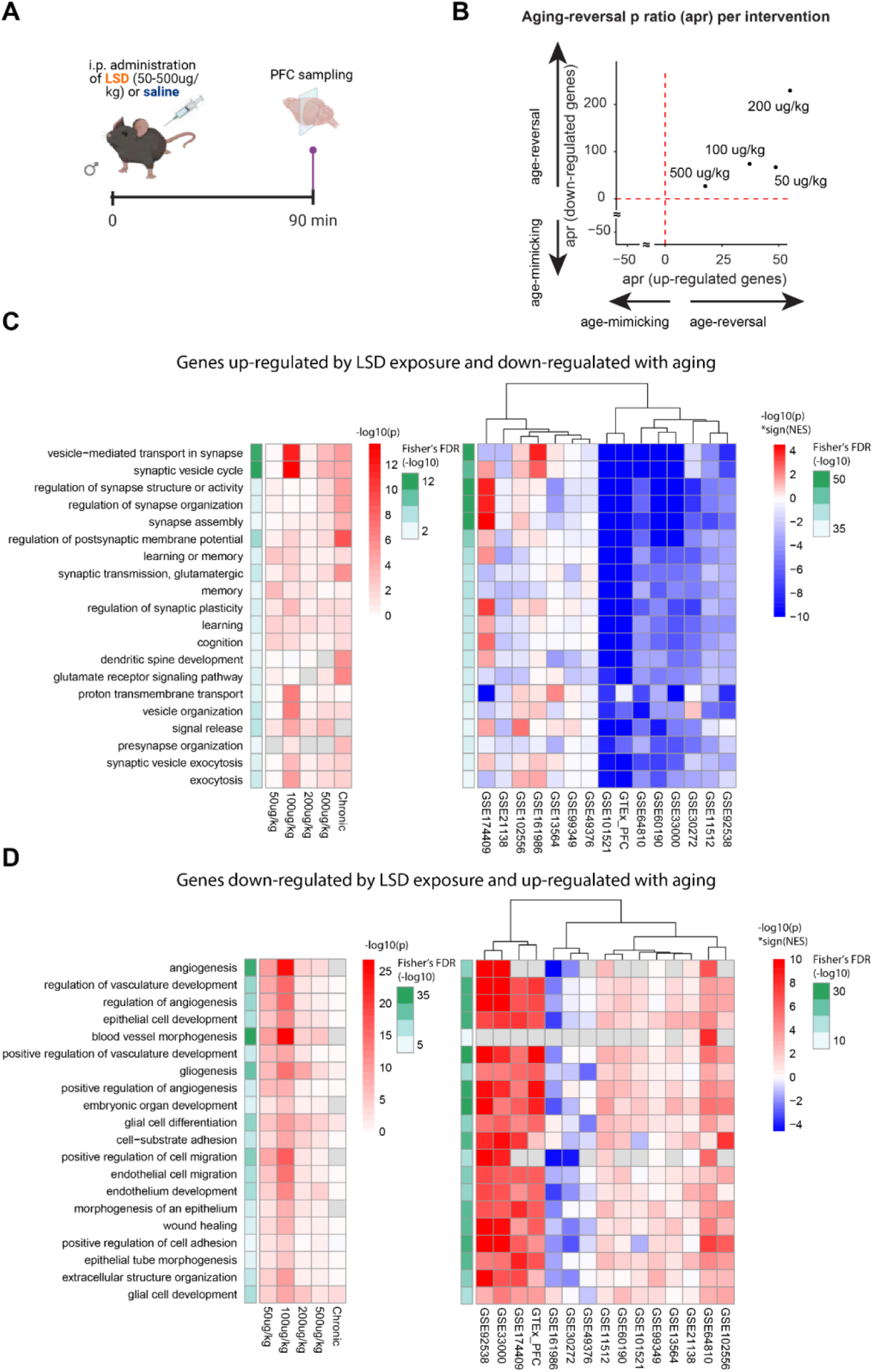
A. Schematic of in vivo treatment and sampling. B. Summary of transcriptional age-signature reversal and mimicking of acute LSD treatment at different dosages in mice. Each point corresponds to the effect in mice’s prefrontal cortex of a different dose of LSD, with coordinates on the x and y axis indicating, respectively, the age-signature reversal potential of up-regulated and down-regulated genes, as in Figure 1E. CD. Top 20 Gene Ontology (GO) categories significantly enriched among genes exhibiting inverse regulation between LSD treatment and aging: genes upregulated by LSD and negatively correlated with age (C), and genes downregulated by LSD and positively correlated with age (D). Left: the colour in the heatmap indicates the GSEA p-value (-log10) with the sign determined by the normalized enrichment score (NES). Red, highly significant positive enrichments; blue, highly significant negative enrichments. The green bar on the left indicates the overall FDR (-log10) aggregated across dataset with the Fisher’s method. Right: GO enrichment p-value (-log10) for genes up- (C) and down-regulated (D) by LSD upon different treatment regimens. The green bar on the left indicates the overall p-value (-log10) combining the p-values obtained for each regimen. i.p.=intraperitoneal

We further validated these findings using two additional independent aging datasets (**Figure S8A**, **Methods**) and observed that genes regulated by chronic LSD treatment also displayed expression changes opposite to those associated with increasing Alzheimer’s disease severity (**Figure S8B**).

### LSD induced aging-signature reversion might impact synaptic and dendritic programs

To elucidate the biological processes underlying LSD-associated age-signature reversal, we performed a Gene Ontology (GO) enrichment analyses on genes showing inverse regulation between LSD treatment and aging (i.e., genes upregulated by LSD and negatively correlated with age, and vice versa). Consistent with prior studies of brain aging (Ham & Lee 2020), GO-categories related to synapse- and dendrite-related processes - typically downregulated with aging - were enriched among genes upregulated by LSD (**Figure 2C**), supporting the biological coherence of the reversal signal (**Table S13**). Conversely, categories associated with angiogenesis and gliogenesis, which are known to increase with aging, were preferentially downregulated following LSD treatment (**Figure 2D**, **Table S14**). Comparable patterns were observed when repeating the analysis using dementia-associated transcriptional signatures instead of aging datasets (**Figure S9**, **Tables S15** and **S16**), indicating that LSD modulates molecular pathways relevant to both physiological brain aging and neurodegenerative progression.

### LSD rescues Amyloid-β-induced structural and molecular deficits in primary cortical neurons

Converging evidence from Alzheimer’s disease (AD) models indicates that early synaptic and cytoskeletal disruption, rather than overt neuron loss, is a primary driver of cognitive decline. Amyloid-β (Aβ) disrupts dendritic architecture and arborization and synaptic plasticity, downregulating Postsynaptic Density Protein 95 (PSD95), while also perturbing neuronal microtubule organisation and axonal transport (Lopes et al. 2022; Staurenghi et al. 2021). Simultaneously, maladaptive astrocyte activation, characterized by upregulation of Glial Fibrillary Acidic Protein (GFAP) and synaptotoxic signaling (Davis et al. 2021), is induced. Aβ-induced dendritic simplification, loss of synaptic integrity, cytoskeletal disruption and reactive astrocytosis emerge therefore as key mechanistic features of age-associated neurodegenerative pathology (Davis et al. 2021; Lopes et al. 2022).

To test whether LSD modulates these processes, we examined its effects in a well-established in vitro model of Aβ-induced neurodegeneration (Beretta et al. 2020). We cultured primary cortical neurons along with residual astrocytes derived from postnatal day 2 (P2) mice for 20 days and exposed them to 100 nM Amyloid-β (Aβ) for 3h, followed by treatment with LSD or vehicle for an additional 24h. As a proxy for neuronal morphology and arborization, we quantified by immunofluorescence the area positive for the expression of the microtubule-associated protein 2 (MAP2). This revealed a marked reduction in dendritic complexity upon Aβ exposure, whereas subsequent LSD treatment significantly restored neuronal arborization (**Figure 3AB**). Notably, LSD treatment alone did not affect neuronal architecture, indicating a specific rescue of Aβ-induced structural deficits.

**Figure 3.**
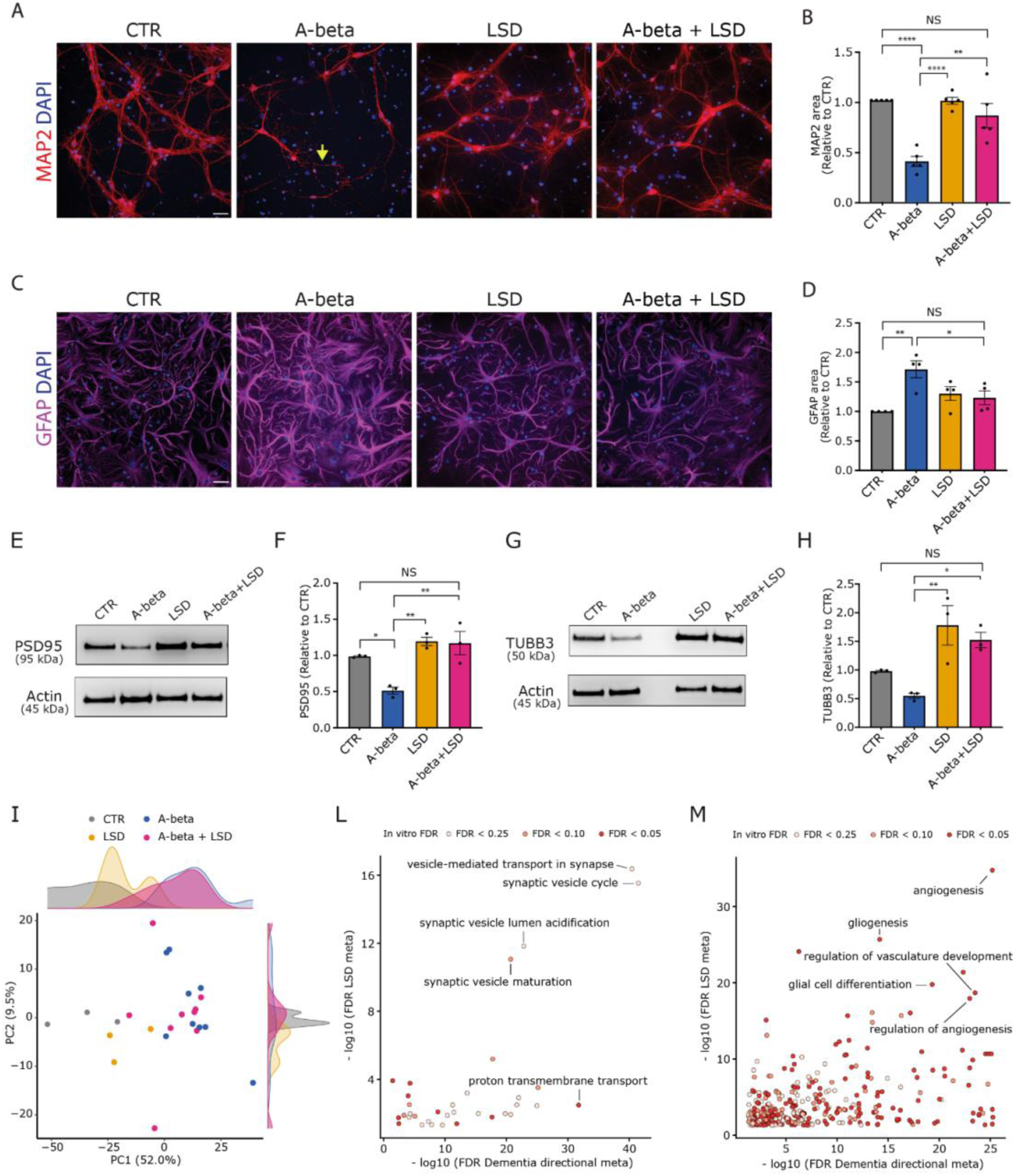
LSD rescues Aβ effects on primary mouse neurons and astrocytes. P2 mouse cortical neurons were grown in culture for 20 days, after which they were treated with 100 nM Aβ for 3 h, followed by 10 µM LSD or vehicle treatment for a further 24 h, generating four different conditions: CTR(no Aβ, DMSO); LSD; Aβ; and Aβ + LSD. A. Immunofluorescence (IF) images for MAP2 (red) along with DAPI staining of treated cortical neurons. Scale bar, 50 µm. The Arrow in Aβ indicates fragmented neuronal processes. B. Quantification of the area of the MAP2 IF signal normalised to the control, average of five independent experiments. Barplots show means ± Standard Error Mean (SEM) of N=5 independent experiments with 4-6 areas of interest imaged per sample; Statistical analyses were performed for indicated comparisons by Ordinary one-way ANOVA(***, p<0.001; **, p<0.01; *, p<0.05). C. IF images for GFAP (magenta) along with DAPI staining of treated astrocytes present in the cortical neuron culture. Scale bar, 50 µm. D. Quantification of the area of the GFAP IF signal normalised to the control, average of four independent experiments, as a measure of astrocyte activation. Barplots show means ± SEM of N=4 independent experiments; Statistical analyses were performed for all comparisons by Ordinary one-way ANOVA; only statistically significant differences are indicated (****, p<0.0001; ***, p<0.001; **, p<0.01). E. Immunoblot for PSD95. Actin was used as a loading control. F. Quantification of the densitometric analysis of PSD95 band normalised to actin band signal and expressed relative to the control average. Barplots show means ± SEM of N=3 independent experiments; Statistical analyses were performed for all comparisons by Ordinary one-way ANOVA; only statistically significant differences are indicated (**, p<0.01; *, p<0.05). G. Immunoblot for bIII-tubulin (TUBB3). Actin was used as a loading control. H. Quantification of the densitometric analysis of bIII-tubulin band, normalised to actin and expressed relative to the control average. Graph shows means ± SEM of N=3 independent experiments; Statistical analyses were performed for all comparisons by Ordinary one-way ANOVA; only statistically significant differences are indicated (**, p<0.01; *, p<0.05) I. Combined Aβ+LSD samples shift toward controls in PCA space defined by Aβ-altered genes. Principal component analysis was performed on genes significantly altered by Aβ. Each point represents one sample, coloured by treatment group. Marginal density plots show the distribution of samples along PC1 and PC2. Axis labels report the percentage of variance explained by each component. LM. Summary scatterplot of GO terms upregulated (L) or downregulated (M) by LSD and oppositely regulated in dementia, showing concordant modulation in genes altered by the addition of LSD to Aβ-treated neurons. Axes report dementia-directional and LSD meta-analysis FDRs, while point colour denotes in vitro significance.

Given that postnatal cortical neuron cultures contain residual astrocytes, we next evaluated astrocyte activation by immunostaining for the astrocytic activation marker GFAP. Aβ treatment induced a significant upregulation of GFAP, whereas additional treatment with LSD significantly reduced it (**Figure 3CD**). Again, LSD alone did not affect GFAP expression.

To determine whether these changes were accompanied by alterations in synaptic integrity, we assessed the expression of the PSD95 synaptic component in the triton-insoluble postsynaptic-enriched fraction by immunoblotting. This revealed that Aβ exposure reduced PSD95 levels, while subsequent LSD treatment restored them to baseline values (**Figure 3EF**). In addition, immunoblot analysis of βIII-tubulin, a neuron-specific tubulin isoform, showed that the Aβ-induced decrease in βIII-tubulin expression was similarly reversed by LSD treatment (**Figure 3GH**). These data indicate that both the disruption of neuronal cytoskeletal and synaptic integrity and the astrocytic activation triggered by Aβ exposure are reinstated by LSD exposure.

### LSD reverses Amyloid-β-induced transcriptional alterations and reinstates neuron-associated gene expression programs

Building on our computational analyses, we investigated whether LSD could reverse Aβ-induced transcriptional alterations in mouse cortical neurons. To this end, we collected RNA from P2 cortical neurons treated with DMSO (control), Aβ at three different concentrations (100 nM, 500 nM, 1 µM), LSD alone, or a combination of Aβ and LSD. RNA sequencing followed by differential expression analysis showed that the number of DEGs (FDR < 0.05) was reduced in the combined treatment compared with Aβ alone (**Figure S10**; paired Wilcoxon test p = 0.13), suggesting that LSD attenuates the transcriptional impact of Aβ. More strikingly, when restricting the analysis to genes dysregulated by Aβ treatment, principal component analysis revealed a gradient separating control from Aβ-treated samples, with combined Aβ + LSD samples clustering closer to controls than Aβ-only samples along both PC1 and PC2 (**Figure 3I**). Of note, Aβ-induced transcriptional signature shows concordance with AD-associated gene expression changes across 4 out of 5 independent Alzheimer’s disease brain datasets (**Figure S11**), supporting the translational relevance of the model.

Notably, Gene Ontology (GO) categories upregulated by LSD in vivo exhibited opposite-direction enrichment as compared to dementia signatures, were reduced in primary neurons treated with Aβ and re-induced by LSD treatment. These categories included synaptic vesicle maturation and transport (**Figure 3L**, **Table S17**). Conversely, GO categories associated with gliogenesis and cell-cell adhesion showed consistent downregulation following LSD treatment both in vivo and in vitro, while displaying upregulation across human dementia datasets (**Figure 3M**).

Collectively, these observations indicate that LSD reinstates a coordinated neuronal transcriptional program while simultaneously suppressing glia- and adhesion-associated responses aberrantly engaged during aging and neurodegeneration. This shift is consistent with a rebalancing of neuron-glia transcriptional states away from reactive or stress-associated programs and toward enhanced functional resilience.

## Discussion

Overall, our findings demonstrate that conserved transcriptional programs associated with brain aging and dementia can be counteracted through targeted pharmacological perturbation. We show that chronic LSD treatment induces gene expression states strongly anti-correlated with aging- and dementia-associated programs in the prefrontal cortex, that this effect is reproducible across species and datasets, it is specific compared to a broad panel of pharmacological perturbations, and linked to functional rescue of amyloid-β-induced structural and molecular alterations in primary cortical neurons.

The specificity of LSD’s age-reversal signature relative to 193 CMap compounds tested in neural progenitor cells argues against a generic perturbational effect and is consistent with the engagement of defined molecular programs rather than a nonspecific transcriptional response.

The comparative analysis across psychoplastogens, entactogens, and dissociative compounds further reveals that age-reversal capacity is not a shared property of psychoactive substances, but varies substantially across pharmacological classes. Notably, MDMA exhibited the strongest age-mimicking transcriptional profile among all compounds tested. However, a direct mechanistic comparison across compounds is complicated by the fact that the available datasets differ substantially in administration regimen, dose, and post-treatment sampling timepoint: variables that influence the transcriptional response to psychoactive compounds.

A systematic study using matched protocols will be necessary to determine whether the observed differences in age-reversal potential reflect intrinsic pharmacological properties or are partly attributable to these experimental factors. Whether the contrasting transcriptional profiles of LSD and MDMA map onto their divergent receptor pharmacology - direct 5-HT2A agonism versus transporter-mediated monoamine release - remains to be established, and represents an important question for future mechanistic studies and potential therapeutic developments. Indeed, our data are correlational in nature and do not allow us to causally attribute the LSD-associated transcriptional reversal to a specific receptor or downstream pathway.

GO analysis indicates that LSD-associated age-signature reversal involves two complementary processes: the upregulation of synapse- and dendrite-related pathways, which typically decline with aging, and the downregulation of gliogenesis and angiogenesis programs, which are instead upregulated with aging. This pattern is consistent with a rebalancing of the neuron–glia transcriptional landscape away from the reactive and degenerative states that characterise the aging brain. Critically, the GO categories modulated by LSD are not only anti-correlated with aging signatures but are also dysregulated in dementia: categories upregulated by LSD in vivo are suppressed across human dementia datasets, while categories downregulated by LSD in vivo are elevated in dementia: a correspondence directly confirmed by the in vitro data, where the addition of LSD to Aβ-treated neurons concordantly reinstated the LSD-upregulated, dementia-suppressed, programs while attenuating the LSD-downregulated and dementia-elevated ones. This three-way convergence - in vivo LSD treatment, human dementia transcriptomics, and an Aβ in vitro model - strengthens the interpretation that LSD engages programs of direct relevance to neurodegenerative pathology, rather than merely opposing statistical aging signatures. Together, these findings link systems-level gene expression changes to concrete cellular outcomes under neurodegeneration-relevant stress.

Several limitations should be considered. The transcriptional datasets employed for the computational analyses derive from bulk tissue, and the cell-type composition of the prefrontal cortex changes with age; single-cell resolution data will be necessary to attribute the reversal signal to specific neuronal and glial populations. The in vitro validation model uses cortical neurons derived from postnatal day 2 mice, which do not fully recapitulate the transcriptional state of mature or aged neurons. Finally, our data do not resolve whether the transcriptional reversal reflects a durable reprogramming of gene expression or a transient response, a distinction that will be critical for assessing its functional and therapeutic significance. Nevertheless, these results motivate further investigations into the molecular mechanisms through which LSD modulates brain aging and into its potential translational relevance. At the same time, although recent studies suggest that psychedelics are safe and well tolerated in older adults (Bouchet et al. 2024), the limited sample sizes examined to date may not capture rare adverse events. Careful evaluation of these factors will be essential as this line of research advances toward clinical evaluation.

All data and results presented in this study are freely available and queryable through an open-source Rshiny app (https://polilab.shinyapps.io/agereversal/), which also allows testing the brain age-signature reversion potential of any treatment for which transcriptional response signatures are available. The underlying code is available at AuroraSavino90/AgingVsLSD (github.com).

## Methods

### Aging datasets collection and curation

We curated human PFC aging transcriptomic datasets from GEO and GTEx, harmonized gene and metadata annotations, applied dataset-specific preprocessing and quality filtering, and retained suitable DLPFC/PFC datasets for age-correlation analyses, as detailed in the **Supplementary Methods**.

### Integration of aging datasets

As dorsolateral prefrontal cortex (DLPFC) samples from healthy subjects were the most represented across our curated aging datasets (**Figure S12C**), we focused on this subset of samples for data integration. Particularly, we collated the transcriptomic profiles of 681 DLPFC samples from healthy individuals available across 11 datasets, focusing on a set of 4,893 genes shared among these datasets. We observed a pronounced batch effect, which hinged the association between gene expression and sample age and was reflective of the diverse platforms used to acquire the different datasets (**Figure S2A-C**). We identified latent variables with SVA (surrogate variable analysis, removing unwanted variation in high-throughput experiments) (Leek et al. 2012), and removed them from the original data employing a linear model. This reduced the batch effect and resulted in the first principal component of the integrated dataset significantly correlating with subjects’ age (Pearson’s R = 0.60, p-value < 2.2 × 10^−16^, **Figure S2D-F**).

### Psychoplastogens’ datasets

We collected transcriptomics datasets of rodents’ PFC treated with psychedelics, entactogens and dissociative compounds (all described as “psychoplastogens”, **Table S4**). For RNA-seq datasets, we identified differentially expressed genes upon treatment using the DESeq2 R package (Love et al. 2014), while we used the Limma R package (Ritchie et al. 2015) for microarrays (FDR<0.05). We derived human homologs of mouse and rat’s genes from HomoloGene (NCBI Resource Coordinators 2016) on the 28/03/2022 for rat genes and the 29/08/2022 for mouse genes.

### Selection of Positive and negative controls

As controls for interventions known to induce or counteract brain aging effects, we chose exercise and exposure to an enriched environment (age-reverting), and ethanol administration (age-inducing), deriving signatures of differentially expressed genes (FDR < 0.05) upon intervention from the following GEO datasets: GSE64607 (Hill & Gammie 2022) (selecting cortical samples, control vs voluntary running for 28 days), GSE164798 (Wierczeiko et al. 2021) (selecting C57BL/6 mice, control vs voluntary running for 30 days), GSE111273 (control vs enriched environment, separately for young and old mice), GSE72507, GSE60676 (Osterndorff-Kahanek et al. 2015) (selecting prefrontal cortex samples, mice analyzed at the end of ethanol/control exposure), GSE28515 (Wolen et al. 2012). As an additional negative control, we included signatures of aging in rats, the model organism used in our primary psychedelics dataset (dataset GSE75772 (Ianov et al. 2016)).

### Identification of genes whose expression correlates with age

We used samples from the DLPFC or mPFC of healthy subjects in the curated aging datasets, discarding datasets with less than 10 samples with annotated age ≥ 20 years. We computed Pearson’s correlation scores between each gene’s expression profile and the pattern of annotated ages, across samples.

### Testing age-signature reversal potential

We ranked all the genes in each sample of the aging dataset based on the correlation of their expression with age (computed as explained above), in decreasing order, i.e. with genes whose expression was strongly correlated with age at the top. We used the resulting ranked lists of genes as background ranked lists for a Gene Set Enrichment Analysis (GSEA) (Subramanian et al. 2005) using sets of differentially expressed genes (up- or down-regulated, used separately) upon psychoplastogen-treatment/exposure-to-enriched-environment/exercise as query signatures. This analysis yielded enrichment scores (ES) with corresponding significance p-values, reflecting the extent to which the genes in the query signatures were ranked consistently across the background lists, i.e. down-regulated genes at the top and up-regulated genes at the bottom (with large positive ES yielded by down-regulated genes’ query signatures and large negative ES yielded by up-regulated genes’ query signatures indicating strong consistency, thus a potential age-reversal effect).

In this way, we obtained an age-reversal p-value for each treatment and each aging dataset separately for up- and down-regulated genes by the treatment, which we then aggregated across datasets using the Fisher method (Edwards 2005), thus computing a global reliability score for age-reversal effects of each intervention (still keeping up- and down-regulated genes upon treatment separated).

### Comparing the age-signature-reversal/-mimicking potential of psychoplastogen with that of positive and negative controls

We quantified the age-signature reversal and mimicking potential of each intervention by testing enrichment of treatment-induced gene sets along aging-associated ranked lists using a directional GSEA framework, followed by aggregation across datasets and comparison of reversal versus mimicking signals, as detailed in the **Supplementary Methods**.

### Connectivity map drug analysis

We downloaded the Connectivity Map (cMAP) (Subramanian et al. 2017) dataset from Gene Expression Omnibus (GSE70138), from the GSE70138_Broad_LINCS_Level4_ZSPCINF_mlr12k_n345976×12328.gctx file containing the z-scores resulting from differential expression of each gene (both measured and inferred) between cells treated with a compound and control cells treated with the vehicle, usually DMSO. We considered only perturbations of neural progenitor cells (NPC), as indicated in cMAP’s metadata, and in case the same perturbation was repeated on multiple samples, we considered the average z-score for each gene. For each drug, we defined a drug signature by taking the top differentially expressed genes upon the drug’s treatment based on the average z-score, and selecting the same number of genes that we found differentially expressed upon LSD treatment. We then applied the procedure described in the “Testing age-signature reversal and mimicking” section of the Methods to assess the transcriptional age-reversal effect of these drugs (**Figures S1 and S2**, **Figure 1CD**).

### Testing psychoplastogens potential in reverting signatures of dementia

We assessed the ability of psychoplastogens to reverse dementia-associated transcriptional signatures by comparing treatment-induced gene expression programs with human PFC dementia datasets using a single-sample GSEA framework, followed by aggregation and directional comparison of reversal versus mimicking effects across datasets, as detailed in the **Supplementary Methods**.

### External validations

As external validations, we repeated the testing of chronic LSD age-signature reversal potential defining age-related genes based on additional datasets not found querying GEO but identified through a literature search (GSE25219 and GSE36192). In addition, we tested the dementia-reversal potential on the Mount Sinai Brain Bank (MSBB) dataset, collecting brain samples from Alzheimer patients, annotated for disease severity. In this case, we selected frontal pole samples from subjects older than 90 years, and ranked the genes based on their correlation with the Braak score (a measure of Alzheimer disease severity) to perform the GSEA on LSD-regulated genes.

### In vivo experiments

All experimental procedures were conducted in agreement with the Italian Legislation after approval by the Ministry of Health (E5FC2.N.FK0 and CC652.173.EXT.78). Animals used for this study were kept either at the specific pathogen-free transgenic unit of the Molecular Biotechnology Center at the University of Turin (for the in vivo LSD treatment experiments) or in standardized hygienic conditions in the Preclinical research facility of Human Technopole in Milan (for the in vitro experiments). The animals had free access to food and water under 12h/12h light/dark cycle. All experiments were performed on C57Bl/ 6J mice. In vivo experiments were performed in 8 week old males and the in vitro experiments were performed in P0-P2 mice, whose sex was not determined as it is not likely to be of relevance for the results obtained in the present study.

8 weeks old male C57BL/6 mice were intraperitoneally injected with either saline (0.9% NaCl) vehicle or LSD at either 50 ug/kg, 100 ug/kg, 200 ug/kg, or 500 ug/kg and euthanized after 90 minutes through cervical dislocation. The animals were monitored throughout the treatment period and the treatment was considered effective based on the presence of head-twitch response (Halberstadt & Geyer 2013). Procedures were conducted at the Molecular Biotechnology Center (University of Turin) in conformity with national and international laws and policies as approved by the Faculty Ethical Committee and the Italian Ministry of Health.

### RNA-sequencing (in vivo LSD treatment)

Brains were removed and sectioned on a vibratome. The range of sections used for analysis corresponds to coronal sections 25-43 of the reference Allen Brain Atlas (Allen Reference Atlas–Mouse Brain [brain atlas]. Available from atlas.brain-map.org). The PFC areas were isolated and selected following published protocols (Laubach et al. 2018). Total RNA was extracted using Trizol reagent according to manufacturer instructions.

RNA Integrity was quantified with the TapeStation automated electrophoresis system, standard mRNA stranded Illumina protocols were employed for library preparation and 40 Million Paired-end 50bp reads per sample were sequenced.

We checked the quality of the sequencing with FastQC (https://www.bioinformatics.babraham.ac.uk/projects/fastqc/), aligned the reads to the Grcm38 mouse genome with hisat2 (Kim et al. 2019) and counted the reads aligned to each gene with HTSeq (Anders et al. 2015).

We filtered the genes with at least 10 reads in at least 3 samples, removed the genes with variance < 0.5 after normalising as reads per million, and computed the differentially expressed genes between treated and controls with DESeq2 (Love et al. 2014).

### Primary cortical neuron culture

Primary cortical neurons were obtained from 0-2 day-old pups, as described previously (Sclip et al. 2013). Brielfy, animals were euthanized and cortex were isolated under a surgical stereomicroscope. Tissues were digested with papain (200 U/ml) in CNDM medium (5.8 mM MgCl2, 0.5 mM CaCl2, 3.2 mM HEPES, 0.2 mM NaOH, 30 mM K2SO4 and 90 mM Na2SO4. pH 7.4, 292 mOsm) supplemented with 0.4% glucose for 30 min at 34°C. Enzymatic activity was blocked with trypsin inhibitors (10 μg/ml, Sigma) in CNDM plus 0.4% glucose for 45 min at room temperature (RT), and the tissue was mechanically dissociated in minimal essential medium (MEM; Invitrogen) supplemented with 10% fetal bovine serum (FBS) (HyperClone) and 0.4% glucose; 0.75 × 105 cells per cm2 were plated on poly-L-lysine-coated (0.1 mg/ml) IBIDI (for immunofluorescence) and 35mm dishes (for RNA extraction and Biochemical analysis). After attachment, the culture medium was switched to Neurobasal-A (Invitrogen) supplemented with 2% B27-plus (Invitrogen), 200 mM glutamine and 100U/ml PenStrep. The medium was changed after 7 days in culture, once a week. No antimitotic agents (e.g., cytosine β-D-arabinofuranoside or aphidicolin), were added to suppress glial proliferation, hence the culture contained a residual amount of astrocytes.

The treatments were performed at 18-20 days in vitro (DIV), when neurons are considered to be differentiated. Neurons were pre-treated with Abeta (Bachem) oligomers at 100nm for 3h, as a synaptotoxic stimuls. After a wash-out neurons were treated with 10 uM LSD (LGC Standards) for 24 hours.

### Immunofluorecence

Cortical neurons plated on IBIDI were fixed in 4% paraformaldehyde (PFA) and 4% sucrose in phosphate buffered saline (PBS) for 8 min. For immunofluorescence the following antibodies were used: chicken anti-MAP2 (Abcam, 1:10000), rabbit anti-GFAP (Dako, 1:3000). Secondary antibodies were conjugated with Alexa-488, Alexa-555 fluorophores (Invitrogen).

### Microscopy and image analysis

Images were acquired using Plan Apo Lambda 20x/0.75 NA objective at confocal microscope system Nikon Ti2E inverted microscope equipped with a CrestOptics X-Light V3 spinning disk unit. Using Fiji ImageJ software the amount of Map2 signal was quantified as percentage of cover area and the amount of GFAP signal was measured as integrated intensity.

### Subcellular fractionation

For the subcellular PSD-enriched fractionation, cortical neurons plated on 35mm dish were homogenized in 0.32M ice-cold sucrose buffer containing: 1mM HEPES pH7.4, 1mM MgCl2, 1mM EDTA, 1mM NaHCO3, and 0.1 PMSF, in presence of a complete set of protease inhibitors (Complete; Roche Diagnostics, Basel, Switzerland) and phosphatase inhibitors (Sigma, St. Louis, MO). Samples were centrifuged at 1,000xg for 10 min. The resulting supernatant (S1) was centrifuged at 3,000xg for 15min to obtain a crude membrane fraction (P2 fraction). The pellet was resuspended in 1mM HEPES plus protease and phosphatase inhibitor and centrifuged at 100,000xg for 1h at 4°C. The pellet (P3) was resuspended in buffer containing 75mM KCl and 1% Triton X-100, incubating 20min on ice and centrifuged at 100,000xg for 1h at 4°C. The final pellet (P4) referred to as triton-insoluble postsynaptic fraction (TIF), was resuspended in 20mM HEPES, sonicated 3 times and stored at -80°C until processing.

### Immunoblot

Protein concentrations were quantified using the BCA pierce and 5μg of TIF extracted proteins were separated by 4-12% SDS polyacrylamide gel electrophoresis (Invitrogen). Nitrocellulose membranes were blocked in Tris-buffered saline (5% no fat milk powder, 0.1% Tween20) (1h, room temperature). After blocking with 5% dry milk for 1 h at room temperature, membranes were incubated overnight at 4°C with rabbit anti PSD95(1:2000, Invitrogen), mouse Beta III tubulin (1:5000, Santa Cruz) and mouse anti Actin (1:20000, Millipore), then with HRP-conjugated secondary antibodies (BioRad) 1 h at room temperature. Peroxidase activity was detected using ECL (BioRad) and visualized with a ChemiDoc imaging system (BioRad). Image Lab software (Bio-Rad) was used for quantitative densitometry of protein bands.

### RNA-sequencing (in vitro)

RNA was extracted using the Qiagen RNeasy Plus MiNI kit and RNA integrity was measured using an Agilent 4200 Tapestation with an RNA Integrity Number ≥9. cDNA libraries were prepared with 150ng of using the Illumina Stranded mRNA Prep, Ligation Kit (Illumina, Inc., San Diego, CA). The libraries were amplified via 14 cycles of PCR and sequenced on a NovaSeq 6000, PE 100, generating an average number of 45 million reads per sample.

Sequences were processed using nf-core/rnaseq v3.18, with GRCm38 from iGenomes as the reference genome. Briefly, reads were trimmed to remove adapter sequences and low-quality bases from read ends using Trimgalore v0.6.10, then aligned to the reference genome using STAR v2.6.1d. The resulting alignments were processed with Salmon v1.10.3 for transcriptome quantification. Library reverse strandedness was assessed using RSeQC v5.0.2.

Ensembl IDs were then mapped to gene symbols: when multiple entries mapped to the same symbol, the entry with the highest mean expression across samples was retained. Rows lacking a valid symbol or containing missing symbols were removed. Principal component analysis (PCA) was performed using FactoMineR (**Figure S14**).

To mitigate systematic differences across experimental replicates, batch correction was applied to the gene-level counts matrix using ComBat-seq (sva package), specifying Replicate as batch. Corrected counts were subsequently converted to library-size normalized values as reads per million (RPM). PCA was repeated on normalized RPM values to assess the impact of correction (**Figure S14**). After batch correction, genes were filtered to retain those expressed at ≥10 counts in at least 3 samples.

Differential expression was performed with DESeq2. For Aβ dose–response comparisons against control, a DESeqDataSet was constructed from the subset including the controls and the three Aβ concentrations. Separate contrasts were extracted for each Aβ concentration versus control.

Aβ-responsive genes were defined as the union of significant differentially expressed genes (FDR < 0.05) across the three Aβ concentrations, separated into upregulated (log2FC>0) and downregulated (log2FC<0) sets. To evaluate whether LSD attenuates the Aβ transcriptional signature, PCA was performed on RPM values restricted to the union of Aβ-upregulated and Aβ-downregulated genes.

To assess whether an Aβ-induced transcriptional program observed in vitro is recapitulated in human Alzheimer’s disease (AD) brain transcriptomes, we performed per-dataset differential expression analyses and gene set enrichment testing across multiple independent cohorts (**Table S8**). Differential expression between AD and control samples was computed independently for each dataset using limma. Genes were ranked by the differential expression log2 fold change (AD vs control), and enrichment was tested separately for the Aβ up and down gene sets after mapping mouse gene identifiers to human orthologs using biomart.

### Functional enrichment analysis

We performed Gene Ontology enrichment analyses across human aging and dementia datasets, in vivo LSD perturbations, and in vitro Aβ/LSD-treated neurons to identify biological processes associated with age- and dementia-related transcriptional changes and to assess their modulation by LSD, integrating results across datasets and testing for concordant or opposing regulation, as detailed in the **Supplementary Methods**.

### ShinyApp

We built a web tool to test the age-reversal potential of a set of user-defined genes. Users can select the conditions to compute the relationship between gene expression and age, setting the age range, gender, ethnicity, health status and brain region: the same human brain transcriptomic data employed in this work are used for this analysis. Plots equivalent to **Figure 1B** and **Figure 1E** can be obtained through this web tool, following the same steps described in the section “Testing age reversal” of the Methods. Similarly, an age-reversal score can be obtained as the ratio of the age-reversal and age-mimicking p-values (as described in the “Testing age reversal” section), scaled to [-1,1], where 1 indicates maximum age reversal while -1 indicates maximum age mimicking potential. The tool is freely accessible at the link https://polilab.shinyapps.io/agereversal/.

## Code availability

The code to reproduce the analyses and generate the figures/tables presented in this work is deposited at https://github.com/AuroraSavino90/AgingVsLSD. R and Package versions are indicated in **Supplementary File 2**.

## Declaration of generative AI and AI-assisted technologies in the writing process

During the preparation of this work, the authors used ChatGPT (OpenAI) to assist with language editing and proofreading, with the aim of improving clarity, grammar, and readability. No content generation, data analysis, or scientific interpretation was performed by the tool. All authors reviewed, revised, and approved the final version of the manuscript and take full responsibility for its content.

## Data availability statement

RNA-sequencing data generated in this work were deposited in the Gene Expression Omnibus and are accessible at https://www.ncbi.nlm.nih.gov/geo/ under the accession numbers GSE263662 and GSE326966. All the other data is publicly available and accessible as detailed in the Methods and the Supplementary Methods.

## Author Contribution Statement

A.S. and F.I designed the study and wrote the original draft of the manuscript. A.S. performed the computational analyses. C.L, S.R., M.C. performed in vivo LSD treatments and collected brain samples. G.R.M. and L.A. supervised the in vivo experiments. I.B. performed in vitro experiments on primary cortical neurons, immunofluorescence and immunoblots. F.I, N.K. and V.P. supervised the project. All authors reviewed, revised, and approved the final version of the manuscript.

## Acknowledgements

We are grateful to the following facilities of Human Technopole: National facility for Genomics, Light Imaging and Preclinical research facility. We thank Parker Kelley for providing transcriptomics data of rats’ brains upon pharmahuasca, harmaline and DMT treatment, and Noèlia Fernàndez-Castillo for sharing the transcriptomics data of mice brains upon MDMA. We thank Oliver Harschnitz for providing feedback on the manuscript.

Research in the Kalebic lab is supported by grants from AIRC (MFAG 2022 ID 27157) and Gilbert Family Foundation (#923004 and #925001) to N.K.

