## Supplementary File 1 for "Lysergic acid diethylamide reverses aging- and neurodegeneration-associated brain transcriptional programs"

1. Molecular Signatures Underlying Selective Regional Vulnerability to Alzheimer's Disease
(Submitter supplied) Alzheimer's disease (AD) is the most common form of dementia, characterized by progressive cognitive impairment and neurodegeneration as a result of abnormal neuronal loss. To elucidate the molecular systems associated with AD, we characterized the gene expression changes associated with multiple clinical and neuropathological traits in 1,053 postmortem brain samples across 19 brain regions from 125 persons dying with varying severities of dementia and variable AD-neuropathology severities.
Organism:	Homo sapiens
Type:		Expression profiling by array
Platforms: GPL570 GPL96 GPL97 2004 Samples
FTP download: GEO (CEL) ftp://ftp.ncbi.nlm.nih.gov/geo/series/GSE84nnn/GSE84422/
Series		Accession: GSE84422	ID: 200084422

2. Zika Virus Disrupts Phospho-TBK1 Localization and Mitosis in Human Neural Stem Cell Model Systems
(Submitter supplied) Purpose: Use single cell RNA-seq technology on neural epithelial stem cell culture to understand the composition and transcriptome property of the model system. The culture system was then used to study ZIKV infection. Methods: Single cell RNA-seq and bioinformatic analysis on NES cells in culture and single cells from  5-6 post conception week (pcw), 16 pcw, as well as 19-20 pcw human fetal neocortex. more...
Organism:	Homo sapiens
Type:		Expression profiling by high throughput sequencing
Platform: GPL11154 1608 Samples
FTP download: GEO (TXT) ftp://ftp.ncbi.nlm.nih.gov/geo/series/GSE81nnn/GSE81475/
SRA Run Selector: https://www.ncbi.nlm.nih.gov/Traces/study/?acc=PRJNA321741
Series		Accession: GSE81475	ID: 200081475

3. Gene expression profiles of human prefrontal cortex brain tissues
(Submitter supplied) Dissecting the shared etiology of different diseases could benefit from a systematic search for associated molecules and their interactions. We investigated genome-wide disruptions in the co-regulation of genes in two neurodegenerative diseases, Alzheimer's or Huntington's disease (AD or HD), using expression profiles from postmortem prefrontal cortex samples of 624 demented patients and non-demented control individuals with matched genotype and clinical data. more...
Organism:	Homo sapiens
Type:		Expression profiling by array
Platform: GPL4372 624 Samples
FTP download: GEO (TXT) ftp://ftp.ncbi.nlm.nih.gov/geo/series/GSE33nnn/GSE33000/
Series		Accession: GSE33000	ID: 200033000

4. The effects of aging on circadian patterns of gene expression in the human prefrontal cortex
(Submitter supplied) With aging, significant changes in circadian rhythms occur, including a shift in phase toward a “morning” chronotype and a loss of rhythmicity in circulating hormones. However, the effects of aging on molecular rhythms in the human brain have remained elusive. Here we employed a previously-described time-of-death analyses to identify transcripts throughout the genome that have a significant circadian rhythm in expression in the human prefrontal cortex (Brodmann’s areas (BA) 11 and 47). more...
Organism:	Homo sapiens
Type:		Expression profiling by array
Platform: GPL11532 420 Samples
FTP download: GEO (CEL, TXT) ftp://ftp.ncbi.nlm.nih.gov/geo/series/GSE71nnn/GSE71620/
Series		Accession: GSE71620	ID: 200071620

5. GPR56/ADGRG1 is associated with response to antidepressant treatment
(Submitter supplied) It remains unclear why many patients with depression do not respond to antidepressant treatment. In three cohorts of individuals with depression and treated with serotonin-norepinephrine reuptake inhibitor (N=424) we show that responders, but not non-responders, display an increase of GPR56 mRNA in the blood. In a small group of subjects we also show that GPR56 is also downregulated in the PFC of individuals with depression that died by suicide. more...
Organism:	Homo sapiens
Type:		Expression profiling by array
Platform: GPL10558 406 Samples
FTP download: GEO (CSV) ftp://ftp.ncbi.nlm.nih.gov/geo/series/GSE146nnn/GSE146446/
Series		Accession: GSE146446	ID: 200146446

6. Epigenetic inactivation of the autophagy-lysosomal system in the Parkinson’s disease appendix
(Submitter supplied) This SuperSeries is composed of the SubSeries listed below.
Organism:	Homo sapiens
Type:		Expression profiling by high throughput sequencing; Genome binding/occupancy profiling by high throughput sequencing; Methylation profiling by high throughput sequencing
Platforms: GPL18573 GPL24676 GPL16791 366 Samples
FTP download: GEO (TXT) ftp://ftp.ncbi.nlm.nih.gov/geo/series/GSE135nnn/GSE135751/
Series		Accession: GSE135751	ID: 200135751

7. Inference of cell-type composition from human brain transcriptomic datasets illuminates the effects of age, manner of death, dissection, and psychiatric diagnosis
(Submitter supplied) Most neuroscientists would agree that psychiatric illness is unlikely to arise from pathological changes that occur uniformly across all cells in a given brain region. Despite this fact, the majority of transcriptomic analyses of the human brain to date are conducted using macro-dissected tissue due to the difficulty of conducting single-cell level analyses on donated post-mortem brains. To address this issue statistically, we compiled a database of several thousand transcripts that were specifically-enriched in one of 10 primary brain cell types identified in published single cell type transcriptomic experiments. more...
Organism:	Homo sapiens
Type:		Expression profiling by array
Platforms: GPL17027 GPL10526 363 Samples
FTP download: GEO (CEL, TXT) ftp://ftp.ncbi.nlm.nih.gov/geo/series/GSE92nnn/GSE92538/
Series		Accession: GSE92538	ID: 200092538

8. Sex-specific Transcriptional Signatures in Human Depression
(Submitter supplied) Rational: Major depressive disorder (MDD) is a leading cause of disease burden worldwide. While the incidence, symptoms and treatment of MDD all point toward major sex differences, the molecular mechanisms underlying this sexual dimorphism remain largely unknown. Methods: Here, combining differential expression and gene coexpression network analyses, we provide a comprehensive characterization of male and female transcriptional profiles associated with MDD across 6 brain regions. more...
Organism:	Homo sapiens; Mus musculus
Type:		Expression profiling by high throughput sequencing
Platforms: GPL11154 GPL13112 341 Samples
FTP download: GEO (TXT) ftp://ftp.ncbi.nlm.nih.gov/geo/series/GSE102nnn/GSE102556/
SRA Run Selector: https://www.ncbi.nlm.nih.gov/Traces/study/?acc=PRJNA398031
Series		Accession: GSE102556	ID: 200102556

9. Gene expression of L3 and L5 pyramidal neurons in the DLPFC comparing schizophrenia from bipolar major depressive disorders and unaffected subjects.
(Submitter supplied) Impairments in certain cognitive processes (e.g., working memory) are typically most pronounced in schizophrenia (SZ), intermediate in bipolar disorder (BP) and least in major depressive disorder (MDD). Given that working memory depends, in part, on neural circuitry that includes pyramidal neurons in layer 3 (L3) and layer 5 (L5) of the dorsolateral prefrontal cortex (DLPFC), we sought to determine if transcriptome alterations in these neurons were shared or distinctive for each diagnosis.
Organism:	Homo sapiens
Type:		Expression profiling by array
Platform: GPL13667 286 Samples
FTP download: GEO (CEL) ftp://ftp.ncbi.nlm.nih.gov/geo/series/GSE87nnn/GSE87610/
Series		Accession: GSE87610	ID: 200087610

10. Hemispheric asymmetry in the human brain and in Parkinson’s disease is linked to divergent epigenetic patterns in neurons
(Submitter supplied) Background Hemispheric asymmetry in neuronal processes is a fundamental feature of the human brain and drives symptom lateralization in Parkinson's disease (PD), but its molecular determinants are unknown. Here, we identify divergent epigenetic patterns involved in hemispheric asymmetry by profiling DNA methylation in isolated prefrontal cortex neurons from control and PD brain hemispheres. DNA methylation is fine-mapped at enhancers and promoters, genome-wide, by targeted bisulfite sequencing in two independent sample cohorts. more...
Organism:	Homo sapiens
Type:		Methylation profiling by high throughput sequencing; Expression profiling by high throughput sequencing
Platforms: GPL16791 GPL18573 279 Samples
FTP download: GEO (TAB, TXT) ftp://ftp.ncbi.nlm.nih.gov/geo/series/GSE135nnn/GSE135037/
Series		Accession: GSE135037	ID: 200135037

11. Temporal Dynamics and Genetic Control of Transcription in the Human Prefrontal Cortex
(Submitter supplied) In order to gain a global molecular perspective on how the human genome gives rise to the brain, we explore the temporal dynamics and genetic control of transcription in human dorsolateral prefrontal cortex in an extensive series of brain tissue from fetal development through aging. We discover a wave of gene expression changes occurring during fetal development which are reversed in early postnatal life. more...
Organism:	Homo sapiens
Type:		Expression profiling by array
Platform: GPL4611 269 Samples
FTP download: GEO (TXT) ftp://ftp.ncbi.nlm.nih.gov/geo/series/GSE30nnn/GSE30272/
Series		Accession: GSE30272	ID: 200030272

12. Multi-tissue gene expression profiles of human brain (VC)
(Submitter supplied) The genetics of complex disease produce alterations in molecular interactions of cellular pathways which collective effect may become clear through the organized structure of molecular networks. To characterize molecular systems associated with late-onset Alzheimer´s disease (LOAD), we constructed gene regulatory networks in hundreds of autopsied brain tissues from LOAD patients and non-demented subjects. more...
Organism:	Homo sapiens
Type:		Expression profiling by array
Platform: GPL4372 230 Samples
FTP download: GEO (TXT) ftp://ftp.ncbi.nlm.nih.gov/geo/series/GSE44nnn/GSE44771/
Series		Accession: GSE44771	ID: 200044771

13. Multi-tissue gene expression profiles of human brain (PFC)
(Submitter supplied) The genetics of complex disease produce alterations in molecular interactions of cellular pathways which collective effect may become clear through the organized structure of molecular networks. To characterize molecular systems associated with late-onset Alzheimer´s disease (LOAD), we constructed gene regulatory networks in hundreds of autopsied brain tissues from LOAD patients and non-demented subjects. more...
Organism:	Homo sapiens
Type:		Expression profiling by array
Platform: GPL4372 230 Samples
FTP download: GEO (TXT) ftp://ftp.ncbi.nlm.nih.gov/geo/series/GSE44nnn/GSE44770/
Series		Accession: GSE44770	ID: 200044770

14. Multi-tissue gene expression profiles of human brain (CR)
(Submitter supplied) The genetics of complex disease produce alterations in molecular interactions of cellular pathways which collective effect may become clear through the organized structure of molecular networks. To characterize molecular systems associated with late-onset Alzheimer´s disease (LOAD), we constructed gene regulatory networks in hundreds of autopsied brain tissues from LOAD patients and non-demented subjects. more...
Organism:	Homo sapiens
Type:		Expression profiling by array
Platform: GPL4372 230 Samples
FTP download: GEO (TXT) ftp://ftp.ncbi.nlm.nih.gov/geo/series/GSE44nnn/GSE44768/
Series		Accession: GSE44768	ID: 200044768

15. Low-input and multiplexed microfluidic assay reveals epigenomic variation across cerebellum and prefrontal cortex
(Submitter supplied) There has been an extensive effort underway to profile epigenetic features at the genome-wide scale using primary ex vivo tissues. Cell-type specificity of epigenomes calls for enrichment to obtain a homogenous cell population from a small quantity of tissues. Thus technologies that permit both ultralow input and high throughput are desired for profiling an array of histone marks. Here we demonstrate a simple microfluidic technology, SurfaceChIP-seq, for profiling genome-wide histone modifications using as few as 30 cells per assay and with up to 8 assays running in parallel. more...
Organism:	Homo sapiens; Mus musculus
Type:		Expression profiling by high throughput sequencing; Genome binding/occupancy profiling by high throughput sequencing
Platforms: GPL20301 GPL21103 176 Samples
FTP download: GEO (BW, TXT) ftp://ftp.ncbi.nlm.nih.gov/geo/series/GSE99nnn/GSE99757/
SRA Run Selector: https://www.ncbi.nlm.nih.gov/Traces/study/?acc=PRJNA389549
Series		Accession: GSE99757	ID: 200099757

16. Epigenetic dysregulation of enhancers in neurons is associated with Alzheimer’s disease pathology and cognitive symptoms
(Submitter supplied) Epigenetic control of enhancers alters neuron functions and may be involved in Alzheimer’s disease (AD). Here, we identify enhancers in neurons contributing to AD by comprehensive fine-mapping of DNA methylation at enhancers, genome-wide. We examine 1.2 million CpG and CpH sites in enhancers in prefrontal cortex neurons of individuals with no/mild, moderate, and severe AD pathology (n=101). We identify 1,224 differentially methylated enhancer regions; most of which are hypomethylated at CpH sites in AD neurons. more...
Organism:	Homo sapiens
Type:		Expression profiling by high throughput sequencing; Methylation profiling by high throughput sequencing
Platforms: GPL18573 GPL16791 154 Samples
FTP download: GEO (TXT) ftp://ftp.ncbi.nlm.nih.gov/geo/series/GSE110nnn/GSE110732/
Series		Accession: GSE110732	ID: 200110732

17. Genetic Neuropathology of Obsessive Psychiatric Syndromes
(Submitter supplied) Anorexia nervosa (AN), bulimia nervosa (BN), and obsessive-compulsive disorder (OCD) are complex psychiatric disorders with shared obsessive features, thought to arise from the interaction of multiple genes of small effect with environmental factors.  Potential candidate genes for AN, BN, and OCD have been identified through clinical association and neuroimaging studies; however, recent genome-wide association studies of eating disorders (ED) so far have failed to report significant findings. more...
Organism:	Homo sapiens
Type:		Expression profiling by array
Platform: GPL6947 133 Samples
FTP download: GEO (TXT) ftp://ftp.ncbi.nlm.nih.gov/geo/series/GSE60nnn/GSE60190/
Series		Accession: GSE60190	ID: 200060190

18. The evolution of gene expression levels in mammalian organs
(Submitter supplied) Changes in gene expression are thought to underlie many of the phenotypic differences between species. However, large-scale analyses of gene expression evolution were until recently prevented by technological limitations. Here we report the sequencing of polyadenylated RNA from six organs across ten species that represent all major mammalian lineages (placentals, marsupials and monotremes) and birds (the evolutionary outgroup), with the goal of understanding the dynamics of mammalian transcriptome evolution. more...
Organism:	Gorilla gorilla; Homo sapiens; Mus musculus; Monodelphis domestica; Gallus gallus; Ornithorhynchus anatinus; Macaca mulatta; Pan paniscus; Pan troglodytes; Pongo pygmaeus
Type:		Expression profiling by high throughput sequencing
10 related Platforms 131 Samples
FTP download: GEO ftp://ftp.ncbi.nlm.nih.gov/geo/series/GSE30nnn/GSE30352/
SRA Run Selector: https://www.ncbi.nlm.nih.gov/Traces/study/?acc=PRJNA143627
Series		Accession: GSE30352	ID: 200030352

19. Gene expression in primate postnatal brain through lifespan
(Submitter supplied) This SuperSeries is composed of the SubSeries listed below.
Organism:	Macaca mulatta; Pan troglodytes; Homo sapiens
Type:		Expression profiling by array
Platform: GPL6244 130 Samples
FTP download: GEO (CEL) ftp://ftp.ncbi.nlm.nih.gov/geo/series/GSE22nnn/GSE22570/
Series		Accession: GSE22570	ID: 200022570

20. Large-scale multi-species survey of metabolome and lipidome
(Submitter supplied) This dataset was generated with the goal of comparative study of gene expression in three brain regions and two non-neural tissues of humans, chimpanzees, macaque monkeys and mice. Using this dataset, we performed studies of gene expression and gene splicing evolution across species and search of tissue-specific gene expression and splicing patterns. We also used the gene expression information of genes encoding metabolic enzymes in this dataset to support a larger comparative study of metabolome evolution in the same set of tissues and species.
Organism:	Macaca mulatta; Homo sapiens; Mus musculus; Pan troglodytes
Type:		Expression profiling by high throughput sequencing
4 related Platforms 120 Samples
FTP download: GEO (TSV) ftp://ftp.ncbi.nlm.nih.gov/geo/series/GSE49nnn/GSE49379/
SRA Run Selector: https://www.ncbi.nlm.nih.gov/Traces/study/?acc=PRJNA213747
Series		Accession: GSE49379	ID: 200049379

21. Longitudinal Gene Expression Analysis in Human Brain identifies biological processes underlying neuropathology in Down Syndrome
(Submitter supplied) Trisomy 21 (Ts21) or Down syndrome (DS) is the most common genetic cause of intellectual disability. To investigate the consequences of Ts21 on human brain development, we have systematically analyzed the transcriptome of dorsolateral prefrontal cortex (DFC) and cerebellar cortex (CBC) using exon array mapping in DS and matched euploid control brains spanning from prenatal development to adulthood. We identify hundreds of differentially expressed (DEX) genes in the DS brains, many of which exhibit temporal changes in expression over the lifespan. To gain insight into how these DEX genes may cause specific DS phenotypes, we identified functional modules of co-expressed genes using several different bioinformatics approaches, including WGCNA and gene ontology analysis.  A module comprised of genes associated with myelination, including those dynamically expressed over the course of oligodendrocyte development, was amongst those with the great levels of differential gene expression. Using Ts65Dn mouse line, the most common rodent  model of DS, w e observed significant and novel defects in oligodendrocyte maturation and myelin ultrastructure; establishing  a correlative proof-of-principle implicating myelin dysgenesis in DS.  Thus, examination of the spatio-temporal transcriptome predicts specific cellular and functional events in the DS brain and is an outstanding resource for determining putative mechanisms involved in the neuropathology of DS.
Organism:	Homo sapiens
Type:		Expression profiling by array
Platform: GPL5175 116 Samples
FTP download: GEO (CEL) ftp://ftp.ncbi.nlm.nih.gov/geo/series/GSE59nnn/GSE59630/
Series		Accession: GSE59630	ID: 200059630

22. Epigenomic analysis of Parkinson's disease neurons identifies TET2 loss as neuroprotective
(Submitter supplied) This SuperSeries is composed of the SubSeries listed below.
Organism:	Mus musculus; Homo sapiens
Type:		Expression profiling by high throughput sequencing; Methylation profiling by high throughput sequencing
Platforms: GPL24676 GPL19057 GPL16791 107 Samples
FTP download: GEO (BED, TXT) ftp://ftp.ncbi.nlm.nih.gov/geo/series/GSE136nnn/GSE136010/
Series		Accession: GSE136010	ID: 200136010

23. Human-specific features of special gene expression and regulation in eight brain regions
(Submitter supplied) Despite the substantial efforts made to compile the molecular maps of the human brain, they do not fully inform us of the features unique to humans. Yet, identification of these features might help us to understand both the evolution and nature of the human cognitive traits. Here, we approached this question by analyzing gene expression and H3K27ac chromatin modification data collected in eight brain regions of humans, chimpanzees, gorillas, gibbon and macaques. more...
Organism:	Homo sapiens; Hylobates lar; Gorilla gorilla; Pan troglodytes
Type:		Expression profiling by high throughput sequencing
7 related Platforms 107 Samples
FTP download: GEO (TXT) ftp://ftp.ncbi.nlm.nih.gov/geo/series/GSE100nnn/GSE100796/
SRA Run Selector: https://www.ncbi.nlm.nih.gov/Traces/study/?acc=PRJNA393104
Series		Accession: GSE100796	ID: 200100796

24. Ribo-depleted RNA-Seq Libraries From Huntington's Disease Brain
(Submitter supplied) Transcriptional dysregulation is well documented in Huntington’s disease (HD) post-mortem brain. We previously identified gene expression profiles from both symptomatic and asymptomatic HD prefrontal cortex which characterize the effects of degeneration in disease versus normal brains. Transcriptional dysregulation in HD brain is well documented by these studies and others, but the extent and causes of this dysregulation remain unknown. more...
Organism:	Homo sapiens
Type:		Expression profiling by high throughput sequencing; Other
Platform: GPL18573 100 Samples
FTP download: GEO (CSV) ftp://ftp.ncbi.nlm.nih.gov/geo/series/GSE159nnn/GSE159940/
SRA Run Selector: https://www.ncbi.nlm.nih.gov/Traces/study/?acc=PRJNA670925
Series		Accession: GSE159940	ID: 200159940

25. Transcriptional analysis of multiple brain regions in Parkinson's disease
(Submitter supplied) Transcriptional analysis of multiple brain regions in Parkinson's disease supports the involvement of specific protein processing, energy metabolism, and signaling pathways, and suggests novel disease mechanisms.  This SuperSeries is composed of the SubSeries listed below.
Organism:	Homo sapiens
Type:		Expression profiling by array
Platform: GPL96 93 Samples
FTP download: GEO (CEL, CHP) ftp://ftp.ncbi.nlm.nih.gov/geo/series/GSE20nnn/GSE20295/
Series		Accession: GSE20295	ID: 200020295

26. Repression of Human and Mouse Brain Inflammaging Transcriptome by Broad Gene Body Histone Hyperacetylation
(Submitter supplied) This SuperSeries is composed of the SubSeries listed below.
Organism:	Homo sapiens; Mus musculus
Type:		Expression profiling by high throughput sequencing; Genome binding/occupancy profiling by high throughput sequencing
Platforms: GPL11154 GPL13112 87 Samples
FTP download: GEO ftp://ftp.ncbi.nlm.nih.gov/geo/series/GSE106nnn/GSE106670/
Series		Accession: GSE106670	ID: 200106670

27. Whole-transcriptome brain expression and exon-usage profiling in major depression and suicide
(Submitter supplied) Brain gene expression profiling studies of suicide and depression using oligonucleotide microarrays have often failed to distinguish these two phenotypes. Moreover, next generation sequencing approaches are more accurate in quantifying gene expression and can detect alternative splicing. Using RNA-seq, we examined whole-exome gene and exon expression in non-psychiatric controls (CON, N=29), DSM-IV major depressive disorder suicides (MDD-S, N=21) and MDD non-suicides (MDD, N=9) in the dorsal lateral prefrontal cortex (Brodmann Area 9) of sudden death medication-free individuals post mortem. more...
Organism:	Homo sapiens
Type:		Expression profiling by high throughput sequencing; Non-coding RNA profiling by high throughput sequencing
Platforms: GPL15520 GPL16791 86 Samples
FTP download: GEO (CSV) ftp://ftp.ncbi.nlm.nih.gov/geo/series/GSE101nnn/GSE101521/
SRA Run Selector: https://www.ncbi.nlm.nih.gov/Traces/study/?acc=PRJNA394722
Series		Accession: GSE101521	ID: 200101521

28. Expression profiling of human adult postmortem brain tissue from subjects with bipolar disorder and healthy controls
(Submitter supplied) Bipolar affective disorder is a severe psychiatric disorder with a strong genetic component but unknown pathophysiology. We used microarray technology (Affymetrix HG-U133A GeneChips) to determine the expression of approximately 22 000 mRNA transcripts in post-mortem brain tissue (dorsolateral prefrontal cortex and orbitofrontal cortex) from patients with bipolar disorder and matched healthy controls. more...
Organism:	Homo sapiens
Type:		Expression profiling by array
Platform: GPL96 82 Samples
FTP download: GEO (CEL) ftp://ftp.ncbi.nlm.nih.gov/geo/series/GSE5nnn/GSE5392/
Series		Accession: GSE5392	ID: 200005392

29. Transcriptional alterations in dorsolateral prefrontal cortex and nucleus accumbens implicate neuroinflammation and synaptic remodeling in opioid use disorder
(Submitter supplied) Transcriptional alterations in dorsolateral prefrontal cortex and nucleus accumbens implicate neuroinflammation and synaptic remodeling in opioid use disorder. Transcriptomic profile of 20 control subjects and 20 OUD subjects in brain region DLPFC and NAC
Organism:	Homo sapiens
Type:		Expression profiling by high throughput sequencing
Platform: GPL18573 80 Samples
FTP download: GEO (CSV) ftp://ftp.ncbi.nlm.nih.gov/geo/series/GSE174nnn/GSE174409/
SRA Run Selector: https://www.ncbi.nlm.nih.gov/Traces/study/?acc=PRJNA729761
Series		Accession: GSE174409	ID: 200174409

30. Gene expression changes during primate postnatal brain development
(Submitter supplied) In development, timing is of the utmost importance, and the timing of various developmental processes are often changed during evolution. During human evolution sexual maturation has been delayed relative to other primates and this may have played a critical role for both the increase of human brain size and the rise of human-specific cognitive traits . We measured the timing of gene expression changes in the brains of humans, chimpanzees, and rhesus macaques throughout postnatal development.
Organism:	Pan troglodytes; Macaca mulatta; Homo sapiens
Type:		Expression profiling by array
Platform: GPL6879 80 Samples
FTP download: GEO (CEL, TSV) ftp://ftp.ncbi.nlm.nih.gov/geo/series/GSE11nnn/GSE11512/
Series		Accession: GSE11512	ID: 200011512

31. Gene and miRNA expression data from primate postnatal brain in prefrontal cortex: time course
(Submitter supplied) This SuperSeries is composed of the SubSeries listed below.
Organism:	Homo sapiens; Macaca mulatta
Type:		Expression profiling by array; Non-coding RNA profiling by high throughput sequencing
Platforms: GPL9160 GPL6244 GPL9115 79 Samples
FTP download: GEO (CEL, TXT, XLS) ftp://ftp.ncbi.nlm.nih.gov/geo/series/GSE18nnn/GSE18069/
Series		Accession: GSE18069	ID: 200018069

32. Conserved expression of lincRNA during human and macaque prefrontal cortex development and maturation
(Submitter supplied) 78 tissue samples from prefrontal cortex (PFC) in human and macaque
Organism:	Homo sapiens; Macaca mulatta
Type:		Expression profiling by high throughput sequencing; Non-coding RNA profiling by high throughput sequencing
Platforms: GPL11154 GPL14954 78 Samples
FTP download: GEO (GTF, TSV) ftp://ftp.ncbi.nlm.nih.gov/geo/series/GSE51nnn/GSE51264/
SRA Run Selector: https://www.ncbi.nlm.nih.gov/Traces/study/?acc=PRJNA222268
Series		Accession: GSE51264	ID: 200051264

33. Disruption of human-specific synaptogenesis program in autism [RNA-seq]
(Submitter supplied) To test the connection between the molecular mechanisms underlying autistic disorder and human cognitive evolution, we analyzed the gene expression changes taking place during prefrontal cortex development in autism patients and healthy controls, as well as non-human primates. We found the genes with expression changes in autism are significantly overlapped with genes showing human-specific developmental profile. more...
Organism:	Pan troglodytes; Homo sapiens
Type:		Expression profiling by high throughput sequencing
Platforms: GPL16809 GPL11154 73 Samples
FTP download: GEO (TXT) ftp://ftp.ncbi.nlm.nih.gov/geo/series/GSE59nnn/GSE59288/
SRA Run Selector: https://www.ncbi.nlm.nih.gov/Traces/study/?acc=PRJNA254971
Series		Accession: GSE59288	ID: 200059288

34. mRNA-Seq expression and MS3 proteomics profiling of human post-mortem BA9 brain tissue for Parkinson Disease and neurologically normal individuals
(Submitter supplied) Parkinson disease (PD) is a neurodegenerative disease characterized by the accumulation of alpha-synuclein (SNCA) and other proteins in aggregates termed “Lewy Bodies” within neurons. PD has both genetic and environmental risk factors, and while processes leading to aberrant protein aggregation are unknown, past work points to abnormal levels of SNCA and other proteins. Although several genome-wide studies have been performed for PD, these have focused on DNA sequence variants by genome-wide association studies (GWAS) and on RNA levels (microarray transcriptomics), while genome-wide proteomics analysis has been lacking. more...
Organism:	Homo sapiens
Type:		Expression profiling by high throughput sequencing
Platform: GPL11154 73 Samples
FTP download: GEO (RDATA, TXT) ftp://ftp.ncbi.nlm.nih.gov/geo/series/GSE68nnn/GSE68719/
SRA Run Selector: https://www.ncbi.nlm.nih.gov/Traces/study/?acc=PRJNA283498
Series		Accession: GSE68719	ID: 200068719

35. Age-related changes in the expression of schizophrenia susceptibility genes in the human prefrontal cortex
(Submitter supplied) The molecular basis of complex neuropsychiatric disorders most likely involves many genes. In recent years, specific genetic variations influencing risk for schizophrenia and other neuropsychiatric disorders have been reported. We have used custom DNA microarrays and qPCR to investigate the expression of putative schizophrenia susceptibility genes and related genes of interest in the normal human brain. more...
Organism:	Homo sapiens
Type:		Expression profiling by array
Platform: GPL6893 72 Samples
FTP download: GEO (TXT) ftp://ftp.ncbi.nlm.nih.gov/geo/series/GSE11nnn/GSE11546/
Series		Accession: GSE11546	ID: 200011546

36. Autism-like phenotype and risk gene-RNA deadenylation by CPEB4 mis-splicing
(Submitter supplied) This SuperSeries is composed of the SubSeries listed below.
Organism:	Mus musculus; Homo sapiens
Type:		Expression profiling by array
Platforms: GPL15207 GPL11180 71 Samples
FTP download: GEO (CEL) ftp://ftp.ncbi.nlm.nih.gov/geo/series/GSE113nnn/GSE113842/
Series		Accession: GSE113842	ID: 200113842

37. Network preservation reveals shared and unique biological processes associated with chronic alcohol abuse in the NAc and PFC
(Submitter supplied) This SuperSeries is composed of the SubSeries listed below.
Organism:	synthetic construct; Homo sapiens
Type:		Expression profiling by array; Genome binding/occupancy profiling by high throughput sequencing
Platforms: GPL571 GPL16384 70 Samples
FTP download: GEO (CEL) ftp://ftp.ncbi.nlm.nih.gov/geo/series/GSE161nnn/GSE161999/
Series		Accession: GSE161999	ID: 200161999

38. mRNA-Seq Expression profiling of human post-mortem BA9 brain tissue for Huntington's Disease and neurologically normal individuals
(Submitter supplied) Huntington’s Disease (HD) is a devastating neurodegenerative disorder that is caused by an expanded CAG trinucleotide repeat in the Huntingtin (HTT) gene. Transcriptional dysregulation in the human HD brain has been documented but is incompletely understood. Here we present a genome-wide analysis of mRNA expression in human prefrontal cortex from 20 HD and 49 neuropathologically normal controls using next generation high-throughput sequencing. more...
Organism:	Homo sapiens
Type:		Expression profiling by high throughput sequencing
Platform: GPL11154 69 Samples
FTP download: GEO (TXT) ftp://ftp.ncbi.nlm.nih.gov/geo/series/GSE64nnn/GSE64810/
SRA Run Selector: https://www.ncbi.nlm.nih.gov/Traces/study/?acc=PRJNA271929
Series		Accession: GSE64810	ID: 200064810

39. Gene expression in primate postnatal brain through lifespan - prefrontal cortex
(Submitter supplied) We investigated molecular changes during human, chimpanzee, and rhesus macaque postnatal brain development at the transcriptome, proteome, and metabolome levels in two brain regions: the prefrontal cortex (PFC) that is involved in several human-specific cognitive processes, and the cerebellar cortex (CBC) that may be functionally more conserved. We find a nearly three-fold excess of human-specific gene expression changes in PFC compared to CBC. more...
Organism:	Macaca mulatta; Homo sapiens; Pan troglodytes
Type:		Expression profiling by array
Platform: GPL6244 68 Samples
FTP download: GEO (CEL, TSV) ftp://ftp.ncbi.nlm.nih.gov/geo/series/GSE22nnn/GSE22521/
Series		Accession: GSE22521	ID: 200022521

40. Global epigenomic reconfiguration during mammalian brain development
(Submitter supplied) Whole-genome single-base resolution methylcytosine and hydroxymethylcytosine maps reveal profound changes that occur during frontal cortex development in humans and mice.
Organism:	Homo sapiens; Mus musculus
Type:		Methylation profiling by high throughput sequencing; Expression profiling by high throughput sequencing
Platforms: GPL13112 GPL11154 65 Samples
FTP download: GEO (BAM, TXT) ftp://ftp.ncbi.nlm.nih.gov/geo/series/GSE47nnn/GSE47966/
SRA Run Selector: https://www.ncbi.nlm.nih.gov/Traces/study/?acc=PRJNA208503
Series		Accession: GSE47966	ID: 200047966

41. Gene expression in primate postnatal brain through lifespan - cerebellar cortex
(Submitter supplied) We investigated molecular changes during human, chimpanzee, and rhesus macaque postnatal brain development at the transcriptome, proteome, and metabolome levels in two brain regions: the prefrontal cortex (PFC) that is involved in several human-specific cognitive processes, and the cerebellar cortex (CBC) that may be functionally more conserved. We find a nearly three-fold excess of human-specific gene expression changes in PFC compared to CBC. more...
Organism:	Macaca mulatta; Pan troglodytes; Homo sapiens
Type:		Expression profiling by array
Platform: GPL6244 62 Samples
FTP download: GEO (CEL) ftp://ftp.ncbi.nlm.nih.gov/geo/series/GSE22nnn/GSE22569/
Series		Accession: GSE22569	ID: 200022569

42. Adult postmortem brain tissue (dorsolateral prefrontal cortex) from subjects with bipolar disorder and healthy controls
(Submitter supplied) Bipolar affective disorder is a severe psychiatric disorder with a strong genetic component but unknown pathophysiology. We used microarray technology (Affymetrix HG-U133A GeneChips) to determine the expression of approximately 22 000 mRNA transcripts in post-mortem brain tissue (dorsolateral prefrontal cortex) from patients with bipolar disorder and matched healthy controls. A cohort of 70 subjects was investigated and the final analysis included 30 bipolar and 31 control subjects. more...
Organism:	Homo sapiens
Type:		Expression profiling by array
Dataset: GDS2190 Platform: GPL96 61 Samples
FTP download: GEO (CEL) ftp://ftp.ncbi.nlm.nih.gov/geo/series/GSE5nnn/GSE5388/
Series		Accession: GSE5388	ID: 200005388

43. Gene Expression Profiles in BA46 of Subjects with Schizophrenia and Matched Controls
(Submitter supplied) Results from clinical and imaging studies provide evidence for changes in schizophrenia with disease progression, however, the underlying molecular differences that may occur at different stages of illness have not been investigated.  To test the hypothesis that the molecular basis for schizophrenia changes from early to chronic illness, we profiled genome-wide expression patterns in prefrontal cortex of schizophrenic subjects at different stages of illness, along with their age- and sex-matched controls. more...
Organism:	Homo sapiens
Type:		Expression profiling by array
Platform: GPL570 59 Samples
FTP download: GEO (CEL) ftp://ftp.ncbi.nlm.nih.gov/geo/series/GSE21nnn/GSE21138/
Series		Accession: GSE21138	ID: 200021138

44. RiboZero Gold paired-end RNA-seq data from postmortem tissue of the dorsolateral prefrontal cortex of autism spectrum disorder samples and healthy control samples
(Submitter supplied) Purpose: The goals of this study was to compare the expression of histamine related genes in ASD patients and controls. Methods: RNA-Seq was performed using strand-specific Ribosomal RNA depletion (RiboZero) library preparation and the TruSeq RNA Sample Preparation v2 kit from Illumina. One hundred base pair paired-end sequencing was run on the HiSeq 2000. TopHat (v2.0.4) was used to enforce strand specificity and to align the sequencing reads to known transcripts of the Ensembl Build GRCh37.67. more...
Organism:	Homo sapiens
Type:		Expression profiling by high throughput sequencing
Platform: GPL11154 52 Samples
FTP download: GEO (RDA, XLSX) ftp://ftp.ncbi.nlm.nih.gov/geo/series/GSE102nnn/GSE102741/
SRA Run Selector: https://www.ncbi.nlm.nih.gov/Traces/study/?acc=PRJNA398545
Series		Accession: GSE102741	ID: 200102741

45. Human cortical transcriptome informs brain imaging
(Submitter supplied) We carried out RNA-sequencing (RNA-seq) of adult human postmortem neocortical brain tissue, and then correlated those expression values with the fMRI signal in each brain region
Organism:	Homo sapiens
Type:		Expression profiling by high throughput sequencing
Platform: GPL11154 52 Samples
FTP download: GEO (TXT) ftp://ftp.ncbi.nlm.nih.gov/geo/series/GSE58nnn/GSE58604/
SRA Run Selector: https://www.ncbi.nlm.nih.gov/Traces/study/?acc=PRJNA253002
Series		Accession: GSE58604	ID: 200058604

46. Gene expression data from primate postnatal brain in prefrontal cortex: time course
(Submitter supplied) Gene expression changes determine functional differentiation during development and are associated with functional decline during aging. While developmental changes are tightly regulated, regulation of aging changes is not well established. To assess the regulatory basis of age-related changes and investigate the mechanism of regulatory transition between development and aging, we measured mRNA and microRNA expression patterns in brains (superior frontal gyrus) of humans and rhesus macaques over the entire species’ lifespan. more...
Organism:	Macaca mulatta; Homo sapiens
Type:		Expression profiling by array
Platform: GPL6244 51 Samples
FTP download: GEO (CEL, TXT, XLS) ftp://ftp.ncbi.nlm.nih.gov/geo/series/GSE17nnn/GSE17757/
Series		Accession: GSE17757	ID: 200017757

47. Comparison of post-mortem tissue from brain BA10 region between schizophrenic and control patients.
(Submitter supplied) Analysis of gene expression in two large schizophrenia cohorts identifies multiple changes associated with nerve terminal function. Schizophrenia is a severe psychiatric disorder with a world-wide prevalence of 1%. The pathophysiology of the illness is not understood, but is thought to have a strong genetic component with some environmental influences on aetiology. To gain further insight into disease mechanism, we used microarray technology to determine the expression of over 30 000 mRNA transcripts in post-mortem tissue from a brain region associated with the pathophysiology of the disease (Brodmann area 10: anterior prefrontal cortex) in 28 schizophrenic and 23 control patients.
Organism:	Homo sapiens
Type:		Expression profiling by array
Dataset: GDS4523 Platform: GPL570 51 Samples
FTP download: GEO (CEL) ftp://ftp.ncbi.nlm.nih.gov/geo/series/GSE17nnn/GSE17612/
Series		Accession: GSE17612	ID: 200017612

48. Profiling Transcriptomic Alterations in Postmortem Prefrontal Cortex Tissues of Individuals with Alcohol Use Disorders
(Submitter supplied) Analysis of transcriptiomic alternations related with alcohol use disorders (AUDs). The hypothesis is that chronic alcohol consumption might alter genome-wide gene expression patterns. The results suggest that differential gene expression in the prefrontal cortex is implicated in neuroadaptations to alcohol.
Organism:	Homo sapiens
Type:		Expression profiling by array
Platform: GPL10904 48 Samples
FTP download: GEO (TXT) ftp://ftp.ncbi.nlm.nih.gov/geo/series/GSE49nnn/GSE49376/
Series		Accession: GSE49376	ID: 200049376

49. Transcriptomic effects of caloric restriction reflected in primate difference
(Submitter supplied) Caloric restriction (CR) can delay morbidity and mortality in a broad range of species, including mice and macaques. Mutations and chemical agents, such as resveratrol or rapamycin that partly mimic the CR effect, can similarly increase survival or extend lifespan. In humans, however, the effects of CR or other life-extending agents have not been investigated systematically. Humans already display lower mortality and greater maximal lifespan compared to closely related species, including chimpanzees and macaques. more...
Organism:	Macaca mulatta; Homo sapiens; Mus musculus; Pan troglodytes
Type:		Expression profiling by array
Platforms: GPL6246 GPL5175 48 Samples
FTP download: GEO (CEL) ftp://ftp.ncbi.nlm.nih.gov/geo/series/GSE44nnn/GSE44147/
Series		Accession: GSE44147	ID: 200044147

50. MicroRNA expression and regulation in human, chimpanzee, and macaque brain
(Submitter supplied) Among other factors, changes in gene expression on the human evolutionary lineage have been suggested to play an important role in the establishment of human-specific phenotypes. However, the molecular mechanisms underlying these expression changes are largely unknown. Here, we have explored the role of microRNA (miRNA) in the regulation of gene expression divergence between adult humans, chimpanzees and rhesus macaques, in two brain regions: prefrontal cortex and cerebellum. more...
Organism:	Pan troglodytes; Macaca mulatta; Homo sapiens
Type:		Expression profiling by array; Non-coding RNA profiling by array; Non-coding RNA profiling by high throughput sequencing
6 related Platforms 46 Samples
FTP download: GEO (CEL, TXT) ftp://ftp.ncbi.nlm.nih.gov/geo/series/GSE26nnn/GSE26545/
Series		Accession: GSE26545	ID: 200026545

51. Gene expression in the human prefrontal cortex during postnatal development
(Submitter supplied) Fresh frozen post mortem prefrontal cortex tissue (Brodman area 46) was obtained from 44 individuals varying in age from 0 to 49 years. RNA was extracted from these samples and hybridized to HG133plus2.0 GeneChips. The data was used to examine patterns of gene expression over the course of human postnatal developmental and ageing.  PMI - postmortem interval, DLPFC - dorsolateral prefrontal cortex
Organism:	Homo sapiens
Type:		Expression profiling by array
Platform: GPL570 44 Samples
FTP download: GEO (CEL) ftp://ftp.ncbi.nlm.nih.gov/geo/series/GSE13nnn/GSE13564/
Series		Accession: GSE13564	ID: 200013564

52. Transcriptome profiling of prefrontal cortex (BA9) in neurodegenerative dementia using Affymetrix Human Transcriptome Array 2.0
(Submitter supplied) We investigated transcriptome alterations in the prefrontal cortex (BA) of patients with Alzheimer’s disease (AD), dementia with Lewy bodies (DLB) and Parkinson's disease dementia (PDD) when compared with elderly controls without neurological or psychiatric diseases (CTRL) using Affymetrix Human Transcriptome Array 2.0 All brain samples were provided by the Brains for Dementia Research (BDR), UK. All cases were prospectively assessed by experienced clinicians using validated clinical rating instruments. more...
Organism:	Homo sapiens
Type:		Expression profiling by array
Platform: GPL17585 42 Samples
FTP download: GEO (CEL) ftp://ftp.ncbi.nlm.nih.gov/geo/series/GSE150nnn/GSE150696/
Series		Accession: GSE150696	ID: 200150696

53. Repression of Human and Mouse Brain Inflammaging Transcriptome by Broad Gene Body Histone Hyperacetylation (RNA-seq)
(Submitter supplied) RNA-seq data of human prefrontal cortex and mouse brain during aging and/or SAHA treatment were generated.
Organism:	Homo sapiens; Mus musculus
Type:		Expression profiling by high throughput sequencing
Platforms: GPL13112 GPL11154 42 Samples
FTP download: GEO (TXT) ftp://ftp.ncbi.nlm.nih.gov/geo/series/GSE106nnn/GSE106669/
SRA Run Selector: https://www.ncbi.nlm.nih.gov/Traces/study/?acc=PRJNA417573
Series		Accession: GSE106669	ID: 200106669

54. Comparison of post-mortem tissue from Brodman Brain BA22 region between schizophrenic and control patients
(Submitter supplied) Transcriptional analysis of the superior temporal cortex (BA22) in schizophrenia: Pathway insight into disease pathology and drug development Schizophrenia is a highly debilitating psychiatric disorder which is known to have heritable genetic and environmental components. To gain some insight into the mechanisms underpinning both positive and negative symptoms of the disease, we determined the genome wide expression of mRNA transcripts in post-mortem tissue from the superior temporal cortex (Brodmann Area 22, BA22) in schizophrenic and control patients. more...
Organism:	Homo sapiens
Type:		Expression profiling by array
Dataset: GDS4522 Platform: GPL570 42 Samples
FTP download: GEO (CEL) ftp://ftp.ncbi.nlm.nih.gov/geo/series/GSE21nnn/GSE21935/
Series		Accession: GSE21935	ID: 200021935

55. Brain Gene Expression Pattern of Individuals with Dual Diagnosis who died by suicide
(Submitter supplied) We compared the gene expression profile in the dorsolateral prefrontal area of subjects with dual diagnosis who died by suicide to the transcriptome of subjects with mood disorders and subjects with substance use disorder who died by suicide.
Organism:	Homo sapiens
Type:		Expression profiling by array
Platform: GPL10558 38 Samples
FTP download: GEO (IDAT) ftp://ftp.ncbi.nlm.nih.gov/geo/series/GSE125nnn/GSE125681/
Series		Accession: GSE125681	ID: 200125681

56. Concordance of inflammatory markers in lymphocytes and prefrontal cortex in schizophrenia
(Submitter supplied) Schizophrenia is a severe neuropsychiatric disorder associated with a wide array of transcriptomic and neurobiochemical changes. Genome-wide transcriptomic profiling conducted in postmortem brain have provided novel insights into the pathophysiology of this disorder, and identified biological processes including immune/inflammatory-related responses, metabolic, endocrine and synaptic function. However, few studies have investigated whether similar changes could be found in peripheral tissue. more...
Organism:	Homo sapiens
Type:		Expression profiling by high throughput sequencing
Platform: GPL16791 37 Samples
FTP download: GEO (XLSX) ftp://ftp.ncbi.nlm.nih.gov/geo/series/GSE165nnn/GSE165604/
SRA Run Selector: https://www.ncbi.nlm.nih.gov/Traces/study/?acc=PRJNA695206
Series		Accession: GSE165604	ID: 200165604

57. Hemispheric asymmetry in the human brain and in Parkinson’s disease is linked to divergent epigenetic patterns in neurons [RNA-seq]
(Submitter supplied) Background Hemispheric asymmetry in neuronal processes is a fundamental feature of the human brain and drives symptom lateralization in Parkinson's disease (PD), but its molecular determinants are unknown. Here, we identify divergent epigenetic patterns involved in hemispheric asymmetry by profiling DNA methylation in isolated prefrontal cortex neurons from control and PD brain hemispheres. DNA methylation is fine-mapped at enhancers and promoters, genome-wide, by targeted bisulfite sequencing in two independent sample cohorts. more...
Organism:	Homo sapiens
Type:		Expression profiling by high throughput sequencing
Platform: GPL18573 36 Samples
FTP download: GEO (TAB) ftp://ftp.ncbi.nlm.nih.gov/geo/series/GSE135nnn/GSE135036/
SRA Run Selector: https://www.ncbi.nlm.nih.gov/Traces/study/?acc=PRJNA557205
Series		Accession: GSE135036	ID: 200135036

58. Gene Network Dysregulation in Dorsolateral Prefrontal Cortex Neurons of Humans with Cocaine Use Disorder
(Submitter supplied) total RNA-sequencing on dlPFC neurons from 19 human cocaine addicts (1-19) and 17 healthy controls (21-40)
Organism:	Homo sapiens
Type:		Expression profiling by high throughput sequencing
Platform: GPL11154 36 Samples
FTP download: GEO (XLSX) ftp://ftp.ncbi.nlm.nih.gov/geo/series/GSE99nnn/GSE99349/
SRA Run Selector: https://www.ncbi.nlm.nih.gov/Traces/study/?acc=PRJNA388174
Series		Accession: GSE99349	ID: 200099349

59. Network preservation reveals shared and unique biological processes associated with chronic alcohol abuse in the NAc and PFC [mRNA]
(Submitter supplied) Chronic alcohol abuse has been linked to the disruption of executive function and allostatic conditioning of reward response dysregulation in the mesocorticolimbic pathway (MCL). Here, we analyzed genome-wide mRNA and miRNA expression from matched cases with alcohol dependence (AD) and controls (n=35) via gene network analysis to identify unique and shared biological processes dysregulated in the prefrontal cortex (PFC) and nucleus accumbens (NAc). more...
Organism:	Homo sapiens
Type:		Expression profiling by array
Platform: GPL571 35 Samples
FTP download: GEO (CEL) ftp://ftp.ncbi.nlm.nih.gov/geo/series/GSE161nnn/GSE161986/
Series		Accession: GSE161986	ID: 200161986

60. Gene expression in human post-mortem brain samples from schizophrenic subjects and matched controls
(Submitter supplied) This laboratory focuses on the molecular mechanisms of neuropsychiatric and neurodegenerative disorders
Organism:	Mus musculus; Homo sapiens
Type:		Expression profiling by array
Platform: GPL11094 32 Samples
FTP download: GEO (CEL) ftp://ftp.ncbi.nlm.nih.gov/geo/series/GSE27nnn/GSE27356/
Series		Accession: GSE27356	ID: 200027356

61. Laser capture microdissection of endothelial and neuronal cells from human dorsolateral prefrontal cortex
(Submitter supplied) We used laser capture microdissection to isolate both microvascular endothelial cells and neurons from post mortem brain tissue from patients with schizophrenia and bipolar disorder and healthy controls. RNA was isolated from these cell populations, amplified, and analysed using Affymetrix HG133plus2.0 GeneChips. In the first instance, we used the dataset to compare the neuronal and endothelial data, in order to demonstrate that the predicted differences between cell types could be detected using this methodology. more...
Organism:	Homo sapiens
Type:		Expression profiling by array
Dataset: GDS3502 Platform: GPL570 32 Samples
FTP download: GEO (CEL) ftp://ftp.ncbi.nlm.nih.gov/geo/series/GSE12nnn/GSE12679/
Series		Accession: GSE12679	ID: 200012679

62. Dysregulation of brain and choroid plexus cell types in severe COVID-19
(Submitter supplied) Though SARS-CoV-2 primarily targets the respiratory system, patients and survivors can suffer neurological symptoms. Yet, an unbiased understanding of the cellular and molecular processes affected in the brains of COVID-19 patients is still missing. Here, we profile 65,309 single-nucleus transcriptomes from 30 frontal cortex and choroid plexus samples across 14 control (including 1 terminal influenza) and 8 COVID-19 patients. more...
Organism:	Homo sapiens
Type:		Expression profiling by high throughput sequencing
Platform: GPL24676 30 Samples
FTP download: GEO (MTX, TSV) ftp://ftp.ncbi.nlm.nih.gov/geo/series/GSE159nnn/GSE159812/
SRA Run Selector: https://www.ncbi.nlm.nih.gov/Traces/study/?acc=PRJNA670674
Series		Accession: GSE159812	ID: 200159812

63. Aberrant transcriptomes and DNA methylomes define pathways that drive pathogenesis and loss of brain laterality/asymmetry in schizophrenia and bipolar disorder [Affymetrix]
(Submitter supplied) Although the loss or reversal of brain laterality is one of the most consistent modalities in schizophrenia (SCZ) and bipolar disorder (BD), its molecular basis remains elusive. Our limited previous studies indicated that epigenetic modifications are key to the asymmetric transcriptomes of brain hemispheres. We used whole-genome expression microarrays to profile post-mortem brain samples from subjects with SCZ, psychotic BD [BD(+)] or non-psychotic BD [BD(-)], or matched controls (n=10/group, corresponding to different brain hemispheres) and performed whole-genome DNA methylation (DNAM) profiling of the same samples (n=3-4/group) to identify pathways associated with SCZ or BD(+) and genes/sites susceptible to epigenetic regulation. more...
Organism:	Homo sapiens
Type:		Expression profiling by array
Platform: GPL16311 30 Samples
FTP download: GEO (CEL) ftp://ftp.ncbi.nlm.nih.gov/geo/series/GSE120nnn/GSE120340/
Series		Accession: GSE120340	ID: 200120340

64. Transcriptional analysis of prefrontal area 9 in Parkinson's disease
(Submitter supplied) Post mortem tissue was dissected from two groups of age and gender matched groups of Parkinson and Control subjects
Organism:	Homo sapiens
Type:		Expression profiling by array
Platform: GPL96 29 Samples
FTP download: GEO (CEL, CHP) ftp://ftp.ncbi.nlm.nih.gov/geo/series/GSE20nnn/GSE20168/
Series		Accession: GSE20168	ID: 200020168

65. perro-affy-human-186940
(Submitter supplied) Our laboratory has developed the first mouse model overexpressing a RNA-binding protein, the ELAV-like protein HuD, in the CNS under the control of the CaMKinII alpha promoter. Initial behavioral characterization of the mice revealed that they had significant learning deficits together with abnormalities in prepulse inhibition (PPI). At the molecular level, we found that the expression of the growth-associated protein GAP-43, one of the targets of HuD, was increased in the hippocampus of HuD transgenic mice. more...
Organism:	Homo sapiens
Type:		Expression profiling by array
Dataset: GDS1917 Platform: GPL570 28 Samples
FTP download: GEO (CEL) ftp://ftp.ncbi.nlm.nih.gov/geo/series/GSE4nnn/GSE4036/
Series		Accession: GSE4036	ID: 200004036

66. Genome-wide analysis of mRNA polyadenylation in prefrontal cortex of idiopathic ASD patients
(Submitter supplied) The aim of this work was to test the mRNA polyadenylation status in the brain of autistic patients. We performed poly(U)-chromatography on RNA purified from post-mortem prefrontal cortex tissue of control and idiopathic ASD cases.
Organism:	Homo sapiens
Type:		Expression profiling by array
Platform: GPL15207 27 Samples
FTP download: GEO (CEL) ftp://ftp.ncbi.nlm.nih.gov/geo/series/GSE113nnn/GSE113834/
Series		Accession: GSE113834	ID: 200113834

67. Gene expression data from primate postnatal brain development - superior frontal gyrus
(Submitter supplied) In development, timing is of the utmost importance, and the timing of various developmental processes are often changed during evolution. During human evolution sexual maturation has been delayed relative to other primates and this may have played a critical role for both the increase of human brain size and the rise of human-specific cognitive traits . We measured the timing of gene expression changes in the superior frontal gyrus region of the brains of humans, chimpanzees, and rhesus macaques throughout postnatal development. more...
Organism:	Pan troglodytes; Macaca mulatta; Homo sapiens
Type:		Expression profiling by array
Platform: GPL6244 27 Samples
FTP download: GEO (CEL, TSV) ftp://ftp.ncbi.nlm.nih.gov/geo/series/GSE15nnn/GSE15163/
Series		Accession: GSE15163	ID: 200015163

68. Epigenetic dysregulation of enhancers in neurons is associated with Alzheimer’s disease pathology and cognitive symptoms (RNA-Seq)
(Submitter supplied) Epigenetic control of enhancers alters neuron functions and may be involved in Alzheimer’s disease (AD). Here, we identify enhancers in neurons contributing to AD by comprehensive fine-mapping of DNA methylation at enhancers, genome-wide. We examine 1.2 million CpG and CpH sites in enhancers in prefrontal cortex neurons of individuals with no/mild, moderate, and severe AD pathology (n=101). We identify 1,224 differentially methylated enhancer regions; most of which are hypomethylated at CpH sites in AD neurons. more...
Organism:	Homo sapiens
Type:		Expression profiling by high throughput sequencing
Platform: GPL18573 25 Samples
FTP download: GEO (TXT) ftp://ftp.ncbi.nlm.nih.gov/geo/series/GSE110nnn/GSE110731/
SRA Run Selector: https://www.ncbi.nlm.nih.gov/Traces/study/?acc=PRJNA434441
Series		Accession: GSE110731	ID: 200110731

69. Evolution of neuronal and endothelial transcriptomes in primates
(Submitter supplied) Neurons and endothelial cells were identified by immunohistochemistry in human brains, isolated by laser-capture-microdissection and used to find genes preferentially expressed in the two cell types.  Keywords: cell type comparison
Organism:	Homo sapiens; Mus musculus
Type:		Expression profiling by array
Platforms: GPL570 GPL81 25 Samples
FTP download: GEO (CEL) ftp://ftp.ncbi.nlm.nih.gov/geo/series/GSE12nnn/GSE12293/
Series		Accession: GSE12293	ID: 200012293

70. Genome-wide analyses of human perisylvian cerebral cortical patterning
(Submitter supplied) Despite the well-established role of the frontal and posterior peri-sylvian cortices in many facets of human-cognitive specializations, including language, little is known about the developmental patterning of these regions in human brain. We performed a genome-wide analysis of human cerebral patterning during mid-gestation, a critical epoch in cortical regionalization. A total of 345 genes were identified as differentially expressed (DE) between superior temporal gyrus (STG) and the remaining cerebral cortex (CTX). more...
Organism:	Homo sapiens
Type:		Expression profiling by array
Platforms: GPL885 GPL96 25 Samples
FTP download: GEO (CEL, EXP, TXT) ftp://ftp.ncbi.nlm.nih.gov/geo/series/GSE9nnn/GSE9335/
Series		Accession: GSE9335	ID: 200009335

71. Single nuclei transcriptomic profiling of human astrocytes and oligodendrocytes in Alzheimer’s disease
(Submitter supplied) This SuperSeries is composed of the SubSeries listed below.
Organism:	Homo sapiens
Type:		Expression profiling by high throughput sequencing
Platform: GPL24676 24 Samples
FTP download: GEO (MTX, TSV) ftp://ftp.ncbi.nlm.nih.gov/geo/series/GSE167nnn/GSE167494/
Series		Accession: GSE167494	ID: 200167494

72. Glial profiling of human tauopathy brain demonstrates enrichment of astrocytic transcripts in tau-related frontotemporal degeneration
(Submitter supplied) To understand how glia may be altered in frontotemporal degeneration with tau pathology (FTD-tau), we used a NanoString glial profiling panel to measure 770 transcripts related to glial biology in human control (n = 8), Alzheimer’s disease (AD) (n = 8), and FTD-tau (n = 8) dorsolateral prefrontal cortex. Compared to control, 43 genes were upregulated and 86 genes were downregulated in the FTD-tau samples. more...
Organism:	Homo sapiens
Type:		Expression profiling by array
Platform: GPL31198 24 Samples
FTP download: GEO (RCC, TXT) ftp://ftp.ncbi.nlm.nih.gov/geo/series/GSE193nnn/GSE193391/
Series		Accession: GSE193391	ID: 200193391

73. Profiling of mRNA transcriptomic changes using RNA-seq in postmortem prefrontal cortex of subjects with alcohol use disorder
(Submitter supplied) Purpose: The goal of this study to examine mRNA transcriptomic changes in reward-related brain regions of subjects with alcohol use disorder. Methods: Total RNAs were extracted from postmortem prefrontal cortex of 12 AUD and 12 control subjects. rRNA depletion RNA sequencing was performed and the sequence reads were processed using the bulk RNA-seq processing pipeline Pipeliner workflow (Federico et al. more...
Organism:	Homo sapiens
Type:		Expression profiling by high throughput sequencing
Platform: GPL16791 24 Samples
FTP download: GEO (TXT) ftp://ftp.ncbi.nlm.nih.gov/geo/series/GSE182nnn/GSE182173/
SRA Run Selector: https://www.ncbi.nlm.nih.gov/Traces/study/?acc=PRJNA755256
Series		Accession: GSE182173	ID: 200182173

74. Comparing Control and Schizophrenic hiPSC-derived NPCs
(Submitter supplied) Cell-based models of many neurological and psychiatric diseases, established by reprogramming patient somatic cells into human induced pluripotent stem cells (hiPSCs), have now been reported.  While numerous reports have demonstrated that neuronal cells differentiated from hiPSCs are electrophysiologically active mature neurons, the “age” of these cells relative to cells in the human brain remains unresolved. more...
Organism:	Homo sapiens
Type:		Expression profiling by array
Platform: GPL6244 24 Samples
FTP download: GEO (CEL, TXT) ftp://ftp.ncbi.nlm.nih.gov/geo/series/GSE40nnn/GSE40102/
Series		Accession: GSE40102	ID: 200040102

75. RNA-sequencing of the brain transcriptome implicates dysregulation of neuroplasticity, circadian rhythms, and GTPase binding in bipolar disorder
(Submitter supplied) See "Akula et al., Molecular Psychiatry in Press". RNA-sequencing (RNA-seq) is a powerful technique to investigate the complexity of gene expression in the human brain. We used RNA-seq to survey the brain transcriptome in high-quality post-mortem dorsolateral prefrontal cortex from 11 individuals diagnosed with bipolar disorder (BD) and from 11 age- and gender-matched controls.  Deep sequencing was performed, with over 350 million reads per specimen. more...
Organism:	Homo sapiens
Type:		Expression profiling by high throughput sequencing
Platforms: GPL15433 GPL9115 22 Samples
FTP download: GEO (TXT) ftp://ftp.ncbi.nlm.nih.gov/geo/series/GSE53nnn/GSE53239/
SRA Run Selector: https://www.ncbi.nlm.nih.gov/Traces/study/?acc=PRJNA231202
Series		Accession: GSE53239	ID: 200053239

76. Targeted, long-read RNA sequencing of non-coding genomic regions associated with neuropsychiatric functions.
(Submitter supplied) The human brain is one of the last frontiers of biomedical research. Genome-wide association studies (GWAS) have succeeded in identifying thousands of haplotype blocks associated with a range of neuropsychiatric traits, including disorders such as schizophrenia, Alzheimer’s and Parkinson’s disease. However, the majority of single nucleotide polymorphisms (SNPs) that mark these haplotype blocks fall within non-coding regions of the genome, hindering their functional validation. more...
Organism:	Homo sapiens
Type:		Expression profiling by high throughput sequencing
Platforms: GPL16791 GPL26167 20 Samples
FTP download: GEO (BED, GTF, TXT) ftp://ftp.ncbi.nlm.nih.gov/geo/series/GSE118nnn/GSE118158/
SRA Run Selector: https://www.ncbi.nlm.nih.gov/Traces/study/?acc=PRJNA484721
Series		Accession: GSE118158	ID: 200118158

77. Molecular signatures underlying neurofibrillary tangle susceptibility in Alzheimer’s disease
(Submitter supplied) Aggregation of hyperphosphorylated tau in neurofibrillary tangles (NFTs) is closely associated with neuronal death and the progression of cognitive deficits in Alzheimer’s disease (AD). Pathological tau appears to contribute to disease progression by disrupting axonal transport and synapse function; however, our understanding of the molecular alterations associated with tau aggregates and their cellular specificity remains limited. more...
Organism:	Homo sapiens
Type:		Expression profiling by high throughput sequencing
Platform: GPL24676 19 Samples
FTP download: GEO (CSV, H5) ftp://ftp.ncbi.nlm.nih.gov/geo/series/GSE129nnn/GSE129308/
SRA Run Selector: https://www.ncbi.nlm.nih.gov/Traces/study/?acc=PRJNA530780
Series		Accession: GSE129308	ID: 200129308

78. Molecular profiles of pyramidal cell dysfunction in the superior temporal cortex in schizophrenia
(Submitter supplied) Gray matter volume in the cerebral cortex has been consistently found to be decreased in patients with schizophrenia. The superior temporal gyrus (STG) is one of the cortical regions that exhibit the most pronounced volumetric reduction. This reduction is generally thought to reflect, at least in part, decreased number of synapses; the majority of these synapses are believed to be furnished by glutamatergic axon terminals onto the dendritic spines on pyramidal neurons. more...
Organism:	Homo sapiens
Type:		Expression profiling by array
Platform: GPL1352 18 Samples
FTP download: GEO (CEL, CHP) ftp://ftp.ncbi.nlm.nih.gov/geo/series/GSE37nnn/GSE37981/
Series		Accession: GSE37981	ID: 200037981

79. Single nuclei transcriptomic profiling of human astrocytes and oligodendrocytes in Alzheimer’s disease I
(Submitter supplied) Single nuclei RNA sequencing (snRNA-seq) dataset characterizing astrocytes and oligodendrocytes isolated from human postmortem APOE e2/3 Alzheimer's disease (AD) and aged-matched non-symptomatic (NS) brains.
Organism:	Homo sapiens
Type:		Expression profiling by high throughput sequencing
Platform: GPL24676 17 Samples
FTP download: GEO (MTX, TSV) ftp://ftp.ncbi.nlm.nih.gov/geo/series/GSE167nnn/GSE167490/
SRA Run Selector: https://www.ncbi.nlm.nih.gov/Traces/study/?acc=PRJNA704815
Series		Accession: GSE167490	ID: 200167490

80. Genome-Scale Transcriptional Regulatory Network Models of Psychiatric and Neurodegenerative Disorders
(Submitter supplied) Transcriptional regulatory changes in the developing and adult brain are prominent features of brain diseases, but the involvement of specific transcription factors (TFs) remains poorly understood. We integrated brain-specific DNase footprinting and TF-gene co-expression to reconstruct a transcriptional regulatory network (TRN) model for the human brain. We identified key regulator TFs whose predicted target genes were enriched for differentially expressed genes in the prefrontal cortex of individuals with psychiatric and neurodegenerative diseases. more...
Organism:	Homo sapiens
Type:		Expression profiling by array
Platform: GPL17077 16 Samples
FTP download: GEO (TXT) ftp://ftp.ncbi.nlm.nih.gov/geo/series/GSE102nnn/GSE102122/
Series		Accession: GSE102122	ID: 200102122

81. Expression profiling of human adult postmortem brain tissue from Down syndrome and healthy control subjects
(Submitter supplied) Down syndrome (DS) is the result of trisomy chromosome 21 but the mechanisms by which the genotype leads to the characteristic disease phenotype are unclear. We performed a microarray study using human adult brain tissue (dorsolateral prefrontal cortex) from DS subjects and healthy controls to characterise for the first time the human adult Down syndrome brain Keywords: disease state analysis
Organism:	Homo sapiens
Type:		Expression profiling by array
Dataset: GDS2941 Platform: GPL96 15 Samples
FTP download: GEO (CEL) ftp://ftp.ncbi.nlm.nih.gov/geo/series/GSE5nnn/GSE5390/
Series		Accession: GSE5390	ID: 200005390

82. Single-nuclei isoform RNA sequencing reveals combination patterns of transcript elements across human brain cell types
(Submitter supplied) Single-nuclei RNA-Seq is widely employed to investigate cell types, especially of human brain and frozen samples. In contrast to single-cell approaches, many single-nuclei reads are purely intronic. Here, using microfluidics, PCR-based artifact removal, target enrichment, and long-read sequencing, we developed single-nuclei isoform RNA-sequencing (‘SnISOr-Seq’), and applied it to human adult frontal cortex. more...
Organism:	Mus musculus; Homo sapiens
Type:		Expression profiling by high throughput sequencing
5 related Platforms 12 Samples
FTP download: GEO (GFF, MTX, TAB, TSV) ftp://ftp.ncbi.nlm.nih.gov/geo/series/GSE178nnn/GSE178175/
SRA Run Selector: https://www.ncbi.nlm.nih.gov/Traces/study/?acc=PRJNA737508
Series		Accession: GSE178175	ID: 200178175

83. mRNA and miRNA expression changes at the neurovascular unit reveal a role for autophagy and DNA binding in the cerebral microvasculature of the ageing brain  (human)
(Submitter supplied) Objective: To examine changes in the transcriptome of the neurovascular unit in ageing individuals
Organism:	Homo sapiens
Type:		Expression profiling by array
Platform: GPL570 12 Samples
FTP download: GEO (CEL) ftp://ftp.ncbi.nlm.nih.gov/geo/series/GSE127nnn/GSE127710/
Series		Accession: GSE127710	ID: 200127710

84. Next Generation Sequencing Facilitates Comparisons of Control and Schizophrenia-Patient derived hiPSC-derived NPCs
(Submitter supplied) Cell-based models of many neurological and psychiatric diseases, established by reprogramming patient somatic cells into human induced pluripotent stem cells (hiPSCs), have now been reported.  While numerous reports have demonstrated that neuronal cells differentiated from hiPSCs are electrophysiologically active mature neurons, the “age” of these cells relative to cells in the human brain remains unresolved. more...
Organism:	Homo sapiens
Type:		Expression profiling by high throughput sequencing
Platform: GPL16791 12 Samples
FTP download: GEO (TXT) ftp://ftp.ncbi.nlm.nih.gov/geo/series/GSE63nnn/GSE63738/
SRA Run Selector: https://www.ncbi.nlm.nih.gov/Traces/study/?acc=PRJNA268951
Series		Accession: GSE63738	ID: 200063738

85. Widespread splicing changes in human brain development and aging
(Submitter supplied) Differences in transcript splicing are well documented among tissues and between sexes in humans and other organisms. Further, splicing changes in selected genes have been reported in human and mouse brain development. Using high-throughput RNA sequencing we characterized the transcriptome-wide splicing changes occurring over the lifespan in the human brain. We found that in two brain regions, prefrontal cortex and cerebellum, as many as 37% of all expressed protein-coding multi-exon genes undergo significant splicing changes between birth and 98 years of age. more...
Organism:	Homo sapiens
Type:		Expression profiling by array
Platform: GPL11440 12 Samples
FTP download: GEO (CEL) ftp://ftp.ncbi.nlm.nih.gov/geo/series/GSE26nnn/GSE26586/
Series		Accession: GSE26586	ID: 200026586

86. The mRNA expression patterns in macaque brains from prenatal to neonatal
(Submitter supplied) We search for developmental changes specific to humans by examining gene expression profiles in the human, chimpanzee and rhesus macaque prefrontal and cerebellar cortex. In both brain regions, developmental patterns were more evolved in humans than in chimpanzees. To distinguish whether the human specific developmental pattern represent novel human-specific developmental patterns or a shift in the timing of the existing patterns, we measured mRNA expression patterns in macaque brains from prenatal to neonatal. more...
Organism:	Macaca mulatta; Homo sapiens
Type:		Expression profiling by array
Platform: GPL6244 12 Samples
FTP download: GEO (CEL) ftp://ftp.ncbi.nlm.nih.gov/geo/series/GSE29nnn/GSE29138/
Series		Accession: GSE29138	ID: 200029138

87. Aberrant transcriptomes and DNA methylomes define pathways that drive pathogenesis and loss of brain laterality/asymmetry in schizophrenia and bipolar disorder [Illumina]
(Submitter supplied) Although the loss or reversal of brain laterality is one of the most consistent modalities in schizophrenia (SCZ) and bipolar disorder (BD), its molecular basis remains elusive. Our limited previous studies indicated that epigenetic modifications are key to the asymmetric transcriptomes of brain hemispheres. We used whole-genome expression microarrays to profile post-mortem brain samples from subjects with SCZ, psychotic BD [BD(+)] or non-psychotic BD [BD(-)], or matched controls (n=10/group, corresponding to different brain hemispheres) and performed whole-genome DNA methylation (DNAM) profiling of the same samples (n=3-4/group) to identify pathways associated with SCZ or BD(+) and genes/sites susceptible to epigenetic regulation. more...
Organism:	Homo sapiens
Type:		Expression profiling by array
Platform: GPL8490 11 Samples
FTP download: GEO (IDAT) ftp://ftp.ncbi.nlm.nih.gov/geo/series/GSE120nnn/GSE120341/
Series		Accession: GSE120341	ID: 200120341

88. Next Generation Sequencing Facilitates Comparisons of Control and Schizophrenia-Patient derived hiPSC-derived neurons
(Submitter supplied) Cell-based models of many neurological and psychiatric diseases, established by reprogramming patient somatic cells into human induced pluripotent stem cells (hiPSCs), have now been reported.  While numerous reports have demonstrated that neuronal cells differentiated from hiPSCs are electrophysiologically active mature neurons, the “age” of these cells relative to cells in the human brain remains unresolved. more...
Organism:	Homo sapiens
Type:		Expression profiling by high throughput sequencing
Platform: GPL16791 11 Samples
FTP download: GEO (TXT) ftp://ftp.ncbi.nlm.nih.gov/geo/series/GSE63nnn/GSE63734/
SRA Run Selector: https://www.ncbi.nlm.nih.gov/Traces/study/?acc=PRJNA268948
Series		Accession: GSE63734	ID: 200063734

89. Single cell epigenomics reveals mechanisms of human cortical development
(Submitter supplied) During mammalian development, chromatin state differences coincide with cellular differentiation and reflect changes in the gene regulatory landscape. In the developing brain, cell fate specification and topographic identity play important roles in defining cell identity and confer selective vulnerabilities to neurodevelopmental disorders. To identify cell type specific chromatin accessibility patterns in the developing human brain, we used a single cell assay for transposase accessibility by sequencing (scATAC-seq) in primary human forebrain tissue samples. more...
Organism:	Homo sapiens
Type:		Expression profiling by high throughput sequencing; Genome binding/occupancy profiling by high throughput sequencing
Platform: GPL24676 10 Samples
FTP download: GEO (CSV, MTX, RDS, SNAP, TSV) ftp://ftp.ncbi.nlm.nih.gov/geo/series/GSE163nnn/GSE163018/
SRA Run Selector: https://www.ncbi.nlm.nih.gov/Traces/study/?acc=PRJNA684335
Series		Accession: GSE163018	ID: 200163018

90. Gene expression data from adult postnatal cerebellum, prefrontal cortex and liver: comparison on tissue differences
(Submitter supplied) To understand the roles of molecules in functional differentiation among adult human tissues, we performed a systematic survey of mRNA, protein, and protein phosphorylation as well as miRNA expression, in three tissues: cerebellum, prefrontal cortex and liver. We found that tissues were clearly distinct from one another at all levels. Furthermore, our results showed that miRNA differently expressed between tissues have significant, but modest effect on expression of mRNA and somewhat stronger effect on expression of proteins among the tissues. more...
Organism:	Homo sapiens
Type:		Expression profiling by array
Platform: GPL5175 9 Samples
FTP download: GEO (CEL) ftp://ftp.ncbi.nlm.nih.gov/geo/series/GSE19nnn/GSE19504/
Series		Accession: GSE19504	ID: 200019504

91. Single cell transcriptome profiling of the human alcohol-dependent brain samples.
(Submitter supplied) Over 16,000 nuclei were isolated from human postmartum brain frozen prefrontal cortex  samples of alcoholic and control individuals. Libraries were prepared with 10X Genomics platform and sequenced using NovaSeq 6000.
Organism:	Homo sapiens
Type:		Expression profiling by high throughput sequencing
Platform: GPL24676 8 Samples
FTP download: GEO (MTX, TSV) ftp://ftp.ncbi.nlm.nih.gov/geo/series/GSE141nnn/GSE141552/
SRA Run Selector: https://www.ncbi.nlm.nih.gov/Traces/study/?acc=PRJNA593861
Series		Accession: GSE141552	ID: 200141552

92. Aberrant expression of lncRNAs in autistic brain
(Submitter supplied) To assess for the potential contribution of dysregulated long non-coding RNA expression in autism pathogenesis, we profiled lncRNAs and mRNAs from post mortem brain tissue from autism patients and age/sex matched controls
Organism:	Homo sapiens
Type:		Non-coding RNA profiling by array; Expression profiling by array
Platform: GPL15314 8 Samples
FTP download: GEO (TXT) ftp://ftp.ncbi.nlm.nih.gov/geo/series/GSE36nnn/GSE36315/
Series		Accession: GSE36315	ID: 200036315

93. Single nuclei transcriptomic profiling of human astrocytes and oligodendrocytes in Alzheimer’s disease II
(Submitter supplied) Single nuclei RNA sequencing (snRNA-seq) dataset characterizing nuclei from a putative astrocyte enrichment method: SOX9-positive sorting.
Organism:	Homo sapiens
Type:		Expression profiling by high throughput sequencing
Platform: GPL24676 7 Samples
FTP download: GEO (MTX, TSV) ftp://ftp.ncbi.nlm.nih.gov/geo/series/GSE167nnn/GSE167492/
SRA Run Selector: https://www.ncbi.nlm.nih.gov/Traces/study/?acc=PRJNA704816
Series		Accession: GSE167492	ID: 200167492

94. RNA sequencing of rhesus macaque prefrontal cortex and human embryonic stem cell after sevoflurane exposure
(Submitter supplied) This SuperSeries is composed of the SubSeries listed below.
Organism:	Macaca mulatta; Homo sapiens
Type:		Expression profiling by high throughput sequencing
Platforms: GPL20795 GPL24522 4 Samples
FTP download: GEO ftp://ftp.ncbi.nlm.nih.gov/geo/series/GSE148nnn/GSE148438/
Series		Accession: GSE148438	ID: 200148438

95. The prefrontal cortex microRNA expression in mice offspring caused by glyphosate exposure during pregnancy and lactation
(Submitter supplied) In this study, the miRNAs expression pattern in prefrontal cortex (PFC) of postnatal day (PND) 28 mice offsprings which were exposured by glyphosate-based herbicide (GBH) during pregnancy and lactation was investigated. MiRNA microarray had detected 55 upregulated and 19 downregulated miRNAs in the PND28 mice offsprings' PFC.
Organism:	Mus musculus; Rattus norvegicus; Homo sapiens
Type:		Expression profiling by array
Platform: GPL18058 2 Samples
FTP download: GEO (GPR) ftp://ftp.ncbi.nlm.nih.gov/geo/series/GSE100nnn/GSE100079/
Series		Accession: GSE100079	ID: 200100079
