## Supplementary Material for "Lysergic acid diethylamide reverses aging- and neurodegeneration-associated brain transcriptional programs"

| no age | in vitro | no human samples | <10 samples | LCM | wrong tissue | neurodegenerative disease | repeated data | targeted screen | single cell |
|---|---|---|---|---|---|---|---|---|---|
| GSE120341 | GSE102122 | GSE99757 | GSE15163 | GSE12293 | GSE127710 | GSE129308 | GSE113834 | GSE11546 | GSE178175 |
| GSE120340 | GSE40102 | GSE29138 | GSE159812 | GSE12679 | GSE146446 | GSE135036 | GSE106669 | GSE118158 | GSE81475 |
| GSE182173 | GSE63734 | GSE4036 | GSE26545 | | GSE165604 | GSE135037 | GSE161986 | GSE27356 | |
| GSE87610 | GSE63738 | GSE99757 | GSE26586 | | GSE22569 | GSE135751 | GSE17757 | | |
| GSE53239 | GSE163018 | | GSE100796 | | | GSE136010 | GSE22521 | | |
| GSE44771 | | | GSE30352 | | | GSE150696 | GSE5392 | | |
| GSE44770 | | | GSE44147 | | | GSE159940 | | | |
| GSE44768 | | | GSE49379 | | | GSE167490 | | | |
| | | | GSE58604 | | | GSE167494 | | | |
| | | | GSE9335 | | | GSE193391 | | | |
| | | | GSE47966 | | | GSE20168 | | | |
| | | | | | | GSE20295 | | | |
| | | | | | | GSE110731 | | | |
| | | | | | | GSE110732 | | | |
| | | | | | | GSE68719 | | | |
| | | | | | | GSE84422 | | | |

**Table S1.** Discarded aging datasets. LCM = laser capture microdissection.

| Dataset | Platform | Reference |
|---|---|---|
| **GSE101521** | RNA-seq | Pantazatos *et al.(Pantazatos et al. 2017)* |
| **GSE102556** | RNA-seq | Labonté *et al.(Labonté et al. 2017)* |
| **GSE102741** | RNA-seq | Wright *et al.(Wright et al. 2017)* |
| **GSE106670 (GSE106669)** | RNA-seq | Cheng *et al.(Cheng et al. 2018)* |
| **GSE113842 (GSE113834)** | Affymetrix Human Gene Expression Array | Parras *et al.(Parras et al. 2018)* |
| **GSE11512** | Affymetrix GeneChip Human Genome U133 Plus 2.0 Array | Somel *et al.(Somel et al. 2009)* |
| **GSE125681** | Illumina HumanHT-12 V4.0 expression beadchip | Cabrera-Mendoza *et al.(Cabrera-Mendoza et al. 2020)* |
| **GSE13564** | Affymetrix Human Genome U133 Plus 2.0 Array | Breen *et al.(Breen et al. 2018)* |
| **GSE161986** | Affymetrix Human Genome U133A 2.0 Array | Vornholt *et al.(Vornholt et al. 2020)* |
| **GSE174409** | RNA-seq | Seney *et al.(Seney et al. 2021)* |
| **GSE17612** | Affymetrix Human Genome U133 Plus 2.0 Array | Maycox *et al.(Maycox et al. 2009)* |
| **GSE21138** | Affymetrix Human Genome U133 Plus 2.0 Array | Narauyan *et al.(Narayan et al. 2008)* |
| **GSE21935** | Affymetrix Human Genome U133 Plus 2.0 Array | Barnes *et al.(Barnes et al. 2011)* |
| **GSE22570** | Affymetrix Human Gene 1.0 ST Array [transcript (gene) version] | Dönertaş *et al.(Dönertaş et al. 2017)* |

| | | |
|---|---|---|
| **GSE37981** | Affymetrix Human X3P Array | Pietersen *et al.(Pietersen et al. 2014)* |
| **GSE49376** | Illumina HumanHT-12 V4.0 expression beadchip (gene symbol) | Xu *et al.(Xu et al. 2014)* |
| **GSE51264** | RNA-seq | He *et al.(He et al. 2014)* |
| **GSE5388** | Affymetrix Human Genome U133A Array | Ryan *et al.(Ryan et al. 2006)* |
| **GSE5390** | Affymetrix Human Genome U133A Array | Lockstone *et al.(Lockstone et al. 2007)* |
| **GSE59288** | RNA-seq | Liu *et al.(Liu et al. 2016)* |
| **GSE59630** | Affymetrix Human Exon 1.0 ST Array [transcript (gene) version] | Olmos-Serrano *et al.(Olmos-Serrano et al. 2016)* |
| **GSE60190** | Illumina HumanHT-12 V3.0 expression beadchip | Jaffe *et al.(Jaffe et al. 2014)* |
| **GSE64810** | RNA-seq | Labadorf *et al.(Labadorf et al. 2015)* |
| **GSE71620** | Affymetrix Human Gene 1.1 ST Array [transcript (gene) version] | French *et al.(French et al. 2017)* |
| **GSE92538** | Affymetrix Human Genome U133A Array | Hagenauer *et al.(Hagenauer et al. 2018)* |
| **GSE99349** | RNA-seq | Ribeiro *et al.(Ribeiro et al. 2017)* |
| **GTEx** | RNA-seq | |
| **GSE33000** | Rosetta/Merck Human 44k 1.1 microarray | Narayanan *et al.(Narayanan et al. 2014)* |
| **GSE30272** | Illumina Human 49K Oligo array | Colantuoni *et al.(Colantuoni et al. 2011)* |

**Table S2.** Aging datasets considered in our analysis.

| Category | # of samples |
| --- | --- |
| **Gender** | |
| Male | 2032 |
| Female | 1018 |
| **Region** | |
| DLPFC | 1335 |
| OFC | 480 |
| PFC | 783 |
| SFG | 84 |
| TC | 72 |
| CBC | 44 |
| VFC | 14 |
| Other | 274 |
| **Diagnosis** | |
| Healthy | 1863 |
| MDD | 237 |
| BD | 84 |
| SCZ | 80 |
| DS | 65 |
| AUD | 49 |
| ASD | 47 |
| OUD | 40 |
| HD | 177 |
| CUD | 18 |
| OCD | 9 |
| ED | 3 |
| AD | 310 |
| **Ethnicity** | |
| Caucasian | 918 |
| African American | 275 |
| **Alcohol** | |
| Yes | 86 |
| No | 173 |
| **Smoke** | |
| Yes | 213 |
| No | 310 |

**Table S3.** Samples' features of the ageing datasets considered in our analysis.

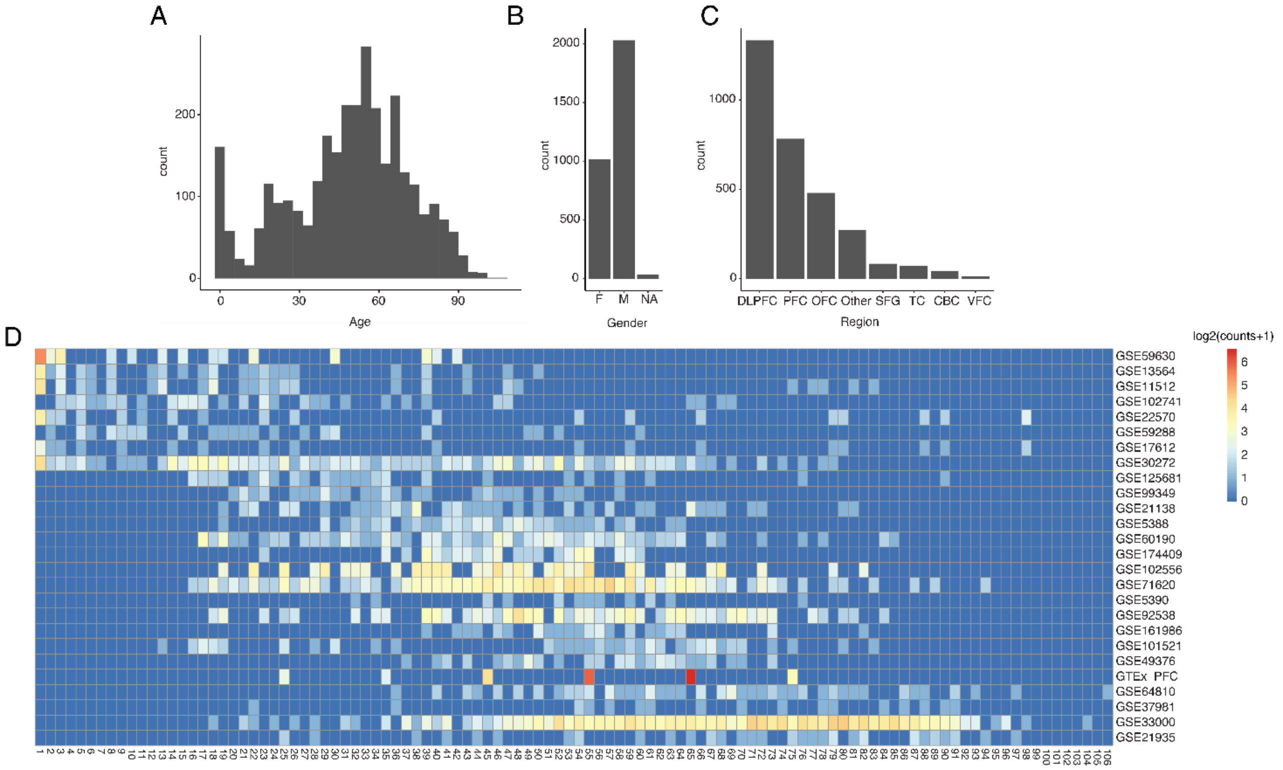


**Figure S1.** Annotations of samples included in the aging datasets collection. A) age distribution, B) gender, C) brain areas, D) number of samples by age (column) for each dataset (row). DLPFC=dorsolateral prefrontal cortex, PFC=prefrontal cortex, OFC=orbitofrontal cortex, SFG=superior frontal gyrus, TC=temporal cortex, CBC=cerebellar cortex, VFC=ventral frontal cortex.

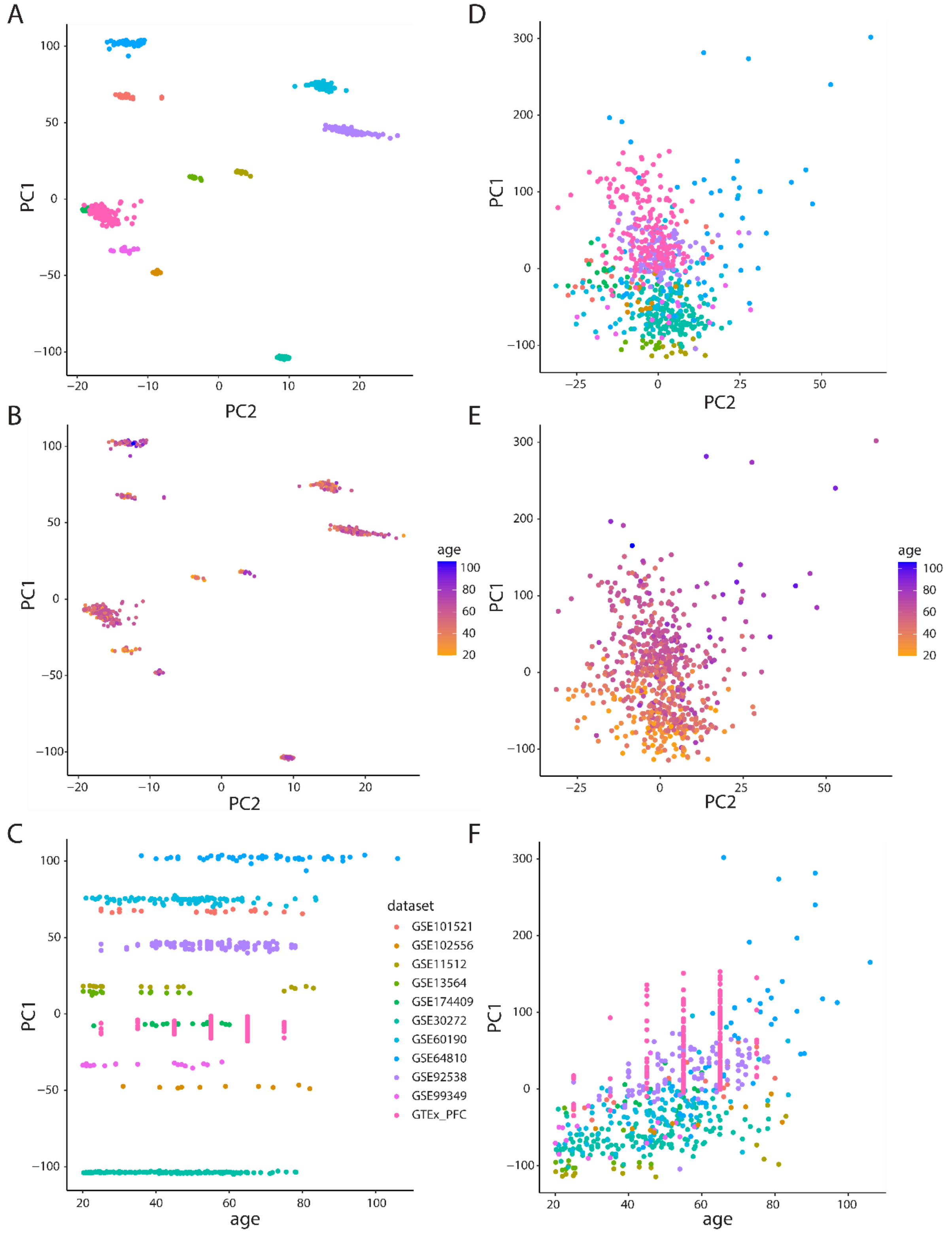


**Figure S2.** Integration of 11 transcriptomic DLPFC datasets. A) Samples of the 11 datasets plotted on the first two principal components (PC) with different colours indicating different datasets. A strong batch effect is observed, with little contribution to sample variability imputable to sample age (B, with sample points coloured according to subject age), as indicated by the lack of correlation between age and samples' position along PC1 (C). The same plots are displayed after batch correction (D-F), showing a more homogenous and less localised placement of the different datasets (D) and a positive and significant correlation (Pearson's R = 0.60, p-value < 2.2 x $10^{-16}$) between point position along PC1 and sample age (E, F).

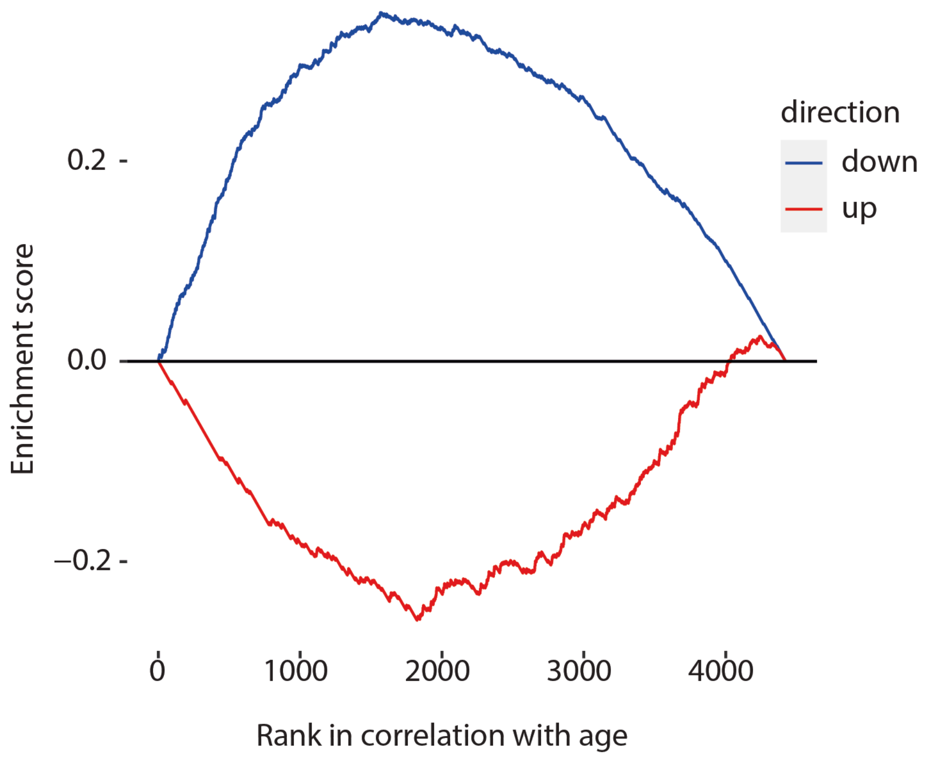


**Figure S3.** Gene Set Enrichment Analysis (GSEA) of genes ranked according to their correlation with age in the integrated dorsolateral prefrontal cortex (DLPF) datasets using differentially expressed genes (DEGs) upon chronic LSD administration (up-, or down-regulated, as indicated by the different colours) as query signatures. Up-regulated genes p-value = $1.8 \times 10^{-25}$, normalised enrichment score (NES) = -5.81, down-regulated genes p-value = $3.6 \times 10^{-44}$, NES = 5.54.

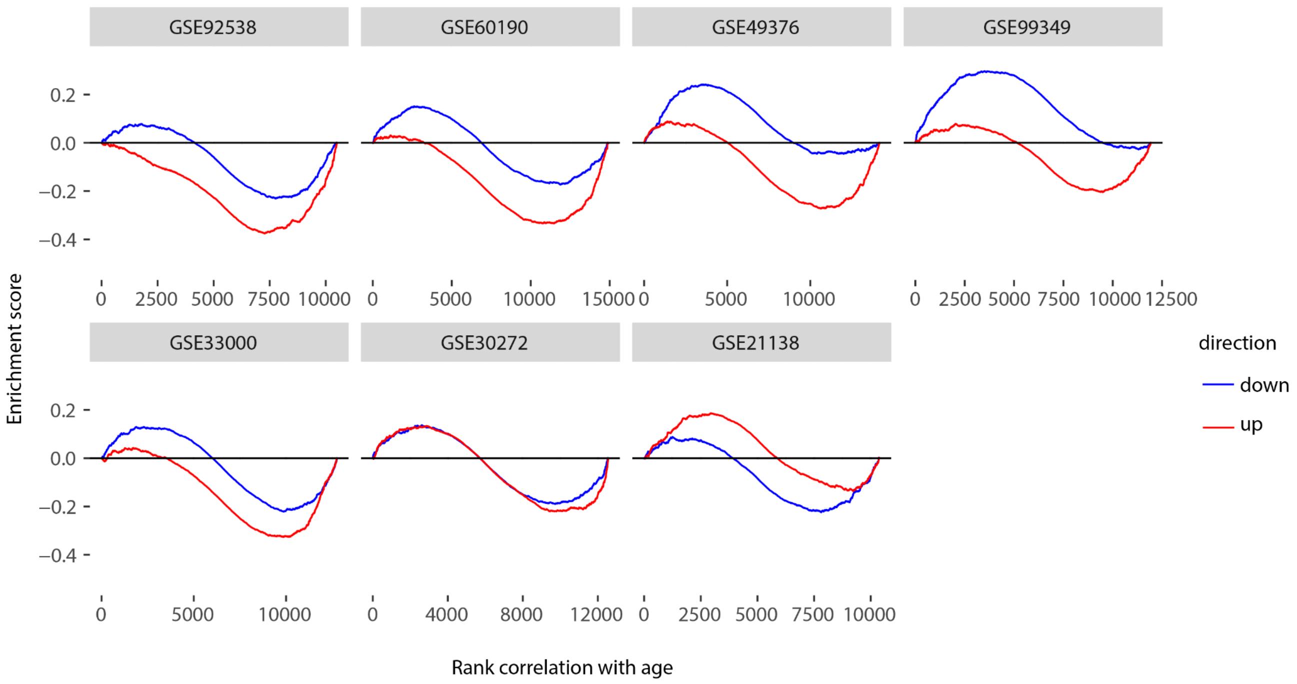


**Figure S4.** Gene Set Enrichment Analysis (GSEA) of genes ranked according to their correlation with age across prefrontal cortex datasets using differentially expressed genes (DEGs) upon chronic LSD administration (up-, or down-regulated, as indicated by the different colours) as query signatures. Shown are the seven datasets exhibiting the lowest absolute enrichment scores (out of fifteen total datasets).

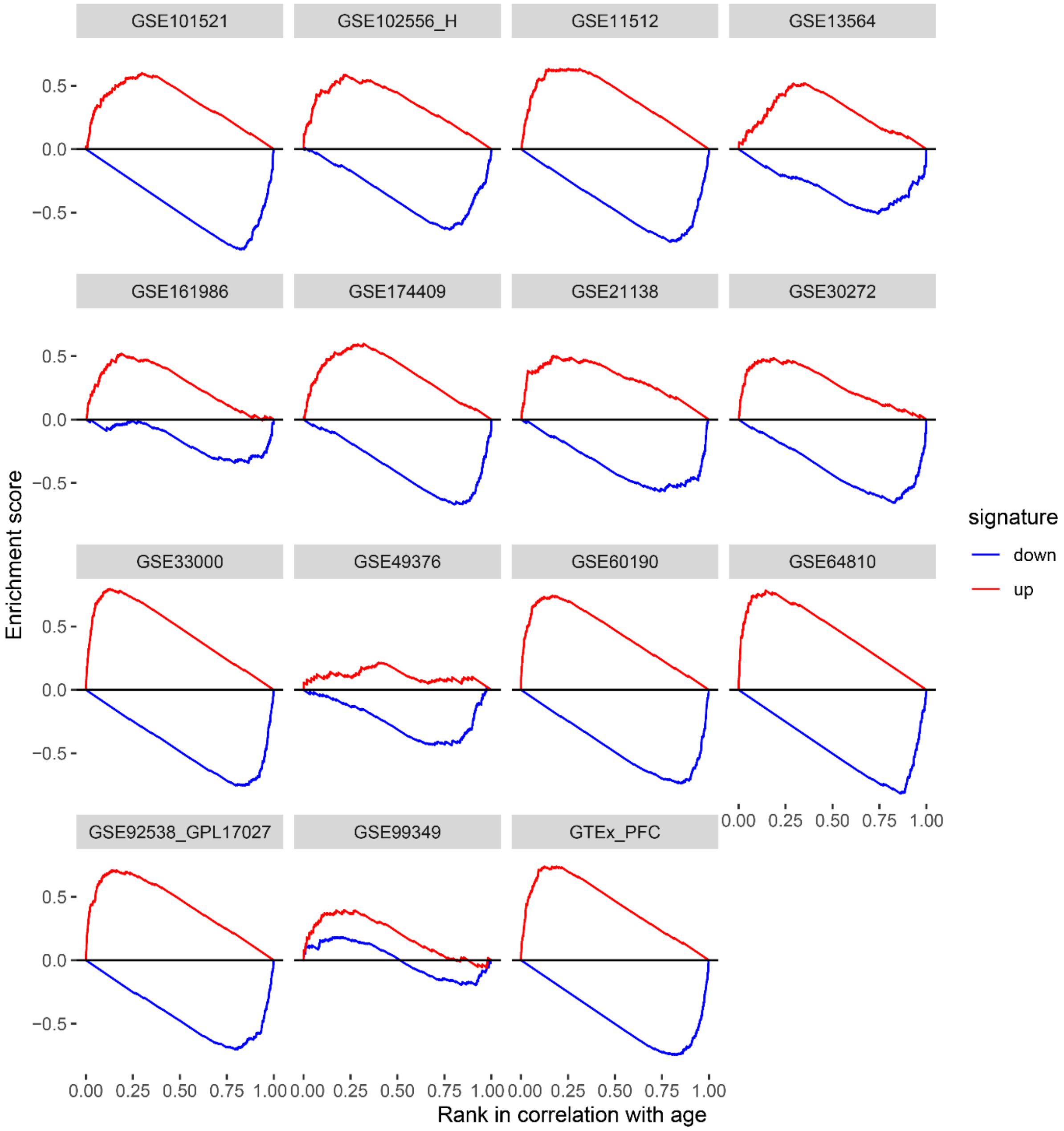


**Figure S5.** Gene Set Enrichment Analysis (GSEA) of genes ranked according to their correlation with age (one panel per dataset of dorsolateral/medial prefrontal cortex samples) using the aging-signature introduced in Donertas et al. (Dönertaş et al. 2018) as query signature.

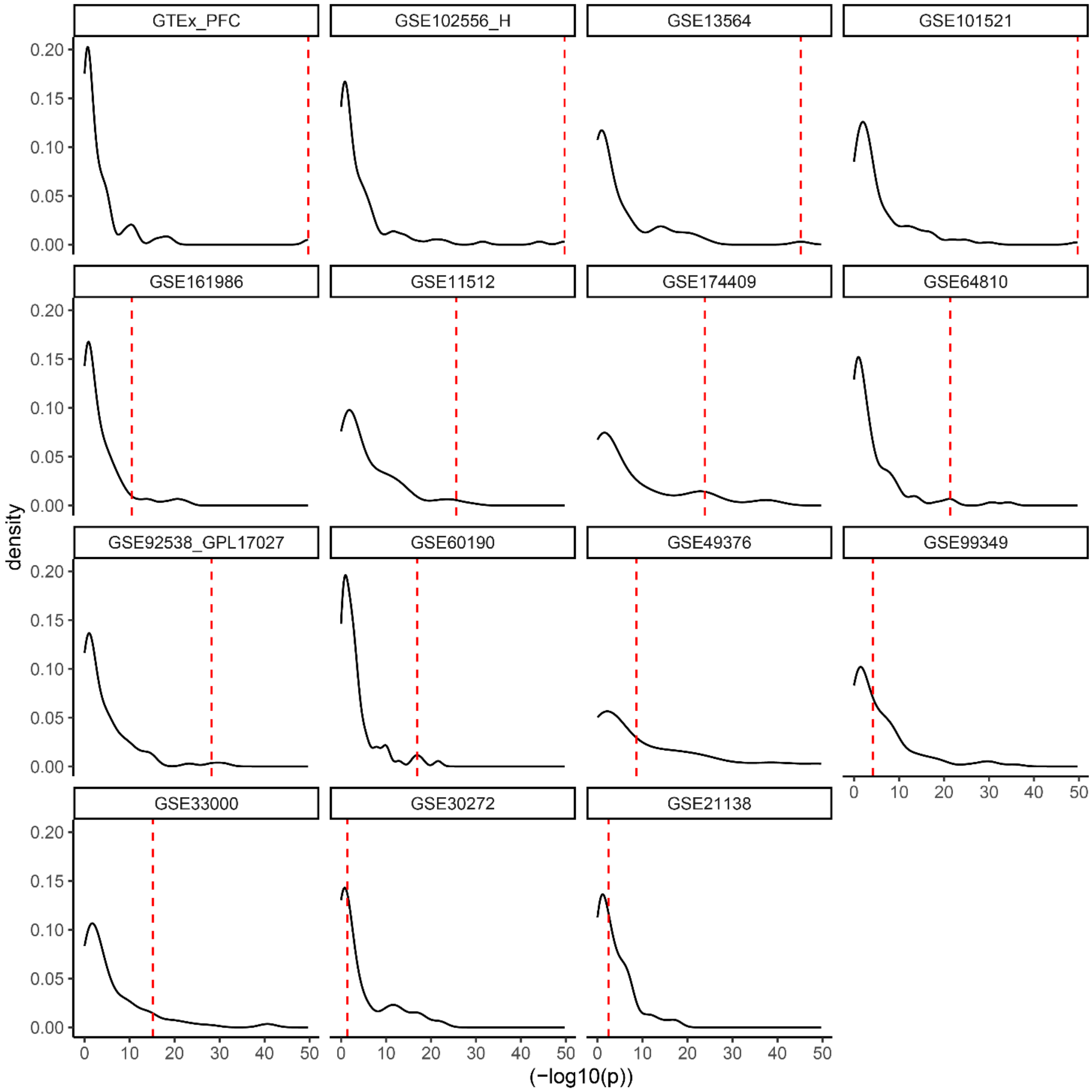


**Figure S6.** Distribution of GSEA p-values for age-reversal potential of connectivity map drugs screened in neural progenitor cells, using genes up-regulated by those drugs as query signature, for each aging dataset. The dashed red line indicates the corresponding p-value of chronic LSD.

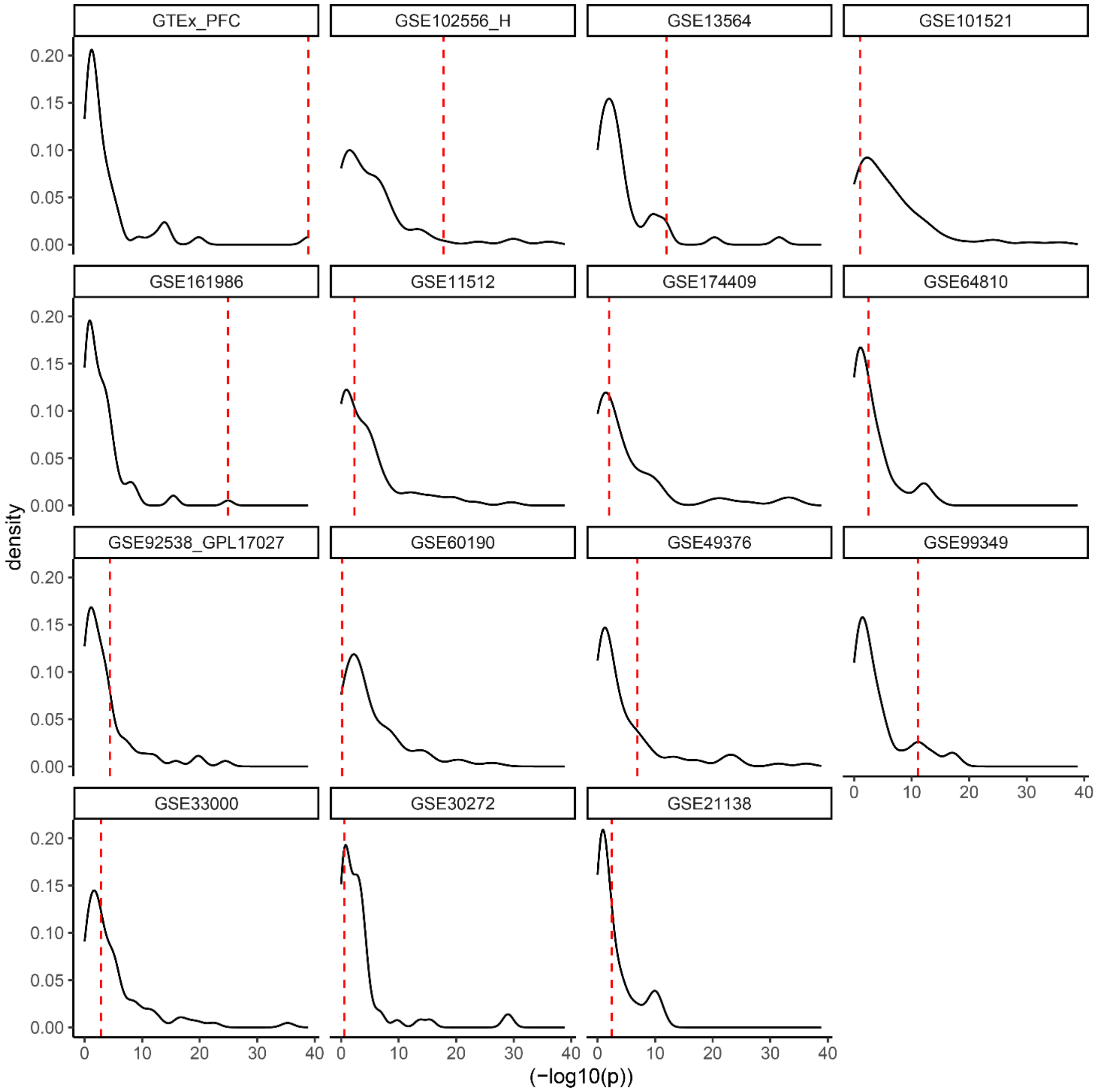


**Figure S7.** Distribution of GSEA p-values for age-reversal potential of connectivity map drugs screened in neural progenitor cells, using genes down-regulated by those drugs as query signature, for each aging dataset. The dashed red line indicates the corresponding p-value of chronic LSD.

| Dataset | Drug | Model | Treatment regime | Sampling time | Reference |
|---|---|---|---|---|---|
| Not deposited | MDMA | | | | Fernànded-Castillo *et al.(Fernàndez-Castillo et al. 2012)* |
| GSE161626 | DOI | M. musculus | Single administration | 24h, 48h, 7days | de la Fuente Revenga *et al.(de la Fuente Revenga et al. 2021)* |
| GSE81672 | Ketamine | | Single administration | | Bagot *et al.(Bagot et al. 2017)* |
| GSE26364 | Ketamine | M. musculus | Long term | | |
| GSE209859 | Psilocybin | M. musculus | Single administration | 3 hours, 4 weeks | Fadhunsi *et al.(Fadahunsi et al. 2022)* |
| GSE23728 (GSE14720) | DOI | R. norvegicus | Single administration | 2 hours | |
| GSE179378 | LSD | R. norvegicus | Single administration | 90 min | Nichols and Sander-Bush(Nichols & Sanders-Bush 2002) |
| GSE179379 | LSD | R. norvegicus | Chronic treatment | 3 weeks after withdrawal | Martin *et al.(Martin et al. 2014)* |
| Not deposited | DMT | | | | Kelley *et al.(Kelley et al. 2022)* |
| Not deposited | Harmaline | | | | Kelley *et al.(Kelley et al. 2022)* |
| Not deposited | Pharmahuasca | | | | Kelley *et al.(Kelley et al. 2022)* |

**Table S4.** Psychoplastogen datasets

| Dataset | Control | Label |
|---|---|---|
| GSE64607 | Positive | Exercise (28d) |
| GSE164798 | Positive | Exercise (30d) |
| GSE111273 | Positive | EE Young / EE Old |
| GSE72507 | Negative | Alcohol (VC) |
| GSE60676 | Negative | Alcohol (VC 2) |
| GSE28515 | Negative | Alcohol (IP injection) |

| | | |
|---|---|---|
| GSE75772 | Negative | rat aging |

**Table S5.** Positive and negative control datasets, with corresponding labels used in Figure 1EF.

**Table S6.** GSEA results for each combination of aging dataset, intervention dataset and direction of differential expression

**Table S7.** Meta-analyzed p-values and age-reversal p-value ratios for each intervention. Infinite values for age-reversal p-value ratios were set to 300.

| Dataset | Disease | Reference |
|---|---|---|
| GSE110731 | Alzheimer's Disease | Li *et al.(Li et al. 2019)* |
| GSE150696 | Alzheimer's Disease, dementia with Lewy bodies, Parkinson's disease dementia | Low *et al.(Low et al. 2021)* |
| GSE33000 | Alzheimer's and Huntington's Disease | Narayan *et al.(Narayanan et al. 2014)* |
| GSE44770 | Alzheimer's Disease | Zhang *et al.(Zhang et al. 2013)* |
| GSE53697 | Alzheimer's Disease | Scheckel *et al.(Scheckel et al. 2016)* |
| GSE84422 | Alzheimer's Disease | Wang *et al.(Wang et al. 2016)* |

**Table S8.** Dementia datasets

**Table S9.** GSEA results for all combinations of dementia dataset, intervention dataset and direction of differential expression

**Table S10.** Meta-analyzed p-values and dementia-reversal p-value ratios for each intervention.

**Table S11.** GSEA results for each combination of aging dataset, LSD concentration and direction of differential expression

**Table S12.** Meta-analyzed p-values and age-reversal p-value ratios for each LSD concentration.

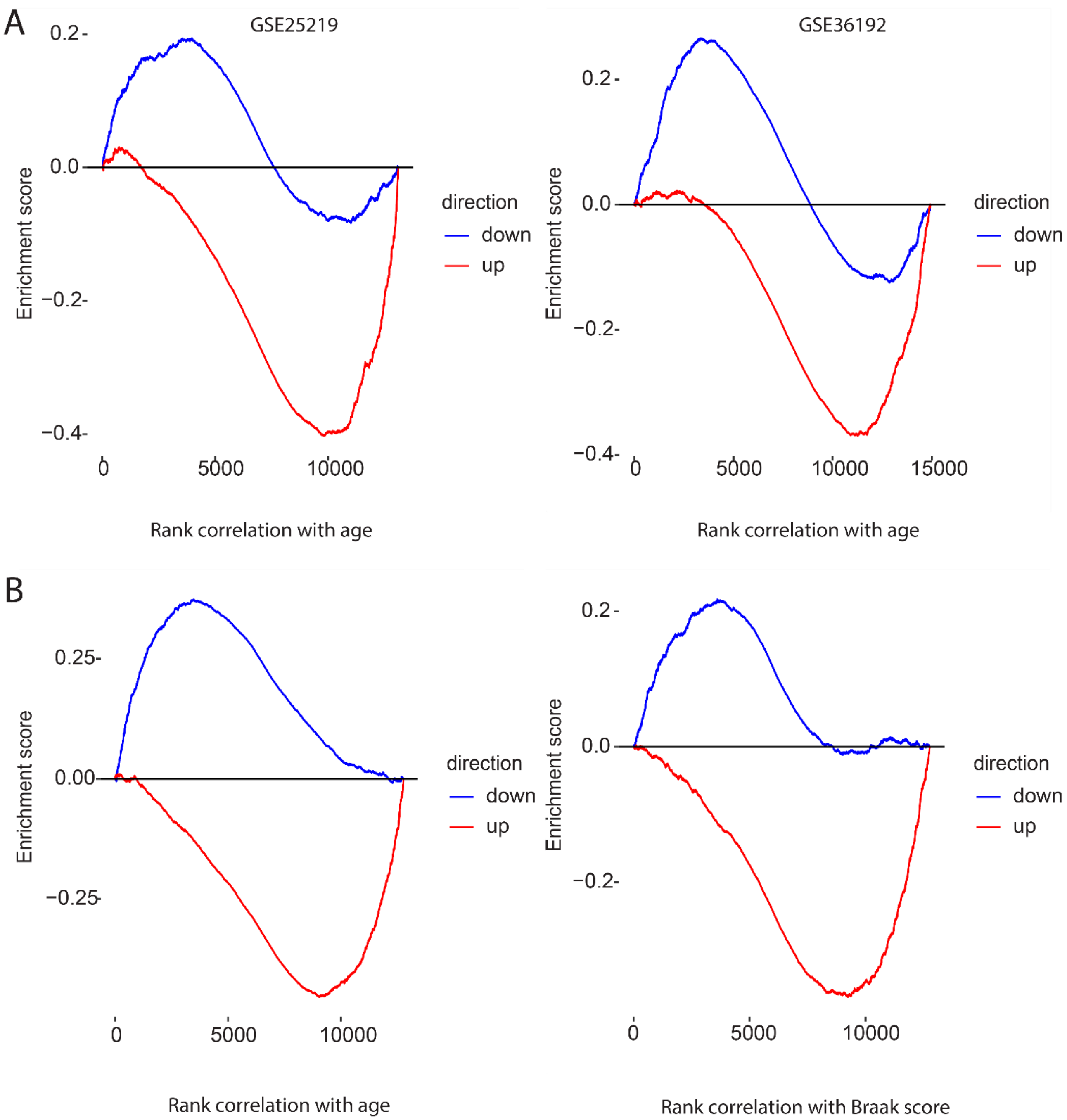


**Figure S8.** A) Gene set enrichment analysis of genes up- and down-regulated upon chronic LSD in rats (used as queries) along background ranked list of genes sorted based on the correlation between their expression and sampled subjects' age in two additional datasets not included in the main analysis, used as external validations**.** B) Gene set enrichment analysis of genes up- and down-regulated upon chronic LSD in rats for genes with high correlation with age according to the Mount Sinai Brain Bank (left) and for genes with high correlation with the Braak score of Alzheimer severity in the same dataset (right).

**Table S13.** Functional Enrichment for genes up-regulated by LSD and down-regulated upon aging

**Table S14.** Functional Enrichment for genes down-regulated by LSD and up-regulated upon aging

**Table S15.** Functional Enrichment for genes up-regulated by LSD and down-regulated in dementia

**Table S16.** Functional Enrichment for genes down-regulated by LSD and up-regulated in dementia

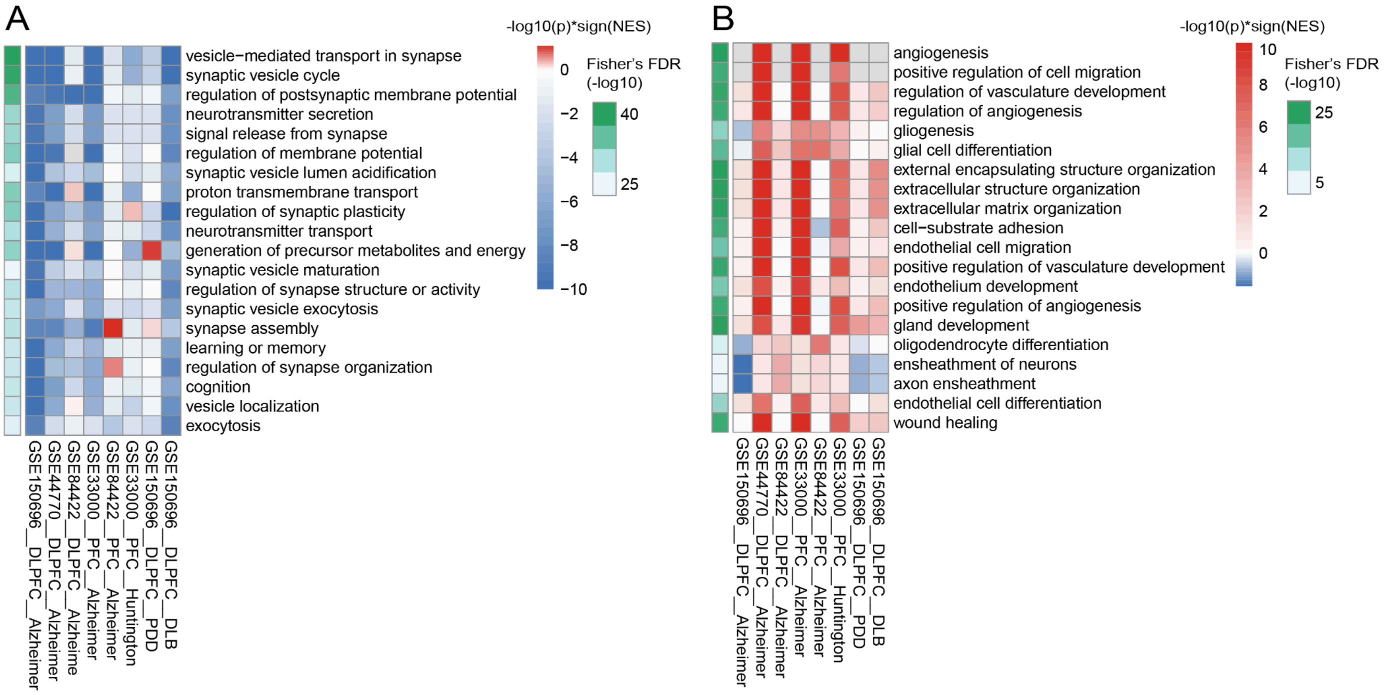


**Figure S9.** Top 20 Gene Ontology (GO) categories statistically enriched in the genes up-regulated by LSD and down-regulated in dementia (A), and in the genes down-regulated by LSD and up-regulated in dementia (B). The colour in the heatmap indicates the GSEA p-value (-log10) with sign depending on the sign of the normalized enrichment score (NES). Red, highly significant positive enrichments; blue, highly significant negative enrichments. The green bar on the left indicates the overall FDR (-log10) combining the p-values obtained for each dataset.

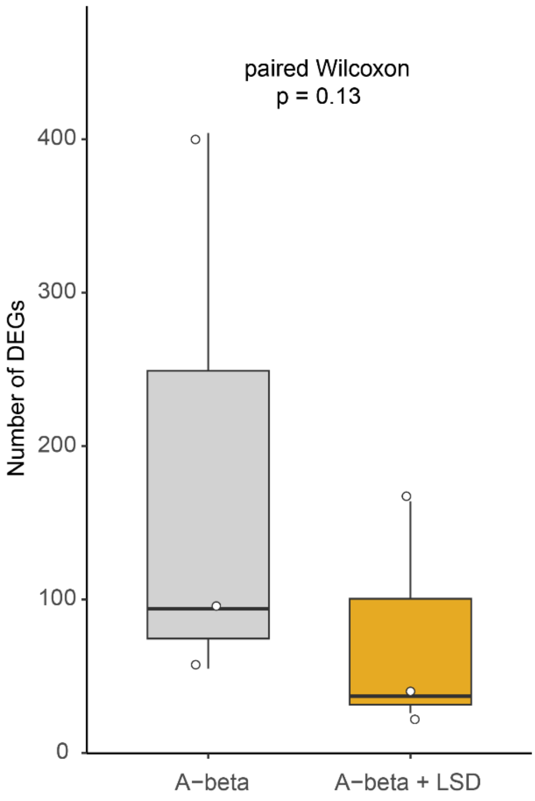


**Figure S10.** Number of significantly differentially expressed genes (DEGs; FDR < 0.05) relative to control for Aβ alone and for the corresponding combined Aβ+LSD treatment across three Aβ concentrations (100 nM, 500 nM, 1 µM). The p-value was computed using a paired Wilcoxon signed-rank test (one-sided), pairing each Aβ dose with its matched Aβ+LSD condition.

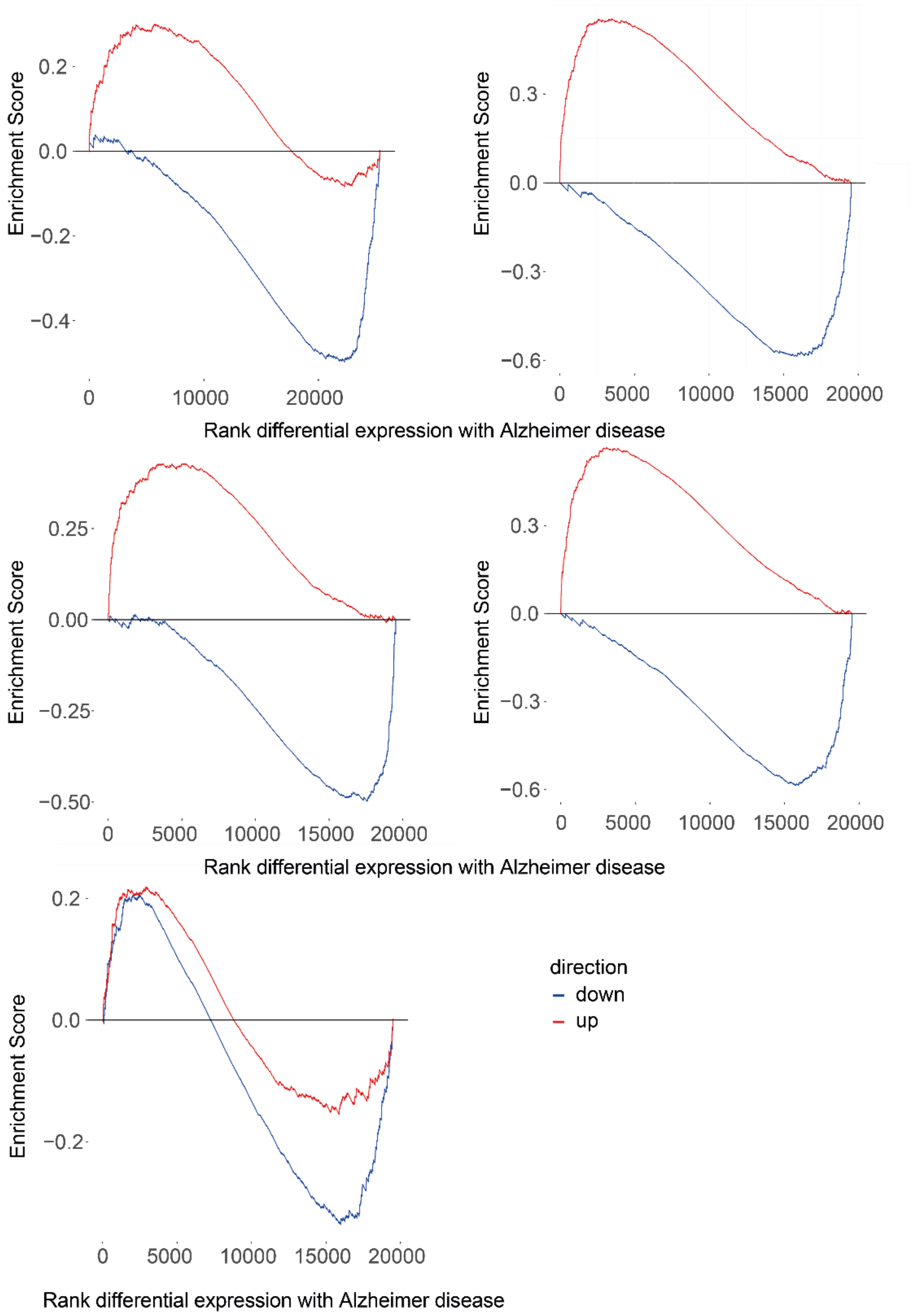


**Figure S11.** Gene set enrichment analysis (GSEA) was performed independently for five human human brain datasets comparing Alzheimer's disease (AD) versus healthy control samples. Genes were ranked by log2 fold change differential expression analysis (AD vs control). Enrichment curves show the running enrichment score for genes upregulated (red) or downregulated (blue) by Aβ treatment in vitro, after mapping to human orthologs. Positive enrichment indicates concordance between the in vitro Aβ signature and transcriptional changes observed in AD brain tissue, whereas negative enrichment indicates inverse regulation.

**Table S17.** Full list of Gene Ontology categories tested for differential expression in primary neurons treated with LSD plus Aβ relative to Aβ alone.

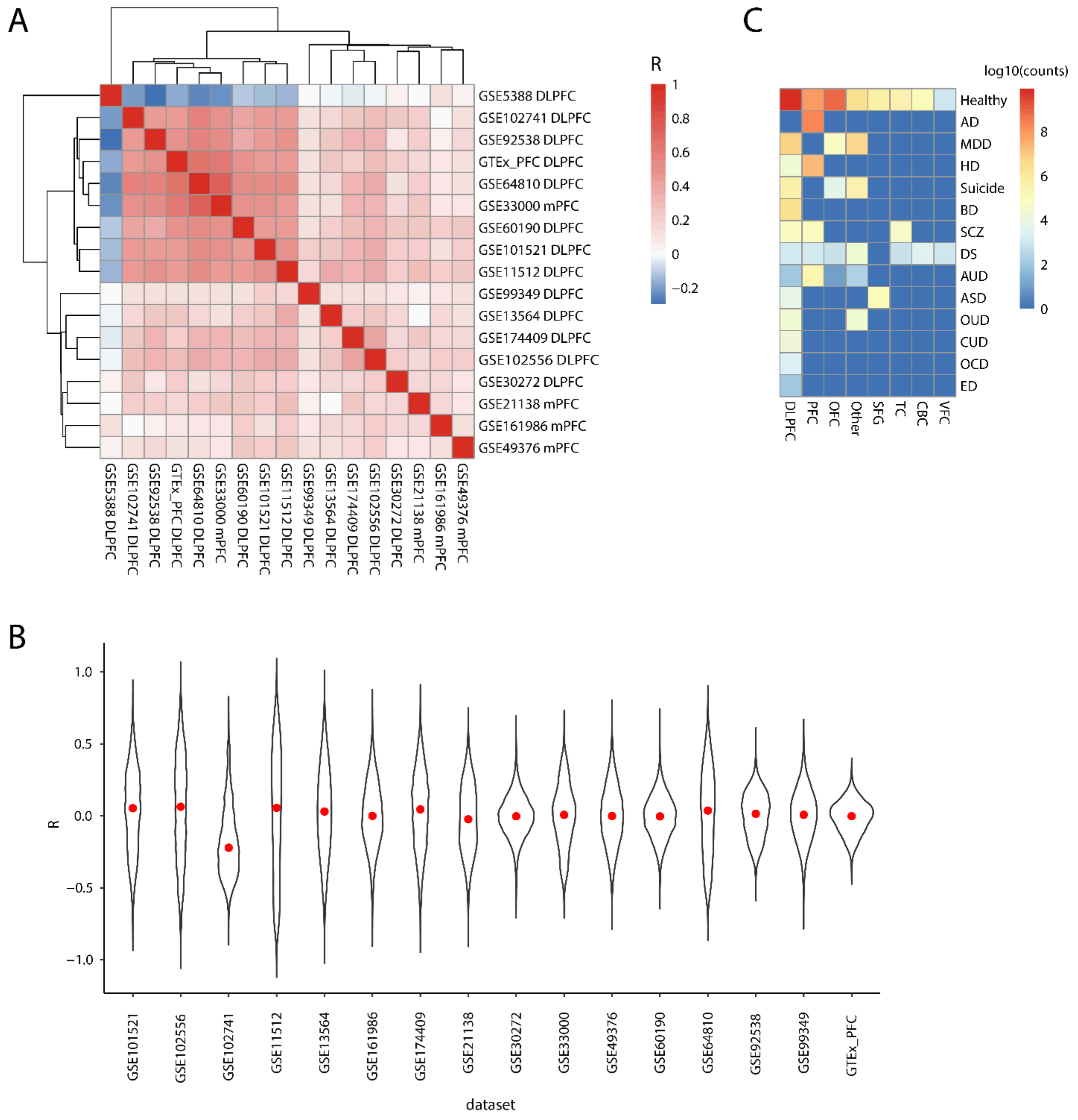


**Figure S12.** A) Similarity (Pearson's correlation) between genes' correlation with age in different ageing datasets. B) Distribution of genes' correlation with age in different ageing datasets (median in red). C) Number of samples in the ageing datasets collection belonging to each combination of categories. DLPFC = dorsolateral prefrontal cortex; mPFC = medial prefrontal cortex; OFC = orbitofrontal cortex; SFG = superior frontal gyrus; TC= temporal cortex, CBC = cerebellar cortex; VFC = ventral frontal cortex; AD = Alzheimer disease; MDD = major depressive disorder; HD = Huntington disease; BD = bipolar disorder; SCZ = schizophrenia; DS =Down syndrome; AUD = alcohol use disorder; ASD = autism spectrum disorder; OUD = opioid use disorder; CUD = cannabis use disorder; OCD = obsessive compulsive disorder; ED = eating disorder.

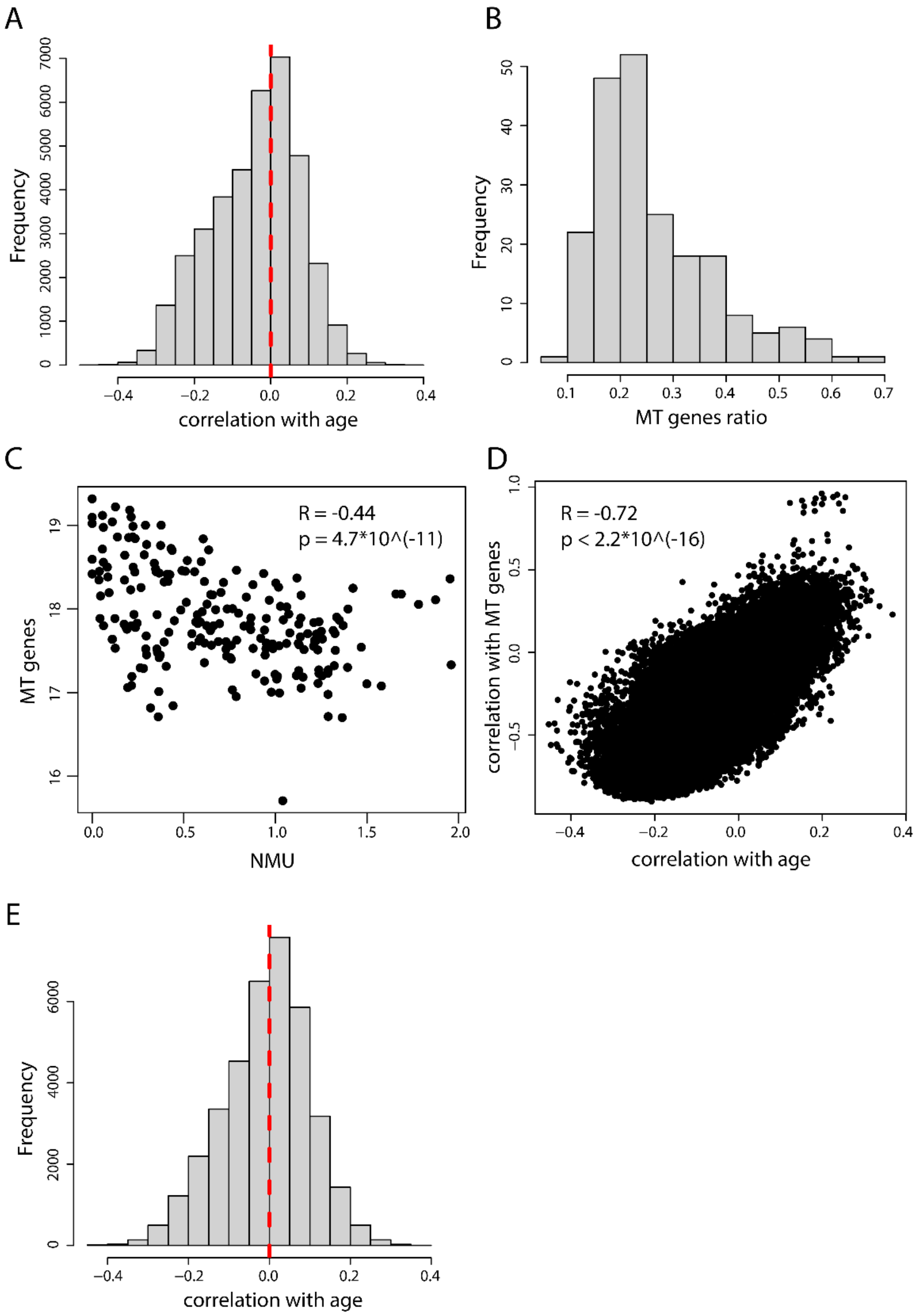


**Figure S13.** Correcting a normalization bias introduced by mitochondrial genes in the GTEx dataset. A) Distribution of gene expression - age correlations across GTEx frontal cortex samples; zero is indicated by a dashed red line. B) fraction of reads mapping to mitochondrial genes over the total number of reads mapping to genes. C) Expression of NMU, the gene with the lowest correlation with age, vs mitochondrial genes across GTEx frontal cortex samples. D) Relationship between a gene's correlation with age and correlation with the expression of mitochondrial genes. E) Distribution of gene expression - age correlations across GTEx frontal cortex samples after removing the top 10% of samples with the highest expression of mitochondrial genes; zero is indicated by a dashed red line.

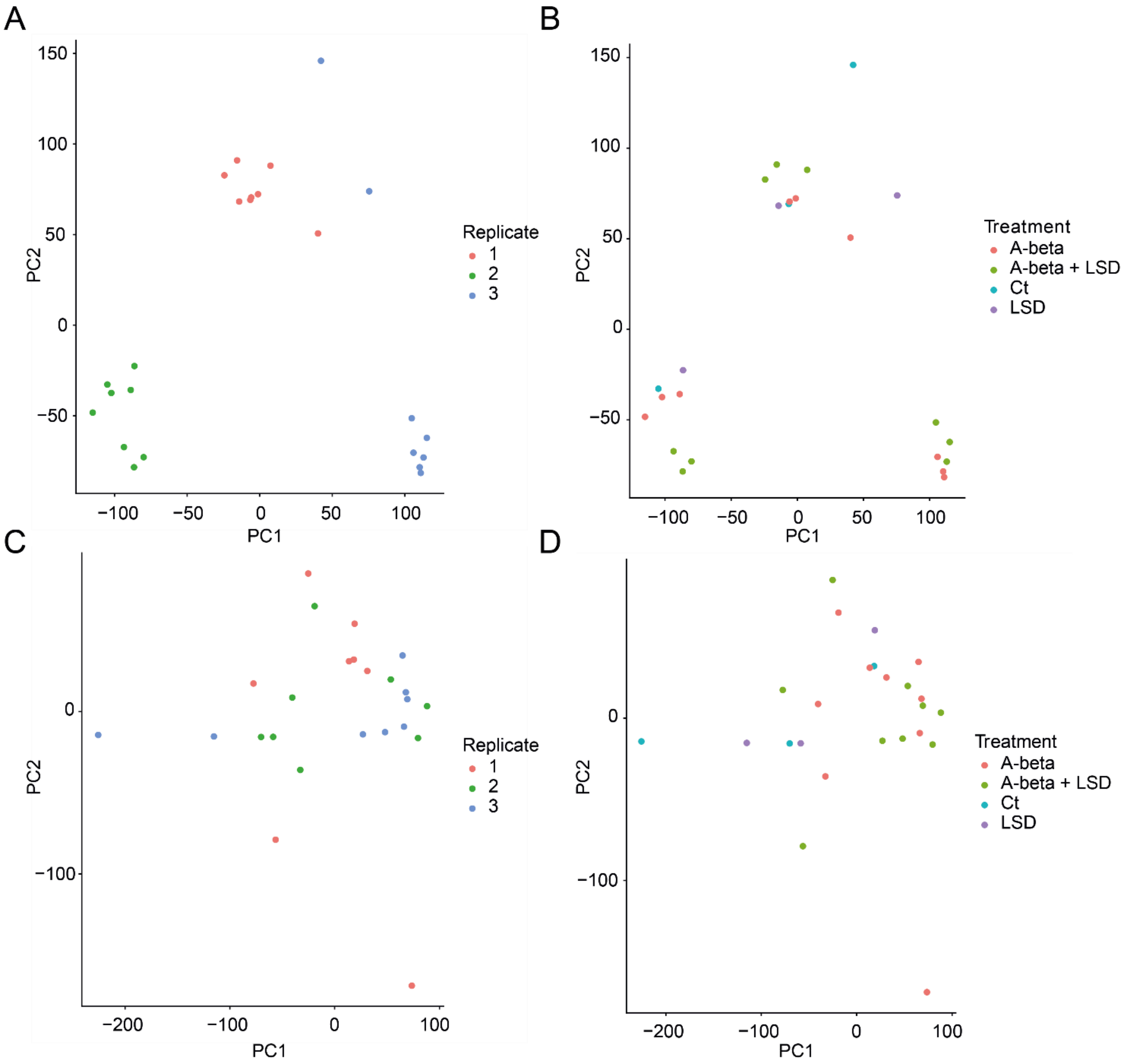


**Figure S14.** Principal component analysis of in vitro RNA-seq samples based on gene expression pre (AB) and post (CD) correction for batch. Each dot is a sample. A,B) are coloured according to replicate, C,D) are coloured according to treatment.

### Supplementary Methods

***Aging datasets collection and curation***

We mined the Gene Expression Omnibus (GEO) database (Clough & Barrett 2016) for transcriptomic data of human prefrontal cortex (PFC) samples with available age annotation of the donors as follows. On the 7th of April 2022, we used the query (prefrontal cortex[All Fields] AND age[All Fields]) AND "Homo sapiens"[porgn] AND ("gse"[Filter] AND ("Expression profiling by array"[Filter] OR "Expression profiling by high throughput sequencing"[Filter])), retrieving 95 raw GEO datasets with at least 10 samples (**Supplementary File S1**), which we downloaded using the GEOquery R package (Davis & Meltzer 2007). We processed Affymetrix arrays with the Robust Multichip Average (RMA) normalization (performing background correction and probe-level quantile normalization) and Illumina arrays with quantile normalization using negative and positive control probes (Stafford 2008). We mapped microarray probes to gene symbols using the respective platforms' information in GEO for each dataset, and when multiple probes matched the same gene symbol, we selected the probe with the highest mean expression across the considered dataset's samples. When data were not already log-transformed, we applied a $\log_2$ transformation, adding an offset of 1.

We thoroughly manually inspected each dataset and kept only data allowing computing correlation scores between sample age and individual genes' expression level across samples, removing samples for which age annotation was not reported, then filtering out datasets with less than 10 remaining samples per analysed brain region, or not including human samples or derived from in vitro models only (**Table S1**). We thus retained 28 datasets (**Table S2**) and included an external dataset from GTEx (GTEx Consortium 2013) comprising samples from 188 subjects transcriptionally profiled in the dorsolateral prefrontal cortex (DLPFC) (after quality filtering, see below).

We observed that several datasets included samples from very young subjects, thus being most useful for the study of brain development (**Figure S1**). Given our focus on aging, we kept datasets with at least 10 samples derived from adult subjects (>= 20 years). This left us with 24 datasets for a total of 3,086 samples spanning a wide range of ages (from 2 days to 106 years) and including also pre-natal samples. Despite our interest in the prefrontal cortex (PFC) of healthy adult subjects, we included in our collection all samples from the selected datasets, and extensively manually curated and harmonized the metadata, comprising annotation for gender, race, psychiatric diagnosis, and the dissected brain regions (Table S3, Figure S1BC). Nevertheless, in our comparisons between aging and LSD transcriptional signatures, we only employed datasets annotated as DLPFC or PFC.

Quality filtering assessments led us to discard an outlier dataset showing a low similarity in gene-age relationships with the remaining 16 DLPFC and PFC datasets (GSE5388, **Figure S12A**) and a dataset with skewed correlation between gene expression and age (GSE102741, **Figure S12B**).

We downloaded GTEx data alongside sample age annotations from the GTEx Portal(GTEx Consortium 2013) on 20/09/2022 (files: 'gene_reads_2017-06-05_v8_brain_frontal_cortex_ba9.gct.gz' and 'https://storage.googleapis.com/gtex_analysis_v8/annotations/GTEx_Analysis_v8_Annotations_SubjectPhenotypesDS.txt' - currently available at https://storage.googleapis.com/adult-gtex/annotations/v8/metadata-files/GTEx_Analysis_v8_Annotations_SubjectPhenotypesDS.txt); we normalised RPM (reads per million), mapped gene identifiers to gene symbols as described above for other datasets, and log-transformed resulting data (with an offset of 1).

We removed the 10% of samples with the highest expression of mitochondrial genes (identified based on the naming “MT-“).

Before this correction, genes’ correlation with age was skewed toward negative values (**Figure S13A**; 1st quartile:-0.12 , median: -0.03, 3rd quartile: 0.04). We observed that mitochondrial genes’ reads represented a high proportion of reads in some samples, accounting for 65% of reads mapped to genes (**Figure S13B**) and we therefore reasoned that this could generate normalisation artefacts in these samples, potentially leading to spurious gene expression-age correlations. Indeed, genes that are negatively correlated with age are expressed at lower levels in samples with high levels of MT genes (**Figure S13CD**). Removing the top 10% of samples with the highest expression of MT genes, correlations’ skewing was reduced (**Figure S13E**; 1st quartile: -0.08, median: 0.0008, 3rd quartile: 0.06). Finally, we retained genes with RPM>0 in at least 10 samples.

***Comparing the age-signature-reversal/-mimicking potential of psychoplastogen with that of positive and negative controls***

We quantified the age-signature reversal and mimicking potential of each intervention by testing enrichment of treatment-induced gene sets along aging-associated ranked lists using a directional GSEA framework, followed by aggregation across datasets and comparison of reversal versus mimicking signals, as detailed in the Supplementary Methods.

For each intervention (psychoplastogens, enriched environment, exercise, ethanol, or rat aging signatures), we identified differentially expressed genes (DEGs; FDR < 0.05), separating up- and down-regulated genes. These gene sets were used independently as query signatures in GSEA against each aging-ranked list. GSEA yielded an enrichment score (ES) and an associated two-sided p-value for each query and dataset.

Under the age-reversal hypothesis, treatment-induced up-regulated genes are expected to be enriched among genes whose expression decreases with age (i.e. at the bottom of the ranked list, corresponding to negative ES). In contrast, treatment-induced down-regulated genes are expected to be enriched among genes whose expression increases with age (i.e. at the top of the ranked list, corresponding to positive ES). Conversely, under an age-mimicking hypothesis, enrichment is expected in the opposite directions.

Because our hypotheses are directional, we converted the two-sided GSEA p-values into one-sided p-values using the two2one function from the metap R package. To incorporate directionality, we adjusted p-values based on the sign of the ES. Specifically, when the observed ES was inconsistent with the tested hypothesis (e.g. positive ES for up-regulated genes under the age-reversal framework), the corresponding one-sided p-value was inverted (1 − p), thereby encoding direction into the statistical measure. For example, a two-sided p-value of 0.05 becomes 0.025 under directional consistency, but 0.975 if the ES sign contradicts the reversal hypothesis. This transformation ensures that small p-values consistently represent evidence in favor of the tested direction.

For each intervention and aging dataset, this procedure yielded two directional p-values (one for up-regulated genes and one for down-regulated genes). We then aggregated p-values across aging datasets using Fisher's method to obtain overall measures of evidence for age-reversal, separately for up- and down-regulated genes.

To assess whether observed signals reflected true age-reversal rather than generic concordance with aging trajectories, we repeated the same procedure under the age-mimicking framework by inverting the directional expectations (i.e. switching the ES sign considered consistent with the hypothesis). This yielded aggregated age-mimicking p-values, again separately for up- and down-regulated genes.

Thus, for each intervention, we obtained four aggregated p-values: age-reversal and age-mimicking statistics for up- and down-regulated gene sets. Finally, to quantify the relative strength of reversal versus mimicking effects, we computed the ratio between aggregated age-reversal and age-mimicking p-values. This ratio provides a directional reliability measure, where values favoring reversal indicate stronger support for opposition to aging-associated transcriptional programs.

##### *Testing psychoplastogens potential in reverting signatures of dementia*

We searched the Gene Expression Omnibus for transcriptomic data of human prefrontal cortex (PFC) in dementia on the 1/4/2022. Our query was: (("dementia"[MeSH Terms] OR dementia[All Fields]) AND prefrontal cortex[All Fields]) AND "Homo sapiens"[porgn] AND ("gse"[Filter] AND ("Expression profiling by array"[Filter] OR "Expression profiling by high throughput sequencing"[Filter])).

The query yielded 20 datasets from which we retained bulk transcriptomic data of DLPFC finally considering 6 datasets (**Table S8**). On these, we tested the relationship between LSD-induced and dementia-related transcriptional changes.

For each dataset, we selected healthy and demented samples of the DLPFC or mPFC. We performed a single sample GSEA (using the GSVA R package (Hänzelmann et al. 2013)) for each selected sample and each considered list of genes, i.e. up-regulated and down-regulated upon treatment with psychoplastogens/ethanol/enriched environment/exercise (FDR<0.05), employing the datasets described in “Psychoplastogen’s datasets” and “Positive and negative controls” sections of the methods. For each treatment, we then compared the single sample GSEA scores resulting from probing healthy and demented samples with a t-test and computed the Cohen’s d to measure the effect size of the difference.

We then proceeded similarly as described in the “testing age reversal” section of the Methods. Specifically, we merged the GSEA p-values using the Fisher method (Edwards 2005) (using the metap R package), thus computing global reliability scores for dementia-reversal and dementia-inducing effects of each intervention (still keeping up- and down-regulated genes upon treatment separated). Finally, we quantified the reliability of dementia-reversal activity for each treatment in comparison to the reliability of dementia-mimicking ability of the same treatments. To this aim, we repeated the procedure testing the dementia mimicking ability of each treatment, switching the Cohen’s d sign. So, for each treatment, we obtained an overall dementia-reversal and dementia-mimicking p-value based on up- or down-regulated genes. We then computed the ratio of the dementia-reversal and dementia-mimicking p-values, obtaining the results shown in **Figure 1E**.

##### *Functional enrichment (human datasets)*

We tested the functional enrichment of genes correlated with aging, analysing one aging dataset at a time through a gene set enrichment analysis (GSEA) using as a background list the genes ranked based on the correlation between basal expression with age, and gene ontology (GO) Biological Processes as gene signature collection. Similarly, for dementia, within each dataset, we computed a gene-wise ranking statistic reflecting differential expression between the case and healthy groups. Specifically, for each gene we computed a t-statistic for the difference in means (case - healthy) using group-specific variances and sample sizes. For each ranked gene list, we performed GO Biological Process gene set enrichment. We merged the p-values of individual datasets (either for aging or for dementia) with the Fisher method and retained GO categories with enrichment FDR < 0.05, correcting p-values for multiple testing with the Benjamini-Hochberg correction (p.adjust function from the stats R package).

##### *Functional enrichment (in vivo LSD treatment)*

We considered genes differentially expressed upon LSD (at an FDR < 0.05) and performed an enrichment analysis for GO biological processes, separately for each LSD dataset (chronic treatment and acute treatment with 50, 100, 200, 500 ug/kg LSD). We retained only the GO categories significantly enriched in genes up-regulated across LSD datasets (and, conversely, enriched in genes down-regulated across LSD datasets), merging the p-values of individual datasets with the Fisher method and finally retaining GO categories with FDR < 0.05.

Finally, we combined the information on GO categories enriched for genes correlated with aging and for genes differentially expressed upon LSD and presented only the categories enriched for genes with an opposite trend upon aging and LSD (Figure 2). Similarly, we combined the GO categories enriched for genes changing in opposite directions with LSD and dementia, and showed them in **Figure S9**.

*Functional enrichment (in vitro treatments)*

To assess whether the Gene Ontology (GO) categories showing opposite-direction regulation between LSD and human dementia were also modulated by LSD in an in vitro model of neurodegenerative stress, we performed gene set analysis on RNA-seq data from primary mouse cortical neurons treated with Aβ alone or with Aβ followed by LSD.

GO categories tested in this analysis were not selected a priori, but were derived directly from the cross-dataset LSD-dementia comparison, focusing on biological process terms that were significantly regulated in opposite directions by LSD in vivo and across human dementia transcriptomes (FDR $< 0.05$). For each selected GO term, the corresponding mouse orthologous genes were retrieved using the org.Mm.eg.db annotation. Only gene sets containing at least 10 expressed genes in the in vitro dataset were retained for downstream analysis.

Competitive gene set testing was performed using CAMERA (Correlation Adjusted MEan RAnk gene set test) as implemented in the limma package. CAMERA evaluates whether genes belonging to a predefined gene set are, on average, more differentially expressed than genes outside the set, while explicitly accounting for inter-gene correlation within each gene set, thereby reducing false positives associated with coordinated gene expression.

CAMERA was applied to log-transformed, ComBat-seq–adjusted gene expression values using a linear model contrasting Aβ + LSD versus Aβ treatment. Results are reported as direction of regulation, p-values, and Benjamini–Hochberg–adjusted false discovery rates (FDR). This analysis enabled us to directly test whether GO programs identified as LSD-reverted in human dementia were concordantly modulated by LSD in neurons exposed to Aβ-induced stress.
